# Metagenomic assembly of new (sub)arctic Cyanobacteria and their associated microbiome from non-axenic cultures

**DOI:** 10.1101/287730

**Authors:** Luc Cornet, Amandine R. Bertrand, Marc Hanikenne, Emmanuelle J. Javaux, Annick Wilmotte, Denis Baurain

## Abstract

Cyanobacteria form one of the most diversified phylum of Bacteria. They are important ecologically as primary producers, for Earth evolution and biotechnological applications. Yet, Cyanobacteria are notably difficult to purify and grow axenically, and most strains in culture collections contain heterotrophic bacteria that were likely associated to Cyanobacteria in the environment. Obtaining cyanobacterial DNA without contaminant sequences is thus a challenging and time-consuming task. Here, we deploy a metagenomic pipeline that enables the easy recovery of high-quality genomes from non-axenic cultures. We tested this pipeline on 17 cyanobacterial cultures from the BCCM/ULC public collection and generated novel genome sequences for 15 arctic or subarctic strains, of which 14 early-branching organisms that will be useful for cyanobacterial phylogenomics. In parallel, we managed to assemble 31 co-cultivated bacteria from the same cultures and showed that they mostly belong to Bacteroidetes and Proteobacteria, some of them being very closely related in spite of geographically distant sampling sites.

**Importance:** Complete genomes of cold-adapted Cyanobacteria are underrepresented in databases, due to the difficulty to grow them axenically. In this work, we report the genome sequencing of 12 (sub)arctic and 3 temperate Cyanobacteria, along with 21 Proteobacteria and 5 Bacteroidetes recovered from their microbiome. Following the use of a state-of-the-art metagenomic pipeline, 12 of our new cyanobacterial genome assemblies are of high-quality, which indicates that even non-axenic cultures can yield complete genomes suitable for phylogenomics and comparative genomics. From a methodological point of view, we investigate the fate of SSU rRNA (16S) genes during metagenomic binning and observe that multi-copy rRNA operons are lost because of higher sequencing coverage and divergent tetranucleotide frequencies. Moreover, we devised a measure of genomic identity to compare metagenomic bins of different completeness, which allowed us to show that Cyanobacteria-associated bacteria can be highly related in spite of considerable distance between collection points.

## Introduction

Cyanobacteria, also called blue-green algae, are an intensively studied group of prokaryotes. This focus is notably due to their ecological importance, as they colonize a very diverse range of ecosystems and are a major component of the phytoplankton (1, 2). They are also of primary interest in terms of evolution and palaeobiogeology, Cyanobacteria having been present on Earth since the Proterozoic (3–5). Emergence of oxygenic photosynthesis in this phylum, which led to the Great Oxygenation Event (GOE) around 2.4 billion years ago, had a critical impact on early Earth and evolution by increasing the level of free oxygen and subsequently creating new ecological niches (6–8). Moreover, Cyanobacteria played a role in another major biological event, the spread of photosynthesis to eukaryotic lineages through an initial endosymbiosis termed “primary”, followed by several higher-order endosymbioses (9). Finally, Cyanobacteria produce a large number of bioactive compounds (e.g., alkaloids, non-ribosomal peptides, polyketides), which make them promising for both biotechnological and biomedical applications (10–12). Cyanobacteria are notoriously difficult to isolate and keep axenic in culture (1), especially polar strains (13), hence the need for tedious purification protocols (14). In consequence, all cyanobacterial culture collections include a majority of non-axenic cultures (e.g., Czech Collection of Algae and Cyanobacteria, CCALA; University of Toronto Culture Collection of Algae and Cyanobacteria, UTCC), with the notable exception of the Pasteur Culture Collection of Cyanobacteria, PCC. The difficulty of reaching axenicity results from bacterial communities living in close relationship with Cyanobacteria in nature. This microbiome has been described both from environmental samples (15–19) and non-axenic cultures (20–22). Moreover, Bacteria/Cyanobacteria associations appear to be stable in culture, as no significant differences could be found between bacterial communities accompanying Cyanobacteria in fresh samples and collection cultures (21). Complex trophic interactions between Cyanobacteria and other bacterial phyla feeding on their sheaths, such as Proteobacteria and Bacteroidetes, have been described (23), as well as specific interactions, such as adhesion to heterocysts (20). The presence of these bacterial communities consequently limits the use of non-axenic cyanobacterial cultures for genomic applications, because fragments of their genomes can eventually become part of published cyanobacterial genomes. Hence, we have recently shown that a large proportion (52%) of publicly available genomes of Cyanobacteria are contaminated by such foreign sequences (Cornet et al., 2018, in revision). In 5% of the surveyed genomes, these non-cyanobacterial contaminants even reach up to 41.5% of the genome sequences deposited in the databases.

Owing to their clear scientific interest, obtaining authentic genome sequences of Cyanobacteria is an important issue. During the last decade, the rise of metagenomics has allowed an ever-better separation of the different components of a mixture of organisms, based on various properties of the metagenomic contigs, e.g., sequencing coverage and oligonucleotide signatures (24). In this work, we use a straightforward pipeline that enables the efficient isolation of cyanobacterial genomes from non-axenic cultures. Easy to deploy, this pipeline is composed of state-of-the-art metagenomic tools, metaSPAdes (25), MetaBAT (26), CheckM (27), followed by DIAMOND blastx analyses (28) and SSPACE (29) scaffolding. This pipeline allowed us to assemble 15 novel cyanobacterial genomes (12 high-quality, 2 medium-quality and 1 low-quality) from 17 arctic and subarctic cultures of the BCCM/ULC public culture collection hosted by the University of Liège (ULiège, Belgium), of which 14 appear to belong to early-branching strains in the cyanobacterial tree of life. In the process, we also characterized 31 different co-cultivated bacteria out of the 17 cyanobacterial cultures. Those “contaminant” organisms mostly belong to Proteobacteria and Bacteroidetes, and some of them are very closely related to each other. Finally, we investigated why SSU rRNA (16S) genes are often lost during metagenomic binning and developed a new metric to compare genome bins with different levels of completeness.

## Materials and Methods

### Cyanobacterial cultures and DNA extraction

The 17 cyanobacterial cultures were selected in order to sequence new genomes of interesting Arctic and Antarctic organisms, from which the biodiversity is still not well known. All the strains used in this study were indeed collected from (sub)arctic regions, at the exception of three Belgian strains, ULC335, added to the sequencing batch to obtain the first genome of the genus *Snowella*, and ULC186 and ULC187, both related to the (sub)polar strains but of temperate origin. The cultures (deposited in the BCCM/ULC collection during the period 2011–2014; **Table 1**) were incubated at 15°C and exposed to a constant white fluorescent light source (about 40 μmol photons m^-2^ s^-1^) for 4 weeks. The DNA was extracted using the GenElute Bacterial Genomic DNA kit (Sigma-Aldrich, Saint-Louis, Mo, USA) following the recommendations of the manufacturer. After control of the integrity of the genomic DNA by electrophoresis and quantification of the dsDNA concentration using the Quan-iT Picogreen dsDNA Assay kit (Thermo Fisher Scientific, Waltham, Mass, USA), a minimum of 1 μg of dsDNA was sent to the sequencing platform.

**Table 1:** Details about ULC strains. All details were extracted from the BCCM/ULC website: http://bccm.belspo.be/about-us/bccm-ulc.

| Assembly | Strain | Name | Type | Prior affiliation | Morphology | Sheath | Deposit date | Habitat |
| --- | --- | --- | --- | --- | --- | --- | --- | --- |
| In process | ULC187 | *Pseudanabaena* sp FW039 | non-axenic | Clade F | Filamentous | NO | 2012 | Belgium, lake Ri Jaune |
| In process | ULC066 | *Pseudanabaena* | non- | Clade F | Filamentous | NO | 2011 | Canadian |
|  |  | *frigida* O-155 | axenic |  |  |  |  | Arctic, Bylot Island |
| In process | ULC068 | *Pseudanabaena* sp. O-202 | non-axenic | Clade F | Filamentous | NO | 2011 | Canadian Subarctic, Québec, Kuujjuarapik |
| In process | ULC065 | *Cyanobium* sp. O-154 | non-axenic | Clade C1 | Unicellular | NO | 2011 | Canadian Arctic, Bylot Island |
| In process | ULC082 | *Cyanobium* sp. Chester Cone | non-axenic | Clade C1 | Unicellular | NO | 2011 | Antarctica, Livingston Island |
| In process | ULC084 | *Cyanobium* sp. Laguna Chica | non-axenic | Clade C1 | Unicellular | NO | 2011 | Antarctica, Livingston Island |
| In process | ULC077 | *Leptolyngbya* sp. O-157 | non-axenic | Clade C3 | Filamentous | NO | 2011 | Canadian Arctic, Bylot Island |
| In process | ULC007 | *Phormidesmis priestleyi* ANT.LH52.4 | axenic | Clade C3 | Filamentous | NO | 2011 | Antarctica, Larsemann Hills |
| NA | ULC165 | *Leptolyngbya* sp. OTC1/1 | non-axenic | Clade C3 | Filamentous | YES | 2012 | Antarctica, Sor Rondane Mountains |
| In process | ULC129 | *Leptolyngbya foveolarum* TM2FOS129 | non-axenic | Clade C3 | Filamentous | NO | 2011 | Antarctica, Transantarctic Mountains |
| In process | ULC027 | *Phormidium priestleyi* ANT.PROGRESS 2.5 | non-axenic | Clade C3 | Filamentous | NO | 2011 | Antarctica, Larsemann Hills |
| In process | ULC186 | *Leptolyngbya* sp. FW074 | non-axenic | Clade C3 | Filamentous | NO | 2012 | Belgium, Renipont lake |
| In process | ULC041 | *Leptolyngbya antarctica* ANT.ACE.1 | non-axenic | Clade C3 | Filamentous | NO | 2011 | Antarctica, Vestfold Hills |
| In process | ULC073 | *Leptolyngbya glacialis* TM1FOS73 | non-axenic | Clade C3 | Filamentous | YES | 2011 | Antarctica, Transantarctic Mountains |
| In process | ULC335 | *Snowella* sp. FW024 | non-axenic | Clade B2 | Unicellular | YES | 2014 | Belgium, lake Falemprie |
| NA | ULC146 | *Nostoc* sp. ANT.UTS.183 | non-axenic | Clade B1 | Filamentous heterocystous | YES | 2012 | Antarctica, Sor Rondane Mountains |
| NA | ULC179 | *Nostoc* sp. OTCcontrol | non-axenic | Clade B2 | Filamentous heterocystous | YES | 2012 | Antarctica, Sor Rondane Mountains |

### Metagenome sequencing and assembly

The 17 cyanobacterial cultures were sequenced (PE 2x 250 nt) on the Illumina MiSeq sequencing platform (GIGA Genomics, ULiège). Nextera XT libraries had a fragment size estimated at 800-900 nt. Raw sequencing reads were trimmed using Trimmomatic v0.35 (30). Sequencing adapters were removed with the option illuminaclip NexteraPE-PE.fa:2:30:20. Trimming values were selected to maximize genome bin sizes (in terms of bp), after preliminary testing. Trailing/leading values were set at 20, the sliding window at 10:20, the crop value at 145 and the minimal length at 80. Trimmed paired-end reads were assembled with metaSPAdes v3.10.1 (25) using default settings. Trimmed paired-end reads were then re-mapped on the metaSPAdes assemblies with BamM v1.7.3 (http://ecogenomics.github.io/BamM/), yielding BAM files suitable for the metagenomic analyses. Genome bins were determined with MetaBAT v0.30.1 (26), trying each built-in parameter set in turn (i.e., verysensitive, sensitive, specific, veryspecific, and superspecific). CheckM v1.0.7 (27) was then used with the option lineage_wf to select the best MetaBAT parameter set for each metaSPAdes assembly. In practice, we first tried to select the MetaBAT parameter set that was the most suitable for the largest genome bin of a given metagenome (in terms of total assembly length), considering CheckM output statistics in the following order: 1) contamination, 2) strain heterogeneity, 3) completeness. When multiple parameter sets were equally optimal for the largest bin, we turned to the next-largest bin(s) for parameter selection. The non-assignment of a given contig to multiple bins was checked using the unique option of CheckM, while binning accuracy was assessed using merge and tree_qa options after generating a marker set for Bacteria. The automatic taxonomic classification of CheckM was then extracted to determine the nature of each bin, either cyanobacterial or foreign. Bins classified as root (i.e., unclassified) by CheckM were discarded from phylogenomic analyses. Contaminants (with respect to the taxon determined by CheckM) in each genome bin were further characterized using DIAMOND blastx v0.8.22 (28) and the companion parser developed in our article about the contamination of public cyanobacterial genomes (Cornet et al., 2018, in revision). To this end, we split the genome bins into non-overlapping pseudo-reads of 250 nt (with a custom Perl script), so as to increase the sensitivity of the analyses. We then used DIAMOND blastx to blast these pseudo-reads against a curated database derived from the release 30 of Ensembl Bacteria. In parallel, contigs within each genome bin were scaffolded with SSPACE v.3.0 (29) using default settings, except that contigs were first extended using paired-end reads (-x 1) and that the minimum of read pairs required to compute a scaffold was set to 3 (-k 3). The fragmentation of the scaffolded genome bins was then analyzed with QUAST v2.3 (31) using default settings, whereas their sequencing coverage was determined with BBMap v37.24 (http://bbmap.sourceforge.net/). Finally, protein sequences were predicted for all genome bins with Prodigal v2.6.2 (32) using the ab_initio mode.

### Phylogenetic analyses

The complete proteomes of 64 cyanobacterial strains chosen to represent the diversity of the whole phylum were downloaded from the NCBI portal (33). Details and download links for the selected proteomes are available in **Tables 2** and **S1**, respectively. Orthology inference was performed with USEARCH v8.1 (64 bits) (34) and OrthoFinder v1.1.2, using the standard inflation parameter of 1.5 (35). Out of 37,261 orthologous groups (OGs), 675 were selected with classify-ali.pl (part of the Bio-MUST-Core software package; D. Baurain; https://metacpan.org/release/Bio-MUST-Core) by enforcing in each OG the presence of ≥62 different organisms, represented by an average of ≤1.1 gene copy per organism. The 675 OGs were enriched with sequences directly mined from the 15 cyanobacterial bins using the software “42” (Baurain et al., to be published elsewhere; https://bitbucket.org/dbaurain/42/), which strictly controls for orthology during enrichment. Enriched OGs were then aligned with MAFFT v7.273 (36) and conserved sites were selected with BMGE v1.12 (37) using moderately severe settings (entropy cut-off 0.5, gap cut-off 0.2). A supermatrix of 79 organisms x 170,983 unambiguously aligned amino-acid positions (3.9% missing character states) was assembled with SCaFoS v1.30k (38) using the minimal evolutionary distance criterion for deciding between the few in-paralogous proteins. Finally, a phylogenomic tree was inferred with PhyloBayes-MPI v1.5a under the CAT+Γ_4_ model (39) by running two independent chains until 1500 cycles were obtained. Convergence of the parameters was assessed using criteria given in the PhyloBayes manual and a conservative burn-in of 620 cycles was used (meandiff = 0.04).

**Table 2:** Details about reference proteomes. All details were extracted from the NCBI metadata.

| Assembly | Bioproject | Taxid | Name |
| --- | --- | --- | --- |
| GCA_000484535.1 | PRJNA162637 | 1183438 | *Gloeobacter kilauensis* JS1 |
| GCF_000011385.1 | PRJNA58011 | 251221 | *Gloeobacter violaceus* PCC 7421 |
| GCF_000013205.1 | PRJNA224116 | 321327 | *Synechococcus* sp. JA-3-3Ab |
| GCF_000013225.1 | PRJNA224116 | 321332 | *Synechococcus* sp. JA-2-3B'a(2-13) |
| GCF_000332275.1 | PRJNA224116 | 195250 | *Synechococcus* sp. PCC 7336 |
| GCF_000317065.1 | PRJNA224116 | 82654 | *Pseudanabaena* sp. PCC 7367 |
| GCF_000332215.1 | PRJNA224116 | 927668 | *Pseudanabaena biceps* PCC 7429 |
| GCF_000317085.1 | PRJNA224116 | 1173263 | *Synechococcus* sp. PCC 7502 |
| GCF_000332175.1 | PRJNA224116 | 118173 | *Pseudanabaena* sp. PCC 6802 |
| GCF_000018105.1 | PRJNA224116 | 329726 | *Acaryochloris marina* MBIC11017 |
| GCA_000022045.1 | PRJNA28337 | 395961 | *Cyanothece* sp. PCC 7425 |
| GCF_000505665.1 | PRJNA224116 | 1394889 | *Thermosynechococcus* sp. NK55a |
| GCF_000316685.1 | PRJNA224116 | 195253 | *Synechococcus* sp. PCC 6312 |
| GCF_000775285.1 | PRJNA224116 | 1497020 | *Neosynechococcus sphagnicola* sy1 |
| GCF_000309945.1 | PRJNA224116 | 864702 | *Oscillatoriales cyanobacterium* JSC-12 |
| GCF_001895925.1 | PRJNA224116 | 1920490 | *Phormidesmis priestleyi* ULC007 |
| GCF_001650195.1 | PRJNA224116 | 1850361 | *Phormidesmis priestleyi* BC1401 |
| GCF_000353285.1 | PRJNA224116 | 272134 | *Leptolyngbya boryana* PCC 6306 |
| GCF_000733415.1 | PRJNA224116 | 1487953 | *Leptolyngbya* sp. JSC-1 |
| GCF_000332095.2 | PRJNA224116 | 1173264 | *Leptolyngbya* sp. PCC 6406 |
| GCF_000763385.1 | PRJNA224116 | 1229172 | *Leptolyngbya* sp. KIOST-1 |
| GCF_000309385.1 | PRJNA224116 | 118166 | *Nodosilinea nodulosa* PCC 7104 |
| GCF_000155595.1 | PRJNA224116 | 91464 | *Synechococcus* sp. PCC 7335 |
| GCF_000482245.1 | PRJNA224116 | 1385935 | *Leptolyngbya* sp. Heron Island J |
| GCF_000316115.1 | PRJNA224116 | 102129 | *Leptolyngbya* sp. PCC 7375 |
| GCF_000464785.1 | PRJNA224116 | 1255374 | *Planktothrix rubescens* NIVA-CYA 407 |
| GCF_000175415.3 | PRJNA224116 | 634502 | *Arthrospira platensis* str. Paraca |
| GCF_000478195.2 | PRJNA224116 | 1348334 | *Lyngbya aestuarii* BL J |
| GCF_000332155.1 | PRJNA224116 | 402777 | *Kamptonema formosum* PCC 6407 |
| GCF_000317475.1 | PRJNA224116 | 179408 | *Oscillatoria nigro-viridis* PCC 7112 |
| GCF_000317105.1 | PRJNA224116 | 56110 | *Oscillatoria acuminata* PCC 6304 |
| GCF_000317515.1 | PRJNA224116 | 1173027 | *Microcoleus* sp. PCC 7113 |
| GCF_000021825.1 | PRJNA224116 | 65393 | *Cyanothece* sp. PCC 7424 |
| GCA_000307995.2 | PRJEA88171 | 1160280 | *Microcystis aeruginosa* PCC 9432 |
| GCF_000021805.1 | PRJNA224116 | 41431 | *Cyanothece* sp. PCC 8801 |
| GCF_000737945.1 | PRJNA256120 | 1527444 | *Candidatus Atelocyanobacterium thalassa* isolate SIO64986 |
| GCF_000284135.1 | PRJNA224116 | 1080228 | *Synechocystis* sp. PCC 6803 substr. GT-I |
| GCF_000715475.1 | PRJNA224116 | 490193 | *Synechococcus* sp. NKBG042902 |
| GCF_000317655.1 | PRJNA39697 | 292563 | *Cyanobacterium stanieri* PCC 7202 |
| GCF_000332055.1 | PRJNA224116 | 102125 | *Xenococcus* sp. PCC 7305 |
| GCF_000317575.1 | PRJNA224116 | 111780 | *Stanieria cyanosphaera* PCC 7437 |
| GCF_000380225.1 | PRJNA224116 | 1128427 | filamentous cyanobacterium ESFC-1 |
| GCF_000317615.1 | PRJNA224116 | 13035 | *Dactylococcopsis salina* PCC 8305 |
| GCF_000317495.1 | PRJNA224116 | 1173022 | *Crinalium epipsammum* PCC 9333 |
| GCF_000317555.1 | PRJNA224116 | 1173026 | *Gloeocapsa* sp. PCC 7428 |
| GCF_000317125.1 | PRJNA224116 | 251229 | *Chroococcidiopsis thermalis* PCC 7203 |
| GCF_000582685.1 | PRJNA224116 | 1469607 | [*Scytonema hofmanni*] UTEX 2349 |
| GCF_000789435.1 | PRJNA224116 | 1532906 | *Aphanizomenon flos-aquae* 2012/KM1/D3 |
| GCF_000196515.1 | PRJNA224116 | 551115 | '*Nostoc azollae*' 0708 |
| GCF_000316645.1 | PRJNA224116 | 28072 | *Nostoc* sp. PCC 7524 |
| GCF_000204075.1 | PRJNA10642 | 240292 | *Anabaena variabilis* ATCC 29413 |
| GCA_000340565.3 | PRJNA185469 | 313624 | *Nodularia spumigena* CCY9414 |
| GCF_000020025.1 | PRJNA224116 | 63737 | *Nostoc punctiforme* PCC 73102 |
| GCF_000332295.1 | PRJNA224116 | 643473 | *Fortiea contorta* PCC 7126 |
| GCF_000346485.2 | PRJNA224116 | 128403 | *Scytonema hofmannii* PCC 7110 |
| GCF_000734895.2 | PRJNA224116 | 1337936 | *Calothrix* sp. 336/3 |
| GCF_000332255.1 | PRJNA224116 | 1173021 | cyanobacterium PCC 7702 |
| GCF_000317225.1 | PRJNA224116 | 98439 | *Fischerella thermalis* PCC 7521 |
| GCF_000012525.1 | PRJNA224116 | 1140 | *Synechococcus elongatus* PCC 7942 |
| GCF_000586015.1 | PRJNA224116 | 1451353 | *Candidatus Synechococcus spongiorum* SH4 |
| GCF_000155635.1 | PRJNA224116 | 180281 | *Cyanobium* sp. PCC 7001 |
| GCA_000015705.1 | PRJNA13496 | 59922 | *Prochlorococcus marinus* str. MIT 9303 |
| GCF_000011485.1 | PRJNA224116 | 74547 | *Prochlorococcus marinus* str. MIT 9313 |
| GCF_000153805.1 | PRJNA224116 | 313625 | *Synechococcus* sp. BL107 |

To study the nature of the organisms co-cultivated in the cyanobacterial cultures, we relied on the release 1.4.0 of the RiboDB database (40) as a taxonomic reference. To this end, the 53 files corresponding to ribosomal proteins occurring in Bacteria were downloaded and aligned with MAFFT. The script ali2phylip.pl (part of Bio-MUST-Core) was then used to discard alignment sites with >50% missing character states. Concatenation of the 53 alignments with SCaFoS yielded a supermatrix of 3474 organisms x 6612 unambiguously aligned amino-acid positions (5.4% missing character states) that was used to infer a fast preliminary tree with RAxML v8.1.17 (41) under the LG4X model (data not shown). This large ribosomal protein tree allowed us to select representative organisms based on patristic distances in order to maximize diversity. At a minimum distance of 0.7 substitution/site, 200 organisms were retained using treeplot (from the MUST software package; (42)). Visual inspection of the tree inferred from this smaller dataset led us to further discard 4 fast-evolving organisms, yielding a total of 196 representative organisms. Both the large (3474 organisms) and the small (196 organisms) datasets were used in subsequent analyses. Hence, the 53 alignments (both large and small versions) were enriched (using again “42”) with sequences from the foreign (i.e., non-cyanobacterial) bins assembled from our 17 cyanobacterial cultures (31 bins in total, excluding unclassified CheckM bins). To control the origins of the enriching sequences, taxonomic filters of “42” were enabled, so as to require all new sequences to belong to the taxon determined by CheckM during its analysis of each whole bin. After this step, 4 incomplete genome bins (ULC066-bin3, ULC073-bin4, ULC082-bin4, ULC146-bin6) were discarded due to their low prevalence in the alignments (<10%). Enriched alignments were then processed as above with either ali2phylip.pl (large dataset) or BMGE (small dataset). The two resulting supermatrices assembled with SCaFoS contained 3501 organisms x 6613 unambiguously aligned amino-acid positions (6.0% missing character states) and 223 organisms x 7060 unambiguously aligned amino-acid positions (7.8% missing character states), respectively. Finally, two different trees were inferred using either RAxML (large dataset) or PhyloBayes (small dataset).

All phylogenetic trees were formatted using the script format-tree.pl (part of Bio-MUST-Core), FigTree v1.4.2 (http://tree.bio.ed.ac.uk/software/figtree/) and further arranged in InkScape v0.92 (43).

### SSU rRNA (16S) analyses

SSU rRNA (16S) genes were predicted using RNAmmer v1.2 (44) in all genome bins for the selected MetaBAT parameter set. Beyond regular bins, we also investigated an additional bin (called nobin) for each metagenome, which contained all the scaffolds rejected by MetaBAT during the binning process. Predicted rRNA sequences were taxonomically classified by SINA v1.2.11 (45), using the release 128 of the SILVA database composed of 1,922,213 SSU rRNA reference sequences (46).

## Results

### Metagenome sequencing and assembly

We obtained a total of 55 different genome bins from the separate sequencing and metagenomic assembly of the 17 cyanobacterial cultures (**Table 3**). Among those, we identified 15 bins as cyanobacterial (ULC007-bin1, ULC027-bin1, ULC041-bin1, ULC065-bin1, ULC066-bin1, ULC068-bin1, ULC073-bin1, ULC077-bin1, ULC082-bin1, ULC084-bin3, ULC129-bin1, ULC165-bin4, ULC186-bin1, ULC187-bin1, ULC335-bin1), based on CheckM classification (Parks et al., 2015), except for ULC165-bin4, which was classified after DIAMOND blastx results. For the two Nostocales strains (ULC146 and ULC179), we failed to recover any cyanobacterial bin (see below however for the analysis of the other bins). In each metagenome, the cyanobacterial bin nearly always corresponded to the largest predicted bin, both in terms of total length and sequencing coverage (**Table 3**; see also **Figure S1**). For two cultures, however, cyanobacterial bins were the smallest predicted (ULC084-bin3 and ULC165-bin4). Genome completeness, evaluated with CheckM, was >90% (median = 97.74%, IQR = 4.04%) for all cyanobacterial bins but lower for ULC165-bin4 (24.14%). As expected, completeness positively correlated with the sequencing coverage of the bins in the metagenomic assemblies, but this correlation was barely significant (Pearson r = 0.52, P-value = 0.05). The contamination level was evaluated to be <1.63% (median = 0.47%, IQR = 0.83%) with CheckM and <2.62% (median = 1.26%, IQR = 0.40%) with our DIAMOND blastx parser (Cornet et al., 2018, in revision). As our libraries were only composed of paired-ends (and not of mate pairs), the number of scaffolds obtained after metaSPAdes assembly and SSPACE scaffolding remained quite high for all cyanobacterial genome bins (≥60, median = 238, IQR = 292) (**Tables 3** and **S2**).

**Table 3:**
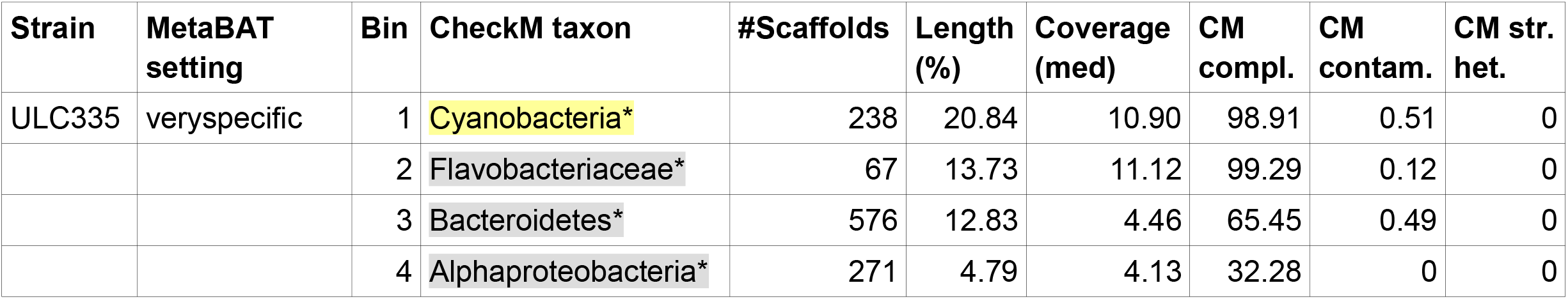
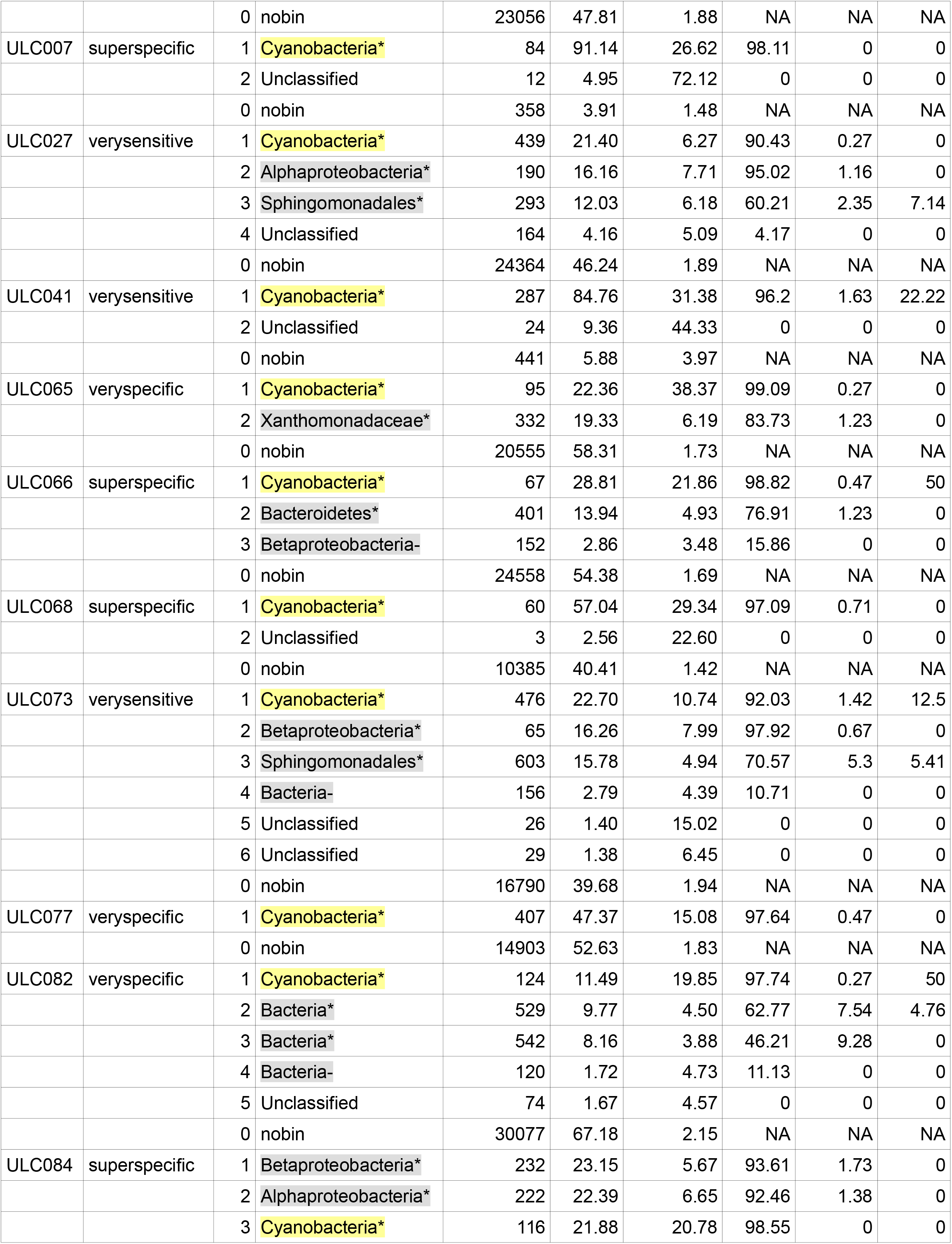
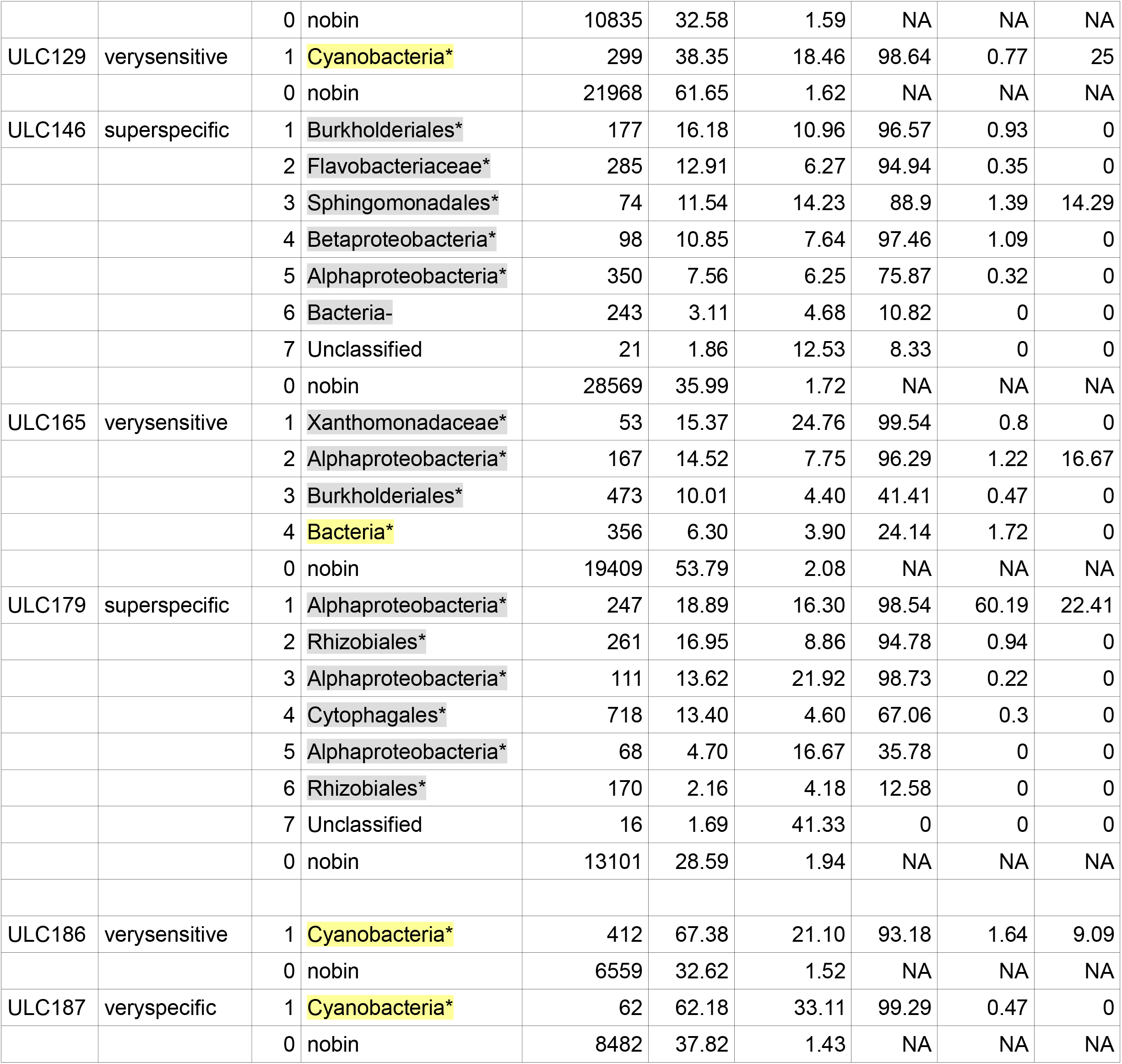
Assembly statistics, taxonomy, completeness, contamination and coverage of genome bins. The taxonomic label (CheckM taxon), the genome completeness (CM compl.), the contamination level (CM contam. and CM str. het.) were computed with CheckM. Sequencing coverage (med) was computed with BBMap, while Length was extracted from QUAST output. Genome bins used in phylogenetic inference are marked by an (*) and discarded bins by an (-).

| Strain | MetaBAT setting | Bin | CheckM taxon | #Scaffolds | Length (%) | Coverage (med) | CM compl. | CM contam. | CM str. het. |
| --- | --- | --- | --- | --- | --- | --- | --- | --- | --- |
| ULC335 | veryspecific | 1 | Cyanobacteria* | 238 | 20.84 | 10.90 | 98.91 | 0.51 | 0 |
|  |  | 2 | Flavobacteriaceae* | 67 | 13.73 | 11.12 | 99.29 | 0.12 | 0 |
|  |  | 3 | Bacteroidetes* | 576 | 12.83 | 4.46 | 65.45 | 0.49 | 0 |
|  |  | 4 | Alphaproteobacteria* | 271 | 4.79 | 4.13 | 32.28 | 0 | 0 |
|  |  | 0 | nobin | 23056 | 47.81 | 1.88 | NA | NA | NA |
| ULC007 | superspecific | 1 | Cyanobacteria* | 84 | 91.14 | 26.62 | 98.11 | 0 | 0 |
|  |  | 2 | Unclassified | 12 | 4.95 | 72.12 | 0 | 0 | 0 |
|  |  | 0 | nobin | 358 | 3.91 | 1.48 | NA | NA | NA |
| ULC027 | verysensitive | 1 | Cyanobacteria* | 439 | 21.40 | 6.27 | 90.43 | 0.27 | 0 |
|  |  | 2 | Alphaproteobacteria* | 190 | 16.16 | 7.71 | 95.02 | 1.16 | 0 |
|  |  | 3 | Sphingomonadales* | 293 | 12.03 | 6.18 | 60.21 | 2.35 | 7.14 |
|  |  | 4 | Unclassified | 164 | 4.16 | 5.09 | 4.17 | 0 | 0 |
|  |  | 0 | nobin | 24364 | 46.24 | 1.89 | NA | NA | NA |
| ULC041 | verysensitive | 1 | Cyanobacteria* | 287 | 84.76 | 31.38 | 96.2 | 1.63 | 22.22 |
|  |  | 2 | Unclassified | 24 | 9.36 | 44.33 | 0 | 0 | 0 |
|  |  | 0 | nobin | 441 | 5.88 | 3.97 | NA | NA | NA |
| ULC065 | veryspecific | 1 | Cyanobacteria* | 95 | 22.36 | 38.37 | 99.09 | 0.27 | 0 |
|  |  | 2 | Xanthomonadaceae* | 332 | 19.33 | 6.19 | 83.73 | 1.23 | 0 |
|  |  | 0 | nobin | 20555 | 58.31 | 1.73 | NA | NA | NA |
| ULC066 | superspecific | 1 | Cyanobacteria* | 67 | 28.81 | 21.86 | 98.82 | 0.47 | 50 |
|  |  | 2 | Bacteroidetes* | 401 | 13.94 | 4.93 | 76.91 | 1.23 | 0 |
|  |  | 3 | Betaproteobacteria- | 152 | 2.86 | 3.48 | 15.86 | 0 | 0 |
|  |  | 0 | nobin | 24558 | 54.38 | 1.69 | NA | NA | NA |
| ULC068 | superspecific | 1 | Cyanobacteria* | 60 | 57.04 | 29.34 | 97.09 | 0.71 | 0 |
|  |  | 2 | Unclassified | 3 | 2.56 | 22.60 | 0 | 0 | 0 |
|  |  | 0 | nobin | 10385 | 40.41 | 1.42 | NA | NA | NA |
| ULC073 | verysensitive | 1 | Cyanobacteria* | 476 | 22.70 | 10.74 | 92.03 | 1.42 | 12.5 |
|  |  | 2 | Betaproteobacteria* | 65 | 16.26 | 7.99 | 97.92 | 0.67 | 0 |
|  |  | 3 | Sphingomonadales* | 603 | 15.78 | 4.94 | 70.57 | 5.3 | 5.41 |
|  |  | 4 | Bacteria- | 156 | 2.79 | 4.39 | 10.71 | 0 | 0 |
|  |  | 5 | Unclassified | 26 | 1.40 | 15.02 | 0 | 0 | 0 |
|  |  | 6 | Unclassified | 29 | 1.38 | 6.45 | 0 | 0 | 0 |
|  |  | 0 | nobin | 16790 | 39.68 | 1.94 | NA | NA | NA |
| ULC077 | veryspecific | 1 | Cyanobacteria* | 407 | 47.37 | 15.08 | 97.64 | 0.47 | 0 |
|  |  | 0 | nobin | 14903 | 52.63 | 1.83 | NA | NA | NA |
| ULC082 | veryspecific | 1 | Cyanobacteria* | 124 | 11.49 | 19.85 | 97.74 | 0.27 | 50 |
|  |  | 2 | Bacteria* | 529 | 9.77 | 4.50 | 62.77 | 7.54 | 4.76 |
|  |  | 3 | Bacteria* | 542 | 8.16 | 3.88 | 46.21 | 9.28 | 0 |
|  |  | 4 | Bacteria- | 120 | 1.72 | 4.73 | 11.13 | 0 | 0 |
|  |  | 5 | Unclassified | 74 | 1.67 | 4.57 | 0 | 0 | 0 |
|  |  | 0 | nobin | 30077 | 67.18 | 2.15 | NA | NA | NA |
| ULC084 | superspecific | 1 | Betaproteobacteria* | 232 | 23.15 | 5.67 | 93.61 | 1.73 | 0 |
|  |  | 2 | Alphaproteobacteria* | 222 | 22.39 | 6.65 | 92.46 | 1.38 | 0 |
|  |  | 3 | Cyanobacteria* | 116 | 21.88 | 20.78 | 98.55 | 0 | 0 |
|  |  | 0 | nobin | 10835 | 32.58 | 1.59 | NA | NA | NA |
| ULC129 | verysensitive | 1 | Cyanobacteria* | 299 | 38.35 | 18.46 | 98.64 | 0.77 | 25 |
|  |  | 0 | nobin | 21968 | 61.65 | 1.62 | NA | NA | NA |
| ULC146 | superspecific | 1 | Burkholderiales* | 177 | 16.18 | 10.96 | 96.57 | 0.93 | 0 |
|  |  | 2 | Flavobacteriaceae* | 285 | 12.91 | 6.27 | 94.94 | 0.35 | 0 |
|  |  | 3 | Sphingomonadales* | 74 | 11.54 | 14.23 | 88.9 | 1.39 | 14.29 |
|  |  | 4 | Betaproteobacteria* | 98 | 10.85 | 7.64 | 97.46 | 1.09 | 0 |
|  |  | 5 | Alphaproteobacteria* | 350 | 7.56 | 6.25 | 75.87 | 0.32 | 0 |
|  |  | 6 | Bacteria- | 243 | 3.11 | 4.68 | 10.82 | 0 | 0 |
|  |  | 7 | Unclassified | 21 | 1.86 | 12.53 | 8.33 | 0 | 0 |
|  |  | 0 | nobin | 28569 | 35.99 | 1.72 | NA | NA | NA |
| ULC165 | verysensitive | 1 | Xanthomonadaceae* | 53 | 15.37 | 24.76 | 99.54 | 0.8 | 0 |
|  |  | 2 | Alphaproteobacteria* | 167 | 14.52 | 7.75 | 96.29 | 1.22 | 16.67 |
|  |  | 3 | Burkholderiales* | 473 | 10.01 | 4.40 | 41.41 | 0.47 | 0 |
|  |  | 4 | Bacteria* | 356 | 6.30 | 3.90 | 24.14 | 1.72 | 0 |
|  |  | 0 | nobin | 19409 | 53.79 | 2.08 | NA | NA | NA |
| ULC179 | superspecific | 1 | Alphaproteobacteria* | 247 | 18.89 | 16.30 | 98.54 | 60.19 | 22.41 |
|  |  | 2 | Rhizobiales* | 261 | 16.95 | 8.86 | 94.78 | 0.94 | 0 |
|  |  | 3 | Alphaproteobacteria* | 111 | 13.62 | 21.92 | 98.73 | 0.22 | 0 |
|  |  | 4 | Cytophagales* | 718 | 13.40 | 4.60 | 67.06 | 0.3 | 0 |
|  |  | 5 | Alphaproteobacteria* | 68 | 4.70 | 16.67 | 35.78 | 0 | 0 |
|  |  | 6 | Rhizobiales* | 170 | 2.16 | 4.18 | 12.58 | 0 | 0 |
|  |  | 7 | Unclassified | 16 | 1.69 | 41.33 | 0 | 0 | 0 |
|  |  | 0 | nobin | 13101 | 28.59 | 1.94 | NA | NA | NA |
| ULC186 | verysensitive | 1 | Cyanobacteria* | 412 | 67.38 | 21.10 | 93.18 | 1.64 | 9.09 |
|  |  | 0 | nobin | 6559 | 32.62 | 1.52 | NA | NA | NA |
| ULC187 | veryspecific | 1 | Cyanobacteria* | 62 | 62.18 | 33.11 | 99.29 | 0.47 | 0 |
|  |  | 0 | nobin | 8482 | 37.82 | 1.43 | NA | NA | NA |

Altogether, we identified 40 bins that were not of cyanobacterial origin out of our 17 cyanobacterial cultures. Among these foreign genome bins, we classified 21 bins as Proteobacteria and 5 as Bacteroidetes, thus 26 bins contained organisms belonging to bacterial phyla known to participate in the cyanobacterial microbiome. The 14 remaining bins could only be classified as Bacteria (5) or were left unclassified (9) by CheckM. While unclassified bins were discarded from subsequent analyses, bins identified at the Bacteria level were retained. Genome completeness of these 31 bacterial bins was very heterogeneous (median = 71.96%, IQR = 51.84%). As for cyanobacterial bins, but more significantly, completeness positively correlated with sequencing coverage, lowly covered bins being the less complete (Pearson r = 0.46, P-value = 0.007). Nevertheless, we managed to recover 13 nearly complete foreign bins (completeness >90%). According to CheckM, the contamination level (foreign sequences not belonging to the taxonomic label of the bin under study) of the 26 classified non-cyanobacterial bins was always <9.28% (median = 0.8%, IQR= 1.13%), except for ULC179-bin1 (60.19%). The contamination level of the bins classified as Bacteria was not recorded, because such a high taxonomic rank made its evaluation meaningless. As for cyanobacterial bins, the number of scaffolds of the 31 bacterial bins remained quite high (>53, median = 232, IQR = 205). In spite of three cases of possible complementarity (in terms of recovered marker genes) suggested by CheckM (ULC027-bin3/ULC027-bin4, ULC146-bin3/ULC146-bin7 and ULC082-bin3/ULC082-bin4), the two first involving unclassified bins, the corresponding bins were not merged because CheckM phylogenetic placement was never congruent. Details about genome bins are available in **Table S2**. We released scaffolded assemblies and protein predictions for all the bins having a completeness >90%, whether classified as cyanobacterial (14) or probable microbiome organisms (13). Raw read data for the 17 cyanobacterial cultutes have been deposited in the NCBI SRA (SAMN08623419 to SAMN08623435; bioproject PRJNA436342). Deposition of our 27 most complete (>90%) genome assemblies and of the corresponding annotations is underway.

### Cyanobacterial phylogenomics

A phylogenomic analysis based on 675 genes and 64 reference Cyanobacteria showed that 14 cyanobacterial bins (i.e., excluding ULC335) were scattered over the basal part of the cyanobacterial tree (**Figure 1**). Statistical support (Bayesian posterior probabilities = PP) was maximal except for three nodes. In the following, we refer to the cyanobacterial clades using the nomenclature defined in Shih et al. (2013), since the latter was the first study to fully sample the cyanobacterial morphological diversity [i.e., Sections I-V from (1)]. Three ULC strains (*Pseudanabaena* sp. ULC187, *Pseudanabaena frigida* ULC066 and *Leptolyngbya* sp. ULC068), are located at a very basal (i.e., “early-branching”) position in clade F, and form a cluster with the reference strain *Pseudanabaena biceps* GCF_000332215.1. Three others strains, identified as *Cyanobium* sp. (ULC065, ULC082 and ULC084), emerge together from the picocyanobacteria clade C1. Although their C1 membership is indisputable, the exact branching point within the clade C1 is not resolved (PP = 0.51). The six *Leptolyngbya* strains (*Leptolyngbya* sp. ULC077/ULC165/ULC186, *L. antarctica* ULC041, *L. glacialis* ULC073 and *L. foveolarum* ULC129) and the two *Phormidesmis/Phormidium priestleyi* (ULC007 and ULC027) are located in clade C3, mainly composed of reference *Leptolyngbya* strains. While two strains (*Leptolyngbya* sp. ULC077 and ULC165) each form an additional single branch within clade C3, five other strains emerge as two new sub-groups: *Leptolyngbya foveolarum* ULC129 and *Phormidium priestleyi* ULC027 on the one hand (yet weakly supported: PP = 0.51), and *Leptolyngbya* sp. ULC186, *Leptolyngbya antarctica* ULC041 and *Leptolyngbya glacialis* ULC073 on the other hand. As expected, our new assembly of *Phormidesmis priestleyi* ULC007 is extremely close to the first release of the same genome [*Phormidesmis priestleyi* GCF_001895925.1 (47)]. Finally, *Snowella* sp. ULC335 is part of clade B2, composed of various cyanobacterial genera from the orders Pleurocapsales and Chroococales (48), again with maximal support.

**Figure 1:**
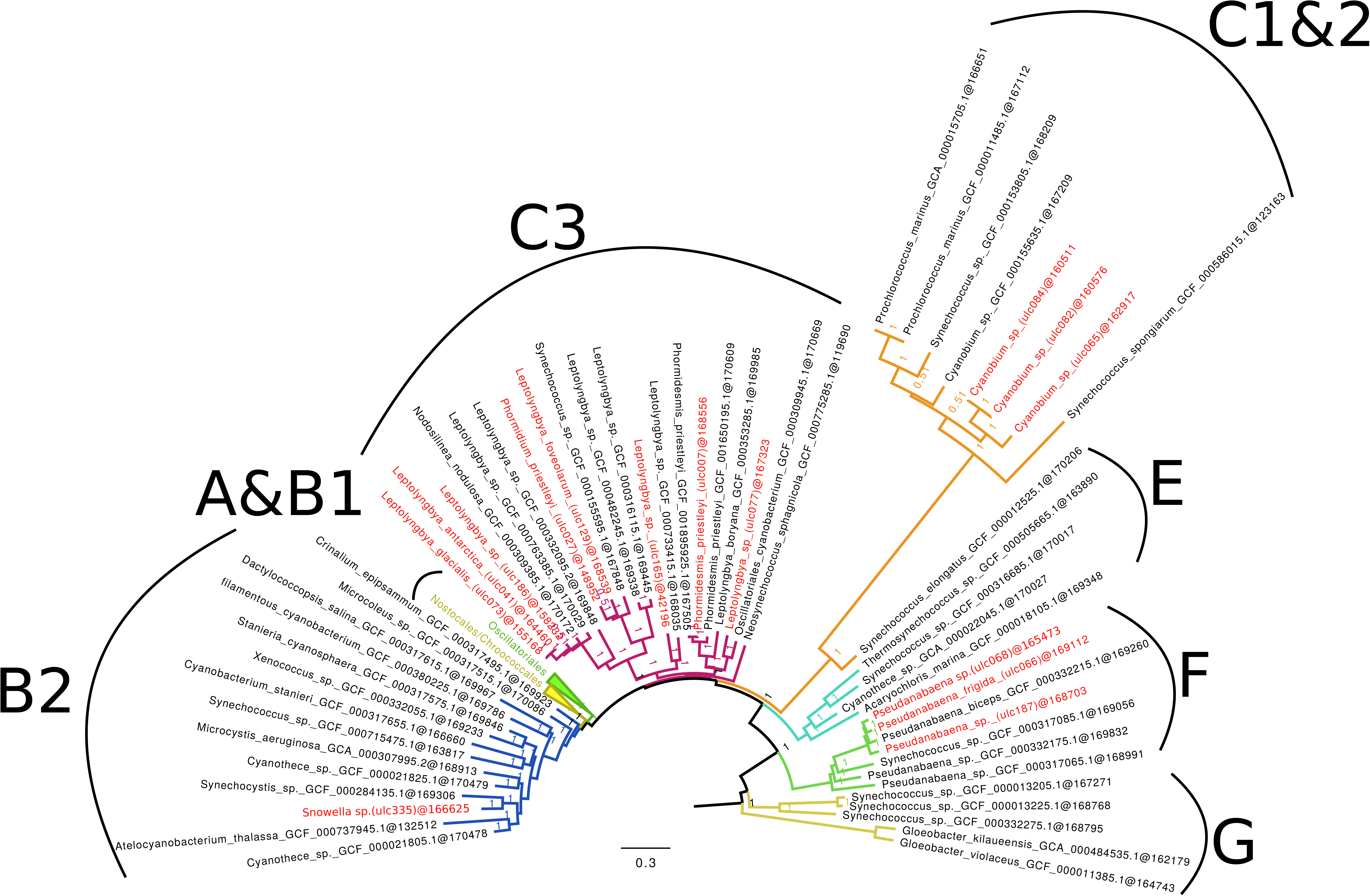
Phylogenomic tree of 64 broadly sampled Cyanobacteria showing the phylogenetic position of the 15 cyanobacterial genome bins. The Bayesian tree was inferred under the CAT+Γ_4_ model from a supermatrix made of 675 genes (79 organisms x 170,983 amino-acid positions). Cyanobacterial clades were named according to the classification of Shih et al. (2013). Trailing numbers in tip labels give the number of amino-acid positions effectively present in the corresponding concatenated sequence, whereas numbers at nodes are posterior probabilities (PP) computed from two independent chains. Genome bins are shown in red.

### Microbiome phylogenomics

To identify the organisms in the putative microbiome bins recovered from the 17 cultures, we built two phylogenomic trees with different taxon samplings of reference prokaryotes from a concatenation of 53 ribosomal proteins (see Materials and Methods). **Figure 2** shows the small tree (193 Bacteria and 30 Archaea), surrounded by zooms in specific regions of the large tree (3374 Bacteria and 127 Archaea). Only 27 bins out of 31 non-cyanobacterial could be included in the tree, 4 bins (marked by a dash in **Table 3**) being too incomplete to be positioned robustly (see Materials and Methods). The resolution of the small tree was quite good, with 78% of the nodes having PPs ≥0.90 and no node having a PP <0.50. This analysis showed that all 27 analyzed microbiome bins fall either in Bacteroidetes (5 bins) or in Proteobacteria (14 bins in Alpha-, 5 bins in Beta- and 3 bins in Gammaproteobacteria) (**Figure 2**), the tree allowing us to precise the CheckM ‘bacterial’ affiliation of ULC082-bin3 to Gammaproteobacteria. In all cases, microbiome bins were sisters to one or more of the representative organisms with PP ≥0.99, except for ULC179-bin3 (PP = 0.63). Insets A-C of **Figure 2** demonstrate that the five Bacteroidetes bins correspond to different organisms, despite the fact that they appear closely clustered in the small tree. However, the picture is different for the bins falling in Proteobacteria (insets E-H). Whereas they are globally scattered across the phylum, there exist five cases (involving 11 bins) for which two or three bins from different cyanobacterial cultures appear extremely close in the large tree: ULC073-bin2/ULC084-bin1/ULC146-bin4 (D), ULC146-bin1/ULC165-bin3 (D), ULC065-bin2/ULC165-bin1 (E), ULC027-bin3/ULC146-bin3 (G) and ULC084-bin2/ULC165-bin2 (H). Taking this into account, the 27 microbiome bins only create 21 terminal branches in the large tree, five of them clustering with a reference strain of *Brevundimonas subvibrioides* (H).

**Figure 2:**
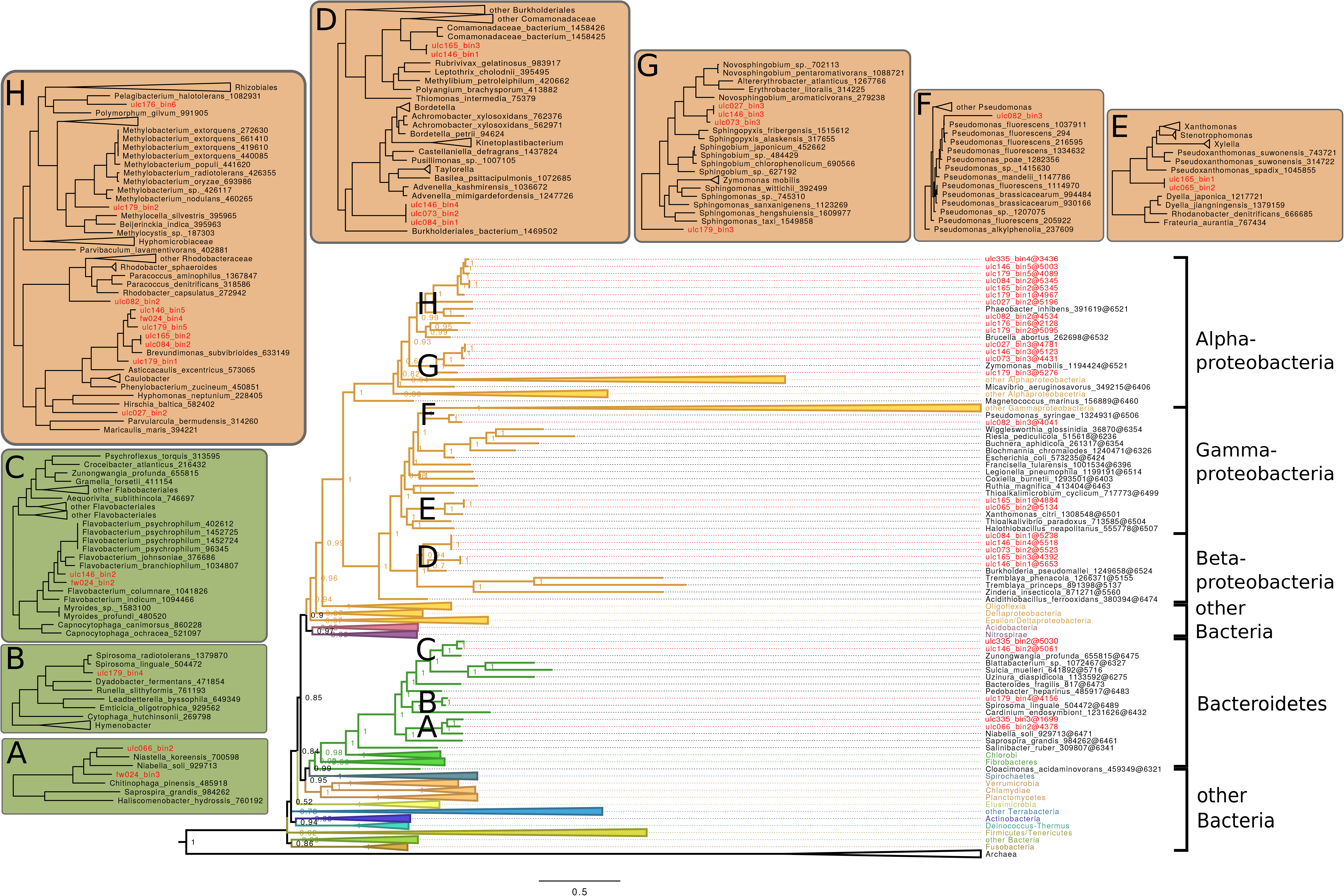
Phylogenomic tree of 196 broadly sampled Bacteria and Archaea showing the phylogenetic position of 27 microbiome genome bins. The Bayesian tree was inferred under the CAT+Γ_4_ model from a supermatrix made of 53 ribosomal genes (223 organisms x 7060 amino-acid positions). Surrounding subtrees are excerpts from a large maximum-likelihood tree inferred under the LG4X model from the full supermatrix (3501 organisms x 6613 amino-acid positions).

### SSU rRNA (16S) analyses

In an attempt to refine the taxonomic analysis of all our genome bins, we predicted their SSU rRNA (16S) with RNAmmer (44). Hence, we managed to predict 38 sequences (**Table 4**). Unfortunately, the vast majority (33) of the rRNA genes were predicted from unbinned metagenomic contigs (nobins; see Materials and Methods). When the taxon corresponding to the rRNA was straightforward to match with the taxon of one of the bins from the same cyanobacterial culture (based on congruent CheckM and SINA classifications), we manually affiliated the rRNA gene to that bin. This was possible for 20 predicted rRNA genes, but 13 sequences could not be reliably affiliated to any genome bin (empty cells in **Table 4**). According to SINA (45), only 10 of the predicted SSU rRNA genes were of cyanobacterial origin, whereas 8 sequences were left unclassified. The 20 remaining sequences were of either Proteobacteria or Bacteroidetes origin, thereby confirming the results of our phylogenomic analysis of microbiome bins based on rRNA proteins. Two best hits were encountered more than once by SINA: *Blastomonas* sp. AAP25 (from a Czech freshwater lake) in ULC073-bin6 and ULC146-bin3, and ‘Uncultured bacterium’ clone B3NR69D12 (from a drinking water biofilm) in ULC073-bin2 and ULC084-bin1.

**Table 4:** SSU rRNA (16S) gene prediction, taxonomy and coverage. The LCA classification and top hits were retrieved from SINA analyses. The bins with SSU rRNA (16S) genes directly predicted from the genome bins (without manual assignment) are indicated by *. Coverage values were computed with BBMap.

| Strain | SSUref_128 taxon | SSUref_128 top hit | Bin affiliation |  | Coverage |
| --- | --- | --- | --- | --- | --- |
| ULC335 | Snowella | *Snowella litoralis* 1LT47S05 | bin0 | bin1 | 37.00 |
| ULC335 | Brevundimonas | Uncultured *Brevundimonas* sp. | bin0 |  | 22.23 |
| ULC335 | Flavobacterium | Uncultured bacterium clone N4_091 | bin0 | bin2 | 58.44 |
| ULC335 | Unclassified | NA | bin0 |  | 10.25 |
| ULC335 | Hydrogenophaga | *Hydrogenophaga palleronii* | bin0 |  | 9.64 |
| ULC335 | Rhodobacteraceae | Uncultured bacterium clone ZWB3-3 | bin0 |  | 7.79 |
| ULC007 | Leptolyngbya | *Phormidesmis priestleyi* ANT.LG2.4 16S | bin0 | bin1 | 85.23 |
| ULC027 | Unclassified | NA | bin2 | bin2* | 54.08 |
| ULC041 | Leptolyngbya | *Leptolyngbya antarctica* ANT.LACV6.1 | bin0 | bin1 | 97.23 |
| ULC065 | Arenimonas | Uncultured bacterium clone a33 | bin0 | bin2 | 40.66 |
| ULC065 | Synechococcus | *Cyanobium* sp. JJ17-5 | bin0 | bin1 | 165.13 |
| ULC066 | Limnobacter | Uncultured bacterium clone S25 | bin0 | bin3 | 14.15 |
| ULC066 | Unclassified | NA | bin0 |  | 21.51 |
| ULC066 | FamilyI | *Pseudanabaena biceps* PCC 7429 | bin0 | bin1 | 50.06 |
| ULC068 | FamilyI | *Pseudanabaena* sp. Sai012 | bin0 | bin1 | 68.53 |
| ULC073 | Sphingomonadaceae | *Blastomonas* sp. AAP25 | bin6 | bin6* | 31.87 |
| ULC073 | Leptolyngbya | *Leptolyngbya antarctica* ANT.LACV6.1 | bin0 | bin1 | 33.60 |
| ULC073 | Limnobacter | Uncultured bacterium clone B3NR69D12 | bin0 | bin2 | 19.58 |
| ULC077 | Unclassified | NA | bin0 | bin1 | 52.80 |
| ULC082 | Hydrogenophaga | Uncultured Comamonadaceae bacterium | bin0 |  | 18.85 |
| ULC082 | Brevundimonas | Uncultured alpha proteobacterium clone KWK6S.50 | bin0 |  | 25.08 |
| ULC082 | Unclassified | NA | bin0 |  | 32.77 |
| ULC082 | Pseudomonas | *Pseudomonas* sp. WCS374 | bin0 |  | 32.71 |
| ULC082 | Synechococcus | *Synechococcus* sp. MW97C4 | bin0 | bin1 | 93.93 |
| ULC084 | Brevundimonas | Uncultured alpha proteobacterium | bin0 | bin2 | 31.39 |
| ULC084 | Synechococcus | Uncultured bacterium clone MS81 | bin0 | bin3 | 87.30 |
| ULC084 | Limnobacter | Uncultured bacterium clone B3NR69D12 | bin0 | bin1 | 16.99 |
| ULC129 | Phormidium | Uncultured bacterium clone GBII-52 | bin0 | bin1 | 52.71 |
| ULC146 | Sphingomonadaceae | *Blastomonas* sp. AAP25 | bin3 | bin3* | 81.39 |
| ULC146 | Flavobacterium | *Flavobacterium* sp. Leaf359 | bin0 | bin2 | 25.91 |
| ULC146 | Hydrogenophaga | *Hydrogenophaga* sp. Root209 | bin1 | bin1* | 61.13 |
| ULC165 | Unclassified | NA | bin0 |  | 85.20 |
| ULC165 | Unclassified | NA | bin0 |  | 98.62 |
| ULC179 | Devosia | *Devosia psychrophila* strain Cr7-05 | bin0 |  | 97.88 |
| ULC179 | Unclassified | NA | bin0 |  | 16.43 |
| ULC179 | Polymorphobacter | Uncultured Sphingomonadaceae bacterium | bin3 | bin3* | 91.23 |
| ULC186 | FamilyI | *Leptolyngbya* sp. 0BB32S02 | bin0 | bin1 | 116.04 |
| ULC187 | FamilyI | *Pseudanabaena* sp. Sai010 | bin0 | bin1 | 81.01 |

## Discussion

According to the classification criteria of Bowers et al. (2017) (24), the vast majority (14) of the cyanobacterial bins are of medium-quality, since their genome completeness is >90% and their contamination level <5% (both with CheckM and DIAMOND blastx). Yet, they are still composed of a large number of scaffolds (≥60), due to the use of short insert DNA libraries for sequencing (**Tables 3** and **S2**). In contrast, the only low-quality cyanobacterial assembly obtained here (ULC165-bin4) shows a completeness of 24.14%, in agreement with the lowest coverage obtained over all 4 ULC165 bins (3.90%). The situation is worse with the two Nostocales cultures (ULC146 and ULC179), for which we could not isolate any cyanobacterial bin. This lack of cyanobacterial contigs can be explained by the fact that these three strains produce a thick polysaccharidic sheath that hinders DNA extraction (1). Such a thick sheath is thought to protect the organisms from the harsh conditions of their hostile environment (Sør Rondane Mountains in Antarctica in all three cases).

When MetaBAT partitioned the metagenomic contigs, it produced 9 small bins that were left unclassified by CheckM. In two cases, unclassified bins were identified as complementary (of CheckM marker genes) to another bin from the same metagenome (ULC027-bin3/ULC027-bin4; ULC146-bin3/ULC146-bin7; see above). Despite similar values in GC content and sequencing coverage, we did not merge these bins, thereby following the recommendations in the CheckM manual, because we had no indication about the phylogenetic affiliation of the unclassified bins. Since they only represented a very small fraction of the metagenomes, we discarded these bins from our phylogenetic analyses. Puzzlingly, such a bin was also recovered from the strain ULC007, for which no foreign bin was expected due to its axenicity. While the sequencing coverage of the unclassified bin (ULC007-bin2) was more than twice the coverage of the main bin (ULC007-bin1), tetranucleotide frequencies (TNFs) were undistinguishable between the two bins (**Figures S1** and **S2**). This suggests that the corresponding contigs originate from the same organism but that the small bin contains contigs encoded in multiple copies in the genome. We attempted to characterize some unclassified bins from a functional point a view using Prodigal (32) and Blast2GO (49). Unfortunately, results were largely inconclusive and we could not ascertain whether these bins (containing some transferases, e.g., acyltransferases, transferring one-carbon groups, transferring nitrogenous groups) correspond to aberrant chromosomal regions (e.g., laterally transferred segments, repetitive elements) or to plasmids (data not shown).

Even if our assemblies are globally of medium-quality, they often lack SSU rRNA (16S) genes. Hence, out of 38 predicted rRNA genes, as few as 5 were predicted from genome bins (of which only foreign bins), leaving 50 bins without any rRNA gene. Apparently, rRNA genes are rejected by MetaBAT, because we could only predict them from unbinned contigs (nobins) in all remaining cases (33). Importantly, this outcome was independent of the parameter set used for MetaBAT (data not shown). We nonetheless elected to favor this software because its binning performance in terms of completeness is better than that of other recent tools, such as CONCOCT (50), GroopM (51), MaxBin (52) and Canopy (53) [see Figure 3 of (26)]. Whenever SINA (45) successfully classified a predicted SSU rRNA (16S) gene, we did our best to manually affiliate it to the corresponding genome bin (**Table 4**). Consequently, 10 of our 15 cyanobacterial bins turned into high-quality genomes according to the classification of Bowers et al. (2017) (24). In this respect, it is worth mentioning that, among the 651 cyanobacterial genome assemblies available on the NCBI as of December 2017, only 458 have a SSU rRNA (16S) gene, based on RNAmmer (44) predictions (data not shown). According to our analyses, the frequent loss of rRNA genes is caused by the presence of multiple copies of the rRNA operon in many bacterial genomes (54), resulting in short rRNA-bearing contigs due to incomplete assembly of repeated regions. Since these contigs are dominated by the rRNA operon, they feature both a higher sequencing coverage and divergent TNFs, two properties that interfere with the binning process carried out by MetaBAT and other metagenomic software (**Supplemental Note 1**).

Our phylogenomic tree of Cyanobacteria is based on the largest dataset to date (64 clean and complete reference strains; >170,000 unambiguously aligned amino-acid positions). It is congruent with other recent cyanobacterial phylogenies (55, 56). Interestingly, all the cyanobacterial bins corresponding to arctic or subarctic strains (12 out of 15) are clearly located in the basal part of the tree. The BCCM/ULC collection has a focus on (sub)arctic cyanobacterial strains that may present interesting features to survive freeze/thaw cycles, seasonally contrasted light intensities, high UV radiations, desiccation and other stresses. Cyanobacterial diversity from such environments is presently underrepresented in comparison to that of marine Cyanobacteria. This is notably due to the difficulty of cultivating these organisms from “cold regions”, such as polar or alpine Cyanobacteria (13). Hence, increasing the sampling of (Cyano)bacteria from these environments may lead to a better understanding of their functional adaptation to environmental pressures, which is especially important in the context of climate change (13). Moreover, the three “early-branching” Pseudanabaena strains (ULC066, ULC068, ULC187 in clade F) should prove useful to improve the resolution of the phylogeny of Cyanobacteria in further studies by increasing their taxon sampling. Two of these strains were isolated from Canadian samples and ULC066 even originates from the Arctic (**Table 1**).

When the sequencing coverage was sufficient, we also assembled the foreign (i.e., non-cyanobacterial) bins. According to the classification of Bowers et al. (2017) (24), 13 of these bins are of medium-quality (completeness ≥90%) and 18 bins are of low-quality (completeness <90%) (**Table 3**). All are either of Proteobacteria or Bacteroidetes origin, as assessed by both CheckM and phylogenomic inference. From our phylogenomic analysis, it appears that the 27 analyzed bins represent 21 different terminal branches in the tree (**Figure 2**). As 11 were indistinguishable (or very closely related) in spite of the use of 53 ribosomal proteins, we investigated whether they represented genuinely different samplings of highly similar associated organisms or were the results of cross-contamination during Cyanobacteria isolation/cultivation or DNA processing (**Supplemental Note 2**). Altogether, genome-wide similarity measurements suggest that cross-contamination may not be involved, even if sampling sites were occasionally very distant (i.e., Arctic and Antarctic samples). Panel H of Figure 2 shows a group of six foreign bins clustered around a reference strain of *Brevundimonas subvibrioides*. As this Alpha-proteobacterium frequently appears as a last common ancestor taxon in SINA classifications of SSU rRNA (16S) sequences (**Table 4**), this indicates that *Brevundimonas* (or related taxa) is regularly present in ULC cultures and probably naturally associated to Cyanobacteria. More generally, the classification of all identifiable foreign bins as either Proteobacteria or Bacteroidetes suggest that the associated organisms come from the original environment and accompanied the Cyanobacteria through the isolation steps. Indeed, these two phyla are known to co-evolve with Cyanobacteria through complex trophic relations (21, 57). We probably identified only these two phyla in our foreign bins because they are the most abundant (21), whereas other associated bacterial phyla (Actinobacteria, Gemmatimonadetes, Planctomycetes, Verrucomicrobia) have been described in the cyanobacterial microbiome (15–17, 21). This result is completely in line with our recent analysis of the level of contamination in publicly available cyanobacterial genomes, in which foreign sequences were also mainly classified as Proteobacteria and Bacteroidetes (Cornet et al., 2018, in revision). In other words, the difficulty to purify non-axenic cyanobacterial cultures, possibly combined to the accidental transfer of associated bacteria during the isolation process (or any subsequent step), is probably the main cause for genome contamination. This certainly highlights the importance of careful bioinformatic protocols for genome data processing. In this respect, we compared our new assembly of ULC007 to the previous release of the same strain, based on a HiSeq run in addition to the MiSeq run used here (47). Interestingly, all CheckM values (completeness, contamination, strain heterogeneity) for ULC007-bin1 were slightly better than those obtained for our previously published assembly (completeness 98.11 vs 95.99, contamination 0 vs 1.18, Strain heterogeneity 0 vs 100). As the latter had used more primary data and had benefited from a thorough curation by hand, this indicates that the fully automated metagenomic pipeline of the present study is also applicable for axenic strains.

## Conclusion

In this work, we showed that a quite straightforward metagenomic protocol allows taking advantage of non-axenic cyanobacterial cultures. Our pipeline yields medium-quality genomes with a high level of completeness (high sensitivity) for a very low level of contaminant sequences (high specificity), which could be very useful for phylogenomic analyses. In contrast, it has the disadvantage of regularly discarding multi-copy SSU rRNA (16S) genes during the binning of metagenomic contigs. We have shown that this loss is due to their higher sequencing coverage and divergent TNFs, which are especially detrimental for short contigs. The metagenomic pipeline reported here has nevertheless the advantage of facilitating the assembly of cyanobacterial genomes, as long as enough genomic DNA can be extracted from the strains. Our results further indicate that the microbiome of different cultures can sometimes contain associated bacteria that are very closely related, even when sampling sites are very distant. Finally, we have released 14 novel cyanobacterial assemblies, including 11 (sub)arctic strains, and 13 assemblies of organisms belonging to the microbiome of (sub)arctic Cyanobacteria.

## Author contributions

LC and DB designed the experiments, LC performed all the computational analyses and drew the figures (except analyses and figures of Supplemental Note 1, which were carried out by ARB), LC and DB wrote the manuscript, with the assistance of ARB, MH, AW and EJJ. All authors read and approved the final manuscript.

## Acknowledgment

We thank Yannick Lara (ULiege) for the cultivation of the BCCM/ULC strains, his help with DNA extraction and his insightful comments on preliminary versions of the present article. Prof. Warwick Vincent (Laval University) is acknowledged for the deposit of strains ULC065, ULC066, ULC68, ULC077, ULC082 and ULC084.

---

**Table S1: Details and download links about reference proteomes**. All details were extracted from NCBI metadata.

**Table S2: Assembly statistics, taxonomy, completeness, contamination and coverage of genome bins.** The taxonomic label (CheckM taxon), the genome completeness (CM compl.), the contamination level (CM contam. and CM str. het.) were computed with CheckM, whereas DBX columns (DBX Cyano., DBX contam., DBX unknown, DBX unclass.) were computed by our DIAMOND blastx parser. Sequencing coverage [Coverage (med) and Coverage (IQR)] was computed with BBMap, while other statistics were extracted from QUAST output. Genome bins used in phylogenetic inference are marked by an (*) and discarded bins by an (-).

