## Supplementary Materials for "Metagenomic assembly of new (sub)arctic Cyanobacteria and their associated microbiome from non-axenic cultures"

| Assembly | Bioproject | Taxid | Name | Date of publication |
|---|---|---|---|---|
| GCA_000484535.1 | PRJNA162637 | 1183438 | Gloeobacter kilaueensis JS1 | 2013/10/28 |
| GCF_000011385.1 | PRJNA58011 | 251221 | Gloeobacter violaceus PCC 7421 | 2003/10/03 |
| GCF_000013205.1 | PRJNA224116 | 321327 | Synechococcus sp. JA-3-3Ab | 2006/02/06 |
| GCF_000013225.1 | PRJNA224116 | 321332 | Synechococcus sp. JA-2-3B'a(2-13) | 2006/02/06 |
| GCF_000332275.1 | PRJNA224116 | 195250 | Synechococcus sp. PCC 7336 | 2013/01/22 |
| GCF_000317065.1 | PRJNA224116 | 82654 | Pseudanabaena sp. PCC 7367 | 2012/12/05 |
| GCF_000332215.1 | PRJNA224116 | 927668 | Pseudanabaena biceps PCC 7429 | 2013/01/18 |
| GCF_000317085.1 | PRJNA224116 | 1173263 | Synechococcus sp. PCC 7502 | 2012/12/05 |
| GCF_000332175.1 | PRJNA224116 | 118173 | Pseudanabaena sp. PCC 6802 | 2013/01/22 |
| GCF_000018105.1 | PRJNA224116 | 329726 | Acaryochloris marina MBIC11017 | 2007/10/16 |
| GCA_000022045.1 | PRJNA28337 | 395961 | Cyanothece sp. PCC 7425 | 2009/01/09 |
| GCF_000505665.1 | PRJNA224116 | 1394889 | Thermosynechococcus sp. NK55a | 2013/12/11 |
| GCF_000316685.1 | PRJNA224116 | 195253 | Synechococcus sp. PCC 6312 | 2012/12/04 |
| GCF_000775285.1 | PRJNA224116 | 1497020 | Neosynechococcus sphagnicola sy1 | 2014/09/29 |
| GCF_000309945.1 | PRJNA224116 | 864702 | Oscillatoriales cyanobacterium JSC-12 | 2012/11/05 |
| GCF_001895925.1 | PRJNA224116 | 1920490 | Phormidesmis priestleyi ULC007 | 2016/12/09 |
| GCF_001650195.1 | PRJNA224116 | 1850361 | Phormidesmis priestleyi BC1401 | 2016/05/23 |
| GCF_000353285.1 | PRJNA224116 | 272134 | Leptolyngbya boryana PCC 6306 | 2013/04/06 |
| GCF_000733415.1 | PRJNA224116 | 1487953 | Leptolyngbya sp. JSC-1 | 2014/07/25 |
| GCF_000332095.2 | PRJNA224116 | 1173264 | Leptolyngbya sp. PCC 6406 | 2014/01/15 |
| GCF_000763385.1 | PRJNA224116 | 1229172 | Leptolyngbya sp. KIOST-1 | 2014/10/03 |
| GCF_000309385.1 | PRJNA224116 | 118166 | Nodosilinea nodulosa PCC 7104 | 2012/11/05 |
| GCF_000155595.1 | PRJNA224116 | 91464 | Synechococcus sp. PCC 7335 | 2008/08/15 |
| GCF_000482245.1 | PRJNA224116 | 1385935 | Leptolyngbya sp. Heron Island J | 2013/10/25 |
| GCF_000316115.1 | PRJNA224116 | 102129 | Leptolyngbya sp. PCC 7375 | 2012/12/03 |
| GCF_000464785.1 | PRJNA224116 | 1255374 | Planktothrix rubescens NIVA-CYA 407 | 2013/09/06 |
| GCF_000175415.3 | PRJNA224116 | 634502 | Arthrospira platensis str. Paraca | 2014/06/06 |
| GCF_000478195.2 | PRJNA224116 | 1348334 | Lyngbya aestuarii BL J | 2013/10/21 |
| GCF_000332155.1 | PRJNA224116 | 402777 | Kamptonema formosum PCC 6407 | 2013/01/22 |
| GCF_000317475.1 | PRJNA224116 | 179408 | Oscillatoria nigro-viridis PCC 7112 | 2012/12/06 |
| GCF_000317105.1 | PRJNA224116 | 56110 | Oscillatoria acuminata PCC 6304 | 2012/12/05 |
| GCF_000317515.1 | PRJNA224116 | 1173027 | Microcoleus sp. PCC 7113 | 2012/12/06 |
| GCF_000021825.1 | PRJNA224116 | 65393 | Cyanothece sp. PCC 7424 | 2008/12/17 |
| GCA_000307995.2 | PRJEA88171 | 1160280 | Microcystis aeruginosa PCC 9432 | 2012/08/06 |
| GCF_000021805.1 | PRJNA224116 | 41431 | Cyanothece sp. PCC 8801 | 2008/12/17 |
| GCF_000737945.1 | PRJNA256120 | 1527444 | Candidatus Atelocyanobacterium thalassa isolate SIO64986 | 2014/08/05 |
| GCF_000284135.1 | PRJNA224116 | 1080228 | Synechocystis sp. PCC 6803 substr. GT-I | 2011/12/01 |
| GCF_000715475.1 | PRJNA224116 | 490193 | Synechococcus sp. NKBG042902 | 2014/06/27 |
| GCF_000317655.1 | PRJNA39697 | 292563 | Cyanobacterium stanieri PCC 7202 | 2012/12/07 |
| GCF_000332055.1 | PRJNA224116 | 102125 | Xenococcus sp. PCC 7305 | 2013/01/17 |
| GCF_000317575.1 | PRJNA224116 | 111780 | Stanieria cyanosphaera PCC 7437 | 2012/12/06 |
| GCF_000380225.1 | PRJNA224116 | 1128427 | filamentous cyanobacterium ESFC-1 | 2013/04/23 |
| GCF_000317615.1 | PRJNA224116 | 13035 | Dactylococcopsis salina PCC 8305 | 2012/12/07 |
| GCF_000317495.1 | PRJNA224116 | 1173022 | Crinalium epipsammum PCC 9333 | 2012/12/06 |
| GCF_000317555.1 | PRJNA224116 | 1173026 | Gloeocapsa sp. PCC 7428 | 2012/12/06 |
| GCF_000317125.1 | PRJNA224116 | 251229 | Chroococcidiopsis thermalis PCC 7203 | 2012/12/05 |
| GCF_000582685.1 | PRJNA224116 | 1469607 | [Scytonema hofmanni] UTEX 2349 | 2014/02/27 |
| GCF_000789435.1 | PRJNA224116 | 1532906 | Aphanizomenon flos-aquae 2012/KM1/D3 | 2014/12/04 |
| GCF_000196515.1 | PRJNA224116 | 551115 | 'Nostoc azollae' 0708 | 2010/06/14 |
| GCF_000316645.1 | PRJNA224116 | 28072 | Nostoc sp. PCC 7524 | 2012/12/04 |
| GCF_000204075.1 | PRJNA10642 | 240292 | Anabaena variabilis ATCC 29413 | 2005/09/15 |
| GCA_000340565.3 | PRJNA185469 | 313624 | Nodularia spumigena CCY9414 | 2014/04/03 |
| GCF_000020025.1 | PRJNA224116 | 63737 | Nostoc punctiforme PCC 73102 | 2008/04/24 |
| GCF_000332295.1 | PRJNA224116 | 643473 | Fortiea contorta PCC 7126 | 2013/01/22 |
| GCF_000346485.2 | PRJNA224116 | 128403 | Scytonema hofmannii PCC 7110 | 2016/03/16 |
| GCF_000734895.2 | PRJNA224116 | 1337936 | Calothrix sp. 336/3 | 2015/05/06 |
| GCF_000332255.1 | PRJNA224116 | 1173021 | cyanobacterium PCC 7702 | 2013/01/22 |
| GCF_000317225.1 | PRJNA224116 | 98439 | Fischerella thermalis PCC 7521 | 2012/12/05 |
| GCF_000012525.1 | PRJNA224116 | 1140 | Synechococcus elongatus PCC 7942 | 2005/11/08 |
| GCF_000586015.1 | PRJNA224116 | 1451353 | Candidatus Synechococcus spongiarum SH4 | 2014/03/07 |
| GCF_000155635.1 | PRJNA224116 | 180281 | Cyanobium sp. PCC 7001 | 2008/08/29 |
| GCA_000015705.1 | PRJNA13496 | 59922 | Prochlorococcus marinus str. MIT 9303 | 2007/01/22 |
| GCF_000011485.1 | PRJNA224116 | 74547 | Prochlorococcus marinus str. MIT 9313 | 2003/08/18 |
| GCF_000153805.1 | PRJNA224116 | 313625 | Synechococcus sp. BL107 | 2006/10/23 |

| Assembly | ftp link |
|---|---|
| GCA_000484535.1 | ftp://ftp.ncbi.nlm.nih.gov/genomes/all/GCA/000/484/535/GCA_000484535.1_ASM48453v1 |
| GCF_000011385.1 | ftp://ftp.ncbi.nlm.nih.gov/genomes/all/GCF/000/011/385/GCF_000011385.1_ASM1138v1 |
| GCF_000013205.1 | ftp://ftp.ncbi.nlm.nih.gov/genomes/all/GCF/000/013/205/GCF_000013205.1_ASM1320v1 |
| GCF_000013225.1 | ftp://ftp.ncbi.nlm.nih.gov/genomes/all/GCF/000/013/225/GCF_000013225.1_ASM1322v1 |
| GCF_000332275.1 | ftp://ftp.ncbi.nlm.nih.gov/genomes/all/GCF/000/332/275/GCF_000332275.1_ASM33227v1 |
| GCF_000317065.1 | ftp://ftp.ncbi.nlm.nih.gov/genomes/all/GCF/000/317/065/GCF_000317065.1_ASM31706v1 |
| GCF_000332215.1 | ftp://ftp.ncbi.nlm.nih.gov/genomes/all/GCF/000/332/215/GCF_000332215.1_ASM33221v1 |
| GCF_000317085.1 | ftp://ftp.ncbi.nlm.nih.gov/genomes/all/GCF/000/317/085/GCF_000317085.1_ASM31708v1 |
| GCF_000332175.1 | ftp://ftp.ncbi.nlm.nih.gov/genomes/all/GCF/000/332/175/GCF_000332175.1_ASM33217v1 |
| GCF_000018105.1 | ftp://ftp.ncbi.nlm.nih.gov/genomes/all/GCF/000/018/105/GCF_000018105.1_ASM1810v1 |
| GCA_000022045.1 | ftp://ftp.ncbi.nlm.nih.gov/genomes/all/GCA/000/022/045/GCA_000022045.1_ASM2204v1 |
| GCF_000505665.1 | ftp://ftp.ncbi.nlm.nih.gov/genomes/all/GCF/000/505/665/GCF_000505665.1_ASM50566v1 |
| GCF_000316685.1 | ftp://ftp.ncbi.nlm.nih.gov/genomes/all/GCF/000/316/685/GCF_000316685.1_ASM31668v1 |
| GCF_000775285.1 | ftp://ftp.ncbi.nlm.nih.gov/genomes/all/GCF/000/775/285/GCF_000775285.1_ASM77528v1 |
| GCF_000309945.1 | ftp://ftp.ncbi.nlm.nih.gov/genomes/all/GCF/000/309/945/GCF_000309945.1_ASM30994v1 |
| GCF_001895925.1 | ftp://ftp.ncbi.nlm.nih.gov/genomes/all/GCF/001/895/925/GCF_001895925.1_ASM189592v1 |
| GCF_001650195.1 | ftp://ftp.ncbi.nlm.nih.gov/genomes/all/GCF/001/650/195/GCF_001650195.1_ASM165019v1 |
| GCF_000353285.1 | ftp://ftp.ncbi.nlm.nih.gov/genomes/all/GCF/000/353/285/GCF_000353285.1_ASM35328v1 |
| GCF_000733415.1 | ftp://ftp.ncbi.nlm.nih.gov/genomes/all/GCF/000/733/415/GCF_000733415.1_ASM73341v1 |
| GCF_000332095.2 | ftp://ftp.ncbi.nlm.nih.gov/genomes/all/GCF/000/332/095/GCF_000332095.2_ASM33209v2 |
| GCF_000763385.1 | ftp://ftp.ncbi.nlm.nih.gov/genomes/all/GCF/000/763/385/GCF_000763385.1_ASM76338v1 |
| GCF_000309385.1 | ftp://ftp.ncbi.nlm.nih.gov/genomes/all/GCF/000/309/385/GCF_000309385.1_ASM30938v1 |
| GCF_000155595.1 | ftp://ftp.ncbi.nlm.nih.gov/genomes/all/GCF/000/155/595/GCF_000155595.1_ASM15559v1 |
| GCF_000482245.1 | ftp://ftp.ncbi.nlm.nih.gov/genomes/all/GCF/000/482/245/GCF_000482245.1_LepHIscaffolds |
| GCF_000316115.1 | ftp://ftp.ncbi.nlm.nih.gov/genomes/all/GCF/000/316/115/GCF_000316115.1_ASM31611v1 |
| GCF_000464785.1 | ftp://ftp.ncbi.nlm.nih.gov/genomes/all/GCF/000/464/785/GCF_000464785.1_NC407_1 |
| GCF_000175415.3 | ftp://ftp.ncbi.nlm.nih.gov/genomes/all/GCF/000/175/415/GCF_000175415.3_ASM17541v3 |
| GCF_000478195.2 | ftp://ftp.ncbi.nlm.nih.gov/genomes/all/GCF/000/478/195/GCF_000478195.2_ASM47819v2 |
| GCF_000332155.1 | ftp://ftp.ncbi.nlm.nih.gov/genomes/all/GCF/000/332/155/GCF_000332155.1_ASM33215v1 |
| GCF_000317475.1 | ftp://ftp.ncbi.nlm.nih.gov/genomes/all/GCF/000/317/475/GCF_000317475.1_ASM31747v1 |
| GCF_000317105.1 | ftp://ftp.ncbi.nlm.nih.gov/genomes/all/GCF/000/317/105/GCF_000317105.1_ASM31710v1 |
| GCF_000317515.1 | ftp://ftp.ncbi.nlm.nih.gov/genomes/all/GCF/000/317/515/GCF_000317515.1_ASM31751v1 |
| GCF_000021825.1 | ftp://ftp.ncbi.nlm.nih.gov/genomes/all/GCF/000/021/825/GCF_000021825.1_ASM2182v1 |
| GCA_000307995.2 | ftp://ftp.ncbi.nlm.nih.gov/genomes/all/GCA/000/307/995/GCA_000307995.2_ASM30799v2 |
| GCF_000021805.1 | ftp://ftp.ncbi.nlm.nih.gov/genomes/all/GCF/000/021/805/GCF_000021805.1_ASM2180v1 |
| GCF_000737945.1 | ftp://ftp.ncbi.nlm.nih.gov/genomes/all/GCF/000/737/945/GCF_000737945.1_ASM73794v1 |
| GCF_000284135.1 | ftp://ftp.ncbi.nlm.nih.gov/genomes/all/GCF/000/284/135/GCF_000284135.1_ASM28413v1 |
| GCF_000715475.1 | ftp://ftp.ncbi.nlm.nih.gov/genomes/all/GCF/000/715/475/GCF_000715475.1_ASM71547v1 |
| GCF_000317655.1 | ftp://ftp.ncbi.nlm.nih.gov/genomes/all/GCF/000/317/655/GCF_000317655.1_ASM31765v1 |
| GCF_000332055.1 | ftp://ftp.ncbi.nlm.nih.gov/genomes/all/GCF/000/332/055/GCF_000332055.1_ASM33205v1 |
| GCF_000317575.1 | ftp://ftp.ncbi.nlm.nih.gov/genomes/all/GCF/000/317/575/GCF_000317575.1_ASM31757v1 |
| GCF_000380225.1 | ftp://ftp.ncbi.nlm.nih.gov/genomes/all/GCF/000/380/225/GCF_000380225.1_ASM38022v1 |
| GCF_000317615.1 | ftp://ftp.ncbi.nlm.nih.gov/genomes/all/GCF/000/317/615/GCF_000317615.1_ASM31761v1 |
| GCF_000317495.1 | ftp://ftp.ncbi.nlm.nih.gov/genomes/all/GCF/000/317/495/GCF_000317495.1_ASM31749v1 |
| GCF_000317555.1 | ftp://ftp.ncbi.nlm.nih.gov/genomes/all/GCF/000/317/555/GCF_000317555.1_ASM31755v1 |
| GCF_000317125.1 | ftp://ftp.ncbi.nlm.nih.gov/genomes/all/GCF/000/317/125/GCF_000317125.1_ASM31712v1 |
| GCF_000582685.1 | ftp://ftp.ncbi.nlm.nih.gov/genomes/all/GCF/000/582/685/GCF_000582685.1_ASM58268v1 |
| GCF_000789435.1 | ftp://ftp.ncbi.nlm.nih.gov/genomes/all/GCF/000/789/435/GCF_000789435.1_ASM78943v1 |
| GCF_000196515.1 | ftp://ftp.ncbi.nlm.nih.gov/genomes/all/GCF/000/196/515/GCF_000196515.1_ASM19651v1 |
| GCF_000316645.1 | ftp://ftp.ncbi.nlm.nih.gov/genomes/all/GCF/000/316/645/GCF_000316645.1_ASM31664v1 |
| GCF_000204075.1 | ftp://ftp.ncbi.nlm.nih.gov/genomes/all/GCF/000/204/075/GCF_000204075.1_ASM20407v1 |
| GCA_000340565.3 | ftp://ftp.ncbi.nlm.nih.gov/genomes/all/GCA/000/340/565/GCA_000340565.3_ASM34056v3 |
| GCF_000020025.1 | ftp://ftp.ncbi.nlm.nih.gov/genomes/all/GCF/000/020/025/GCF_000020025.1_ASM2002v1 |
| GCF_000332295.1 | ftp://ftp.ncbi.nlm.nih.gov/genomes/all/GCF/000/332/295/GCF_000332295.1_ASM33229v1 |
| GCF_000346485.2 | ftp://ftp.ncbi.nlm.nih.gov/genomes/all/GCF/000/346/485/GCF_000346485.2_ASM34648v2 |
| GCF_000734895.2 | ftp://ftp.ncbi.nlm.nih.gov/genomes/all/GCF/000/734/895/GCF_000734895.2_ASM73489v2 |
| GCF_000332255.1 | ftp://ftp.ncbi.nlm.nih.gov/genomes/all/GCF/000/332/255/GCF_000332255.1_ASM33225v1 |
| GCF_000317225.1 | ftp://ftp.ncbi.nlm.nih.gov/genomes/all/GCF/000/317/225/GCF_000317225.1_FisPCC7521_1.0 |
| GCF_000012525.1 | ftp://ftp.ncbi.nlm.nih.gov/genomes/all/GCF/000/012/525/GCF_000012525.1_ASM1252v1 |
| GCF_000586015.1 | ftp://ftp.ncbi.nlm.nih.gov/genomes/all/GCF/000/586/015/GCF_000586015.1_SynSpo.0 |
| GCF_000155635.1 | ftp://ftp.ncbi.nlm.nih.gov/genomes/all/GCF/000/155/635/GCF_000155635.1_ASM15563v1 |
| GCA_000015705.1 | ftp://ftp.ncbi.nlm.nih.gov/genomes/all/GCA/000/015/705/GCA_000015705.1_ASM1570v1 |
| GCF_000011485.1 | ftp://ftp.ncbi.nlm.nih.gov/genomes/all/GCF/000/011/485/GCF_000011485.1_ASM1148v1 |
| GCF_000153805.1 | ftp://ftp.ncbi.nlm.nih.gov/genomes/all/GCF/000/153/805/GCF_000153805.1_ASM15380v1 |

| Strain | MetaBAT setting | Bin | CheckM taxon | #Scaffolds | #Contigs | #C. (>1000 nt) | Length (nt) | Length (%) | L. (>1000 nt) | GC (%) | L50 | N75 (nt) |
|---|---|---|---|---|---|---|---|---|---|---|---|---|
| ULC335 | veryspecific | 1 | Cyanobacteria* | 238 | 306 | 294 | 4602507 | 20.84 | 4595893 | 40.74 | 40.75 | 24097 |
| | | 2 | Flavobacteriaceae* | 67 | 114 | 109 | 3031816 | 13.73 | 3027871 | 32.72 | 32.74 | 54021 |
| | | 3 | Bacteroidetes* | 576 | 1335 | 938 | 2834244 | 12.83 | 2592321 | 45.95 | 45.96 | 3591 |
| | | 4 | Alphaproteobacteria* | 271 | 645 | 387 | 1056769 | 4.79 | 905578 | 67.05 | 67.06 | 3052 |
| | | 0 | nobin | 23056 | 24863 | 1517 | 10558473 | 47.81 | 2338854 | 61.30 | 52.35 | 2306 |
| ULC007 | superspecific | 1 | Cyanobacteria* | 84 | 106 | 106 | 5223811 | 91.14 | 5223491 | 48.62 | 48.63 | 72864 |
| | | 2 | Unclassified | 12 | 17 | 16 | 283907 | 4.95 | 282902 | 48.48 | 48.54 | 22102 |
| | | 0 | nobin | 358 | 358 | 49 | 224030 | 3.91 | 157489 | 48.69 | 48.51 | 6599 |
| ULC027 | verysensitive | 1 | Cyanobacteria* | 439 | 1018 | 870 | 4766714 | 21.40 | 4673662 | 50.35 | 50.4 | 7298 |
| | | 2 | Alphaproteobacteria* | 190 | 553 | 471 | 3599900 | 16.16 | 3549598 | 63.22 | 63.25 | 11716 |
| | | 3 | Sphingomonadales* | 293 | 782 | 599 | 2678920 | 12.03 | 2571356 | 62.95 | 62.97 | 5694 |
| | | 4 | Unclassified | 164 | 414 | 280 | 927349 | 4.16 | 850288 | 65.33 | 65.69 | 4292 |
| | | 0 | nobin | 24364 | 26287 | 1032 | 10298357 | 46.24 | 1737206 | 60.46 | 52.73 | 2732 |
| ULC041 | verysensitive | 1 | Cyanobacteria* | 287 | 319 | 313 | 4697657 | 84.76 | 4694951 | 58.23 | 58.23 | 21370 |
| | | 2 | Unclassified | 24 | 24 | 24 | 518635 | 9.36 | 518635 | 51.50 | 51.5 | 33120 |
| | | 0 | nobin | 441 | 444 | 68 | 325698 | 5.88 | 256390 | 54.55 | 53.23 | 8350 |
| ULC065 | veryspecific | 1 | Cyanobacteria* | 95 | 119 | 117 | 3082321 | 22.36 | 3080895 | 66.03 | 66.03 | 38827 |
| | | 2 | Xanthomonadaceae* | 332 | 767 | 601 | 2664261 | 19.33 | 2565831 | 67.81 | 67.84 | 5855 |
| | | 0 | nobin | 20555 | 21891 | 649 | 8036576 | 58.31 | 1004543 | 63.22 | 62.66 | 2548 |
| ULC066 | superspecific | 1 | Cyanobacteria* | 67 | 86 | 86 | 5380362 | 28.81 | 5379882 | 41.84 | 41.84 | 99258 |
| | | 2 | Bacteroidetes* | 401 | 904 | 670 | 2603362 | 13.94 | 2457932 | 39.33 | 39.53 | 4764 |
| | | 3 | Betaproteobacteria- | 152 | 367 | 219 | 534032 | 2.86 | 442359 | 52.02 | 52.15 | 2784 |
| | | 0 | nobin | 24558 | 26258 | 1225 | 10154658 | 54.38 | 1892140 | 58.57 | 48.27 | 2394 |
| ULC068 | superspecific | 1 | Cyanobacteria* | 60 | 84 | 83 | 4551587 | 57.04 | 4550868 | 42.22 | 42.23 | 80830 |
| | | 2 | Unclassified | 3 | 4 | 4 | 203969 | 2.56 | 203969 | 42.75 | 42.75 | 61745 |
| | | 0 | nobin | 10385 | 10655 | 40 | 3224460 | 40.41 | 165266 | 62.14 | 43.04 | 103751 |
| ULC073 | verysensitive | 1 | Cyanobacteria* | 476 | 619 | 572 | 4603354 | 22.70 | 4578777 | 57.76 | 57.74 | 10574 |
| | | 2 | Betaproteobacteria* | 65 | 237 | 224 | 3297508 | 16.26 | 3287197 | 52.41 | 52.44 | 25326 |
| | | 3 | Sphingomonadales* | 603 | 1581 | 1009 | 3199566 | 15.78 | 2861160 | 63.66 | 63.78 | 3639 |
| | | 4 | Bacteria- | 156 | 372 | 200 | 566613 | 2.79 | 462681 | 63.90 | 64.34 | 2882 |
| | | 5 | Unclassified | 26 | 27 | 27 | 283727 | 1.40 | 283717 | 49.79 | 49.79 | 13765 |
| | | 6 | Unclassified | 29 | 70 | 58 | 279658 | 1.38 | 273663 | 57.84 | 57.96 | 6091 |
| | | 0 | nobin | 16790 | 18486 | 1325 | 8046800 | 39.68 | 2063350 | 62.84 | 58.35 | 2376 |
| ULC077 | veryspecific | 1 | Cyanobacteria* | 407 | 451 | 446 | 5440088 | 47.37 | 5436761 | 48.17 | 48.17 | 17549 |
| | | 0 | nobin | 14903 | 16237 | 658 | 6044297 | 52.63 | 1134577 | 56.66 | 50.09 | 3489 |
| ULC082 | veryspecific | 1 | Cyanobacteria* | 124 | 148 | 145 | 2790347 | 11.49 | 2788897 | 65.29 | 65.31 | 37235 |
| | | 2 | Bacteria* | 529 | 1318 | 842 | 2371940 | 9.77 | 2087863 | 63.55 | 63.6 | 3235 |
| | | 3 | Bacteria* | 542 | 1266 | 756 | 1981145 | 8.16 | 1666752 | 59.58 | 59.71 | 2939 |
| | | 4 | Bacteria- | 120 | 260 | 158 | 417999 | 1.72 | 357553 | 66.70 | 66.9 | 2961 |
| | | 5 | Unclassified | 74 | 172 | 125 | 405790 | 1.67 | 377521 | 53.86 | 53.85 | 3737 |
| | | 0 | nobin | 30077 | 33680 | 3169 | 16310948 | 67.18 | 4763602 | 63.05 | 59.77 | 2314 |
| ULC084 | superspecific | 1 | Betaproteobacteria* | 232 | 718 | 585 | 2976717 | 23.15 | 2892002 | 52.52 | 52.63 | 6715 |
| | | 2 | Alphaproteobacteria* | 222 | 573 | 482 | 2878549 | 22.39 | 2826872 | 64.33 | 64.34 | 8706 |
| | | 3 | Cyanobacteria* | 116 | 139 | 129 | 2813103 | 21.88 | 2809623 | 65.45 | 65.46 | 33348 |
| | | 0 | nobin | 10835 | 11315 | 276 | 4188263 | 32.58 | 775675 | 59.91 | 53.48 | 6615 |
| ULC129 | verysensitive | 1 | Cyanobacteria* | 299 | 317 | 316 | 4724332 | 38.35 | 4723951 | 51.02 | 51.02 | 20681 |
| | | 0 | nobin | 21968 | 22744 | 305 | 7596258 | 61.65 | 588929 | 53.45 | 49.42 | 4773 |
| ULC146 | superspecific | 1 | Burkholderiales* | 177 | 347 | 319 | 4889081 | 16.18 | 4871892 | 64.90 | 64.91 | 24186 |
| | | 2 | Flavobacteriaceae* | 285 | 679 | 592 | 3900875 | 12.91 | 3842453 | 37.74 | 37.87 | 8727 |
| | | 3 | Sphingomonadales* | 74 | 133 | 125 | 3487715 | 11.54 | 3483390 | 63.94 | 63.95 | 43449 |
| | | 4 | Betaproteobacteria* | 98 | 289 | 258 | 3278028 | 10.85 | 3257798 | 52.43 | 52.46 | 19528 |
| | | 5 | Alphaproteobacteria* | 350 | 771 | 582 | 2282825 | 7.56 | 2172115 | 66.42 | 66.53 | 5042 |
| | | 6 | Bacteria- | 243 | 550 | 336 | 940367 | 3.11 | 810232 | 61.75 | 61.77 | 3017 |
| | | 7 | Unclassified | 21 | 30 | 29 | 561075 | 1.86 | 560413 | 65.64 | 65.64 | 29757 |
| | | 0 | nobin | 28569 | 29987 | 976 | 10872327 | 35.99 | 1638657 | 57.78 | 56.11 | 2467 |
| ULC165 | verysensitive | 1 | Xanthomonadaceae* | 53 | 72 | 66 | 3099833 | 15.37 | 3097278 | 68.28 | 68.28 | 83934 |
| | | 2 | Alphaproteobacteria* | 167 | 409 | 367 | 2927747 | 14.52 | 2903874 | 64.53 | 64.56 | 11367 |
| | | 3 | Burkholderiales* | 473 | 1056 | 681 | 2018813 | 10.01 | 1786411 | 63.70 | 63.8 | 3287 |
| | | 4 | Bacteria* | 356 | 674 | 471 | 1270968 | 6.30 | 1141258 | 47.34 | 47.65 | 3006 |
| | | 0 | nobin | 19409 | 21418 | 2321 | 10846889 | 53.79 | 3733515 | 57.99 | 53.8 | 2416 |
| ULC179 | superspecific | 1 | Alphaproteobacteria* | 247 | 359 | 344 | 5293532 | 18.89 | 5284967 | 66.86 | 66.87 | 23221 |
| | | 2 | Rhizobiales* | 261 | 671 | 577 | 4749544 | 16.95 | 4692884 | 65.61 | 65.66 | 12479 |
| | | 3 | Alphaproteobacteria* | 111 | 162 | 154 | 3817570 | 13.62 | 3813929 | 67.71 | 67.72 | 39185 |
| | | 4 | Cytophagales* | 718 | 1658 | 1180 | 3753883 | 13.40 | 3454758 | 51.71 | 51.91 | 3772 |
| | | 5 | Alphaproteobacteria* | 68 | 95 | 89 | 1318133 | 4.70 | 1314626 | 64.79 | 64.79 | 24810 |
| | | 6 | Rhizobiales* | 170 | 391 | 231 | 604053 | 2.16 | 504692 | 61.84 | 62.25 | 2997 |
| | | 7 | Unclassified | 16 | 22 | 22 | 473840 | 1.69 | 473830 | 61.96 | 61.96 | 30325 |
| | | 0 | nobin | 13101 | 14683 | 1888 | 8010556 | 28.59 | 3418733 | 57.66 | 54.68 | 3032 |
| ULC186 | verysensitive | 1 | Cyanobacteria* | 412 | 441 | 433 | 5062396 | 67.38 | 5057101 | 57.41 | 57.41 | 16589 |
| | | 0 | nobin | 6559 | 6646 | 190 | 2450340 | 32.62 | 628243 | 50.93 | 53.77 | 9233 |
| ULC187 | veryspecific | 1 | Cyanobacteria* | 62 | 71 | 71 | 4461408 | 62.18 | 4461288 | 43.15 | 43.15 | 115626 |
| | | 0 | nobin | 8482 | 8566 | 40 | 2713800 | 37.82 | 234659 | 59.70 | 43.51 | 40647 |

| Strain | MetaBAT setting | Bin | CheckM taxon | Coverage (med) | Coverage (IQR) | CM compl. | CM contam. | CM str. het. | DBX Cyano. | DBX contam. |
|---|---|---|---|---|---|---|---|---|---|---|
| ULC335 | veryspecific | 1 | Cyanobacteria* | 10.90 | 1.02 | 98.91 | 0.51 | 0 | 57.00 | 1.26 |
| | | 2 | Flavobacteriaceae* | 11.12 | 1.37 | 99.29 | 0.12 | 0 | 0.06 | 69.60 |
| | | 3 | Bacteroidetes* | 4.46 | 0.68 | 65.45 | 0.49 | 0 | 0.23 | 44.02 |
| | | 4 | Alphaproteobacteria* | 4.13 | 0.75 | 32.28 | 0 | 0 | 0.14 | 58.32 |
| | | 0 | nobin | 1.88 | 1.10 | NA | NA | NA | 0.63 | 46.97 |
| ULC007 | superspecific | 1 | Cyanobacteria* | 26.62 | 1.26 | 98.11 | 0 | 0 | 58.12 | 0.71 |
| | | 2 | Unclassified | 72.12 | 4.38 | 0 | 0 | 0 | 42.82 | 3.06 |
| | | 0 | nobin | 1.48 | 24.50 | NA | NA | NA | 31.18 | 2.51 |
| ULC027 | verysensitive | 1 | Cyanobacteria* | 6.27 | 1.24 | 90.43 | 0.27 | 0 | 52.26 | 1.02 |
| | | 2 | Alphaproteobacteria* | 7.71 | 1.37 | 95.02 | 1.16 | 0 | 0.36 | 36.58 |
| | | 3 | Sphingomonadales* | 6.18 | 1.20 | 60.21 | 2.35 | 7.14 | 0.13 | 58.84 |
| | | 4 | Unclassified | 5.09 | 0.99 | 4.17 | 0 | 0 | 0.00 | 58.47 |
| | | 0 | nobin | 1.89 | 1.11 | NA | NA | NA | 2.61 | 47.76 |
| ULC041 | verysensitive | 1 | Cyanobacteria* | 31.38 | 7.36 | 96.2 | 1.63 | 22.22 | 48.68 | 1.30 |
| | | 2 | Unclassified | 44.33 | 13.47 | 0 | 0 | 0 | 38.64 | 2.73 |
| | | 0 | nobin | 3.97 | 31.72 | NA | NA | NA | 26.86 | 1.86 |
| ULC065 | veryspecific | 1 | Cyanobacteria* | 38.37 | 10.53 | 99.09 | 0.27 | 0 | 54.48 | 1.34 |
| | | 2 | Xanthomonadaceae* | 6.19 | 1.32 | 83.73 | 1.23 | 0 | 0.06 | 68.62 |
| | | 0 | nobin | 1.73 | 0.88 | NA | NA | NA | 0.27 | 49.47 |
| ULC066 | superspecific | 1 | Cyanobacteria* | 21.86 | 2.26 | 98.82 | 0.47 | 50 | 64.79 | 0.97 |
| | | 2 | Bacteroidetes* | 4.93 | 0.75 | 76.91 | 1.23 | 0 | 0.32 | 53.00 |
| | | 3 | Betaproteobacteria- | 3.48 | 0.54 | 15.86 | 0 | 0 | 0.00 | 74.50 |
| | | 0 | nobin | 1.69 | 0.90 | NA | NA | NA | 0.33 | 42.49 |
| ULC068 | superspecific | 1 | Cyanobacteria* | 29.34 | 0.80 | 97.09 | 0.71 | 0 | 64.09 | 1.37 |
| | | 2 | Unclassified | 22.60 | 1.32 | 0 | 0 | 0 | 54.52 | 2.69 |
| | | 0 | nobin | 1.42 | 0.62 | NA | NA | NA | 1.67 | 42.78 |
| ULC073 | verysensitive | 1 | Cyanobacteria* | 10.74 | 1.99 | 92.03 | 1.42 | 12.5 | 48.29 | 1.45 |
| | | 2 | Betaproteobacteria* | 7.99 | 0.54 | 97.92 | 0.67 | 0 | 0.02 | 73.70 |
| | | 3 | Sphingomonadales* | 4.94 | 1.21 | 70.57 | 5.3 | 5.41 | 0.05 | 60.55 |
| | | 4 | Bacteria- | 4.39 | 1.59 | 10.71 | 0 | 0 | 0.00 | 69.36 |
| | | 5 | Unclassified | 15.02 | 2.12 | 0 | 0 | 0 | 39.22 | 4.96 |
| | | 6 | Unclassified | 6.45 | 1.52 | 0 | 0 | 0 | 0.62 | 46.64 |
| | | 0 | nobin | 1.94 | 1.37 | NA | NA | NA | 1.80 | 51.59 |
| ULC077 | veryspecific | 1 | Cyanobacteria* | 15.08 | 2.30 | 97.64 | 0.47 | 0 | 57.21 | 1.00 |
| | | 0 | nobin | 1.83 | 1.16 | NA | NA | NA | 3.62 | 44.61 |
| ULC082 | veryspecific | 1 | Cyanobacteria* | 19.85 | 5.24 | 97.74 | 0.27 | 50 | 57.40 | 1.84 |
| | | 2 | Bacteria* | 4.50 | 1.00 | 62.77 | 7.54 | 4.76 | 0.04 | 58.67 |
| | | 3 | Bacteria* | 3.88 | 0.70 | 46.21 | 9.28 | 0 | 0.00 | 80.26 |
| | | 4 | Bacteria- | 4.73 | 1.44 | 11.13 | 0 | 0 | 0.12 | 63.46 |
| | | 5 | Unclassified | 4.57 | 0.64 | 0 | 0 | 0 | 0.00 | 70.78 |
| | | 0 | nobin | 2.15 | 1.32 | NA | NA | NA | 0.28 | 55.82 |
| ULC084 | superspecific | 1 | Betaproteobacteria* | 5.67 | 0.69 | 93.61 | 1.73 | 0 | 0.06 | 74.26 |
| | | 2 | Alphaproteobacteria* | 6.65 | 1.23 | 92.46 | 1.38 | 0 | 0.03 | 67.26 |
| | | 3 | Cyanobacteria* | 20.78 | 5.58 | 98.55 | 0 | 0 | 57.26 | 2.63 |
| | | 0 | nobin | 1.59 | 0.74 | NA | NA | NA | 1.33 | 43.66 |
| ULC129 | verysensitive | 1 | Cyanobacteria* | 18.46 | 2.64 | 98.64 | 0.77 | 25 | 53.27 | 1.03 |
| | | 0 | nobin | 1.62 | 0.74 | NA | NA | NA | 1.83 | 36.96 |
| ULC146 | superspecific | 1 | Burkholderiales* | 10.96 | 1.97 | 96.57 | 0.93 | 0 | 0.05 | 68.54 |
| | | 2 | Flavobacteriaceae* | 6.27 | 0.88 | 94.94 | 0.35 | 0 | 0.04 | 65.50 |
| | | 3 | Sphingomonadales* | 14.23 | 1.93 | 88.9 | 1.39 | 14.29 | 0.10 | 60.31 |
| | | 4 | Betaproteobacteria* | 7.64 | 0.72 | 97.46 | 1.09 | 0 | 0.03 | 73.74 |
| | | 5 | Alphaproteobacteria* | 6.25 | 1.48 | 75.87 | 0.32 | 0 | 0.04 | 63.80 |
| | | 6 | Bacteria- | 4.68 | 1.62 | 10.82 | 0 | 0 | 0.13 | 63.90 |
| | | 7 | Unclassified | 12.53 | 1.26 | 8.33 | 0 | 0 | 0.04 | 60.31 |
| | | 0 | nobin | 1.72 | 0.93 | NA | NA | NA | 8.43 | 42.17 |
| ULC165 | verysensitive | 1 | Xanthomonadaceae* | 24.76 | 4.65 | 99.54 | 0.8 | 0 | 0.09 | 70.37 |
| | | 2 | Alphaproteobacteria* | 7.75 | 1.58 | 96.29 | 1.22 | 16.67 | 0.04 | 68.60 |
| | | 3 | Burkholderiales* | 4.40 | 1.05 | 41.41 | 0.47 | 0 | 0.04 | 66.06 |
| | | 4 | Bacteria* | 3.90 | 0.77 | 24.14 | 1.72 | 0 | 50.56 | 1.23 |
| | | 0 | nobin | 2.08 | 1.58 | NA | NA | NA | 15.26 | 29.20 |
| ULC179 | superspecific | 1 | Alphaproteobacteria* | 16.30 | 5.72 | 98.54 | 60.19 | 22.41 | 0.05 | 74.90 |
| | | 2 | Rhizobiales* | 8.86 | 2.08 | 94.78 | 0.94 | 0 | 0.00 | 65.89 |
| | | 3 | Alphaproteobacteria* | 21.92 | 5.67 | 98.73 | 0.22 | 0 | 0.10 | 48.99 |
| | | 4 | Cytophagales* | 4.60 | 0.77 | 67.06 | 0.3 | 0 | 0.11 | 68.27 |
| | | 5 | Alphaproteobacteria* | 16.67 | 6.03 | 35.78 | 0 | 0 | 0.00 | 63.92 |
| | | 6 | Rhizobiales* | 4.18 | 1.28 | 12.58 | 0 | 0 | 0.00 | 67.76 |
| | | 7 | Unclassified | 41.33 | 11.73 | 0 | 0 | 0 | 1.26 | 38.69 |
| | | 0 | nobin | 1.94 | 1.77 | NA | NA | NA | 5.12 | 48.73 |
| ULC186 | verysensitive | 1 | Cyanobacteria* | 21.10 | 4.81 | 93.18 | 1.64 | 9.09 | 47.80 | 1.49 |
| | | 0 | nobin | 1.52 | 0.76 | NA | NA | NA | 8.95 | 37.47 |
| ULC187 | veryspecific | 1 | Cyanobacteria* | 33.11 | 1.39 | 99.29 | 0.47 | 0 | 62.11 | 0.77 |
| | | 0 | nobin | 1.43 | 0.60 | NA | NA | NA | 2.98 | 41.52 |

### Supplemental Note 1

During the genome binning step with MetaBAT, we lost most of our SSU rRNA (16S) sequences. As far as we know, the causes for this common issue (Cornet et al., 2018 in review) have not yet been discussed in the metagenomic literature. In order to understand why those rRNA genes are discarded instead of being placed in their respective genome bin, we investigated the possible differences in sequencing coverage and tetranucleotide frequencies (TNFs) between the contigs bearing rRNA genes and the other contigs composing the MetaBAT genome bins.

#### Methods

For each contig, the sequencing coverage (totalAvgDepth) was extracted from the output files of MetaBAT (1). Coverage values were then grouped by genome bin and plotted as box-and-whiskers plots, themselves grouped by metagenome. The contigs bearing unbinned SSU rRNA (16S) sequences were all pooled in a single special pseudo-bin (r) for each metagenome.

TNFs were obtained for each contig with the program compseq [from the EMBOSS software package; (2)] using default settings. The resulting observed frequencies were used to perform 17 principal component analyses (PCAs; using the R function prcomp of the stats package; https://stat.ethz.ch/R-manual/R-devel/library/stats/html/prcomp.html), each one combining all the contigs of all genome bins from a single metagenome, including rRNA-bearing contigs, whether correctly binned by MetaBAT, manually affiliated by us or left unaffiliated (see main text). The two first components were then plotted for each metagenome.

For each contig bearing a rRNA sequence, the length was extracted from the output files of MetaBAT. rRNA-bearing contig lengths were then grouped in two categories (binned or unbinned) and plotted as box-and-whiskers plots. The same partition in two categories was used to compare modified Z-scores (3) measuring the deviation of the sequencing coverage of rRNA-bearing contigs with respect to the median coverage of the corresponding bins. Because of this requirement, rRNA genes left unaffiliated had to be excluded from the latter analysis.

#### Results and Discussion

Our working hypothesis was that the presence of multi-copy rRNA operons in Bacteria (4) would result in a higher sequencing coverage, which might interfere with MetaBAT binning. As shown in **Figure S1**, most contigs with rRNA genes that could not be automatically binned (in red) display markedly higher coverage values than “regular” genome bin contigs. In contrast, when the coverage of rRNA-bearing contigs is less extreme, they can be included in a genome bin (black arrowheads in

ULC027-bin2, ULC073-bin6, ULC146-bin1/bin3, and ULC179-bin3). When it is not the case, the problem stems from the divergent TNFs of rRNA genes. Indeed, the PCAs in **Figure S2** show that unbinned rRNA-bearing contigs (colored arrowheads) either cluster outside all bin clouds (ULC335-nobin) or cluster in a bin cloud for which the coverage is incompatible (ULC082-nobin, ULC179-nobin), whereas the five rRNA-bearing contigs that were automatically binned all lie in their respective bin cloud (black arrowheads). Inversely, there exist rRNA-bearing contigs that could not be binned because of their incompatible coverage only (e.g., ULC073-bin2, ULC084-bin1/bin2, ULC129-bin1), while a number could not be binned due to the incompatibility of both their coverage and TNFs (e.g., ULC007, ULC041, ULC068, ULC082-bin1, ULC146-bin2, ULC165, ULC187). Given that MetaBAT uses a distance combining the sequencing coverage and the TNFs to partition the contigs into distinct genome bins, it is not surprising that it discards the majority of rRNA-bearing contigs in light of these results.

To better understand the factors governing the binning fate of rRNA-bearing contigs, we compared their lengths. As shown in **Figure S3a**, binned contigs (in black) are considerably longer than unbinned contigs (in red). One possible cause for this difference might be that unbinned rRNA-bearing contigs exist in multiple copies in their respective genome. Such a situation would prevent the assembler software to extend the contigs much beyond the rRNA operon itself, because of the different genomic contexts surrounding the operon. Interestingly, unbinned rRNA-bearing contigs that we were able to manually affiliate a posteriori show a coverage significantly higher than the median coverage of their respective genome bin (**Figure S3b**), which suggests that they indeed do exist in multiple copies.

#### Conclusion

Our analyses suggest that the loss of rRNA sequences during the binning process carried out by MetaBAT is both driven by the higher sequencing coverage of these genes with respect to the coverage of other genome contigs and by their TNFs that are also markedly different. While these two factors matter, they are not completely independent. In particular, the divergent TNFs of rRNA genes are overwhelmed by the “regular” TNFs of the surrounding sequences when the rRNA operon is included in a long contig, which allows single-copy rRNA operons to be automatically binned. In contrast, multi-copy rRNA operons, which lie on shorter contigs, do not benefit from this dilution phenomenon. Consequently, their divergent TNFs add up to their higher sequencing coverage, resulting in the loss of rRNA genes. Altogether, automatic binning of rRNA genes appears to remain very challenging and would probably be improved by the use of longer sequencing reads, so as to assemble the multi-copy rRNA operons on longer contigs.

### Figures

**Figure S1: Comparison of the sequencing coverage of unbinned contigs bearing SSU rRNA (16S) genes to the coverage of other genome contigs grouped by genome bin and by metagenome.** Numbers on the X axis refer to the bin numbers in the main text (r = rRNA genes) and always follow the same color code: from the first to the last, blue, green, purple, orange, yellow, brown and pink. Stars denote cyanobacterial bins, while arrowheads indicate the coverage (and bins) of the rRNA-bearing contigs that were automatically binned by MetaBAT.

**Figure S2: Principal Component Analyses of tetranucleotide frequencies in all contigs for each metagenome.** Unbinned contigs bearing rRNA genes are colored in red. Each genome bin is colored, from the first to the last, in blue, green, purple, orange, yellow, brown and pink (as in **Figure S1**). Cyanobacterial bins were circled by hand using dashed lines. Arrowheads point to rRNA-bearing contigs, either automatically binned by MetaBAT (in black) or manually affiliated (colored after their MetaBAT bin) or left unaffiliated (in red).

**Figure S3: Comparison between binned and unbinned contigs bearing rRNA genes for a) their length and b) their deviation from the median sequencing coverage of their respective bin.** Numbers on top of box-and-whiskers are the number of contigs contributing to each box. Unbinned contigs used in **b)** are restricted to those that we were able to manually affiliate.

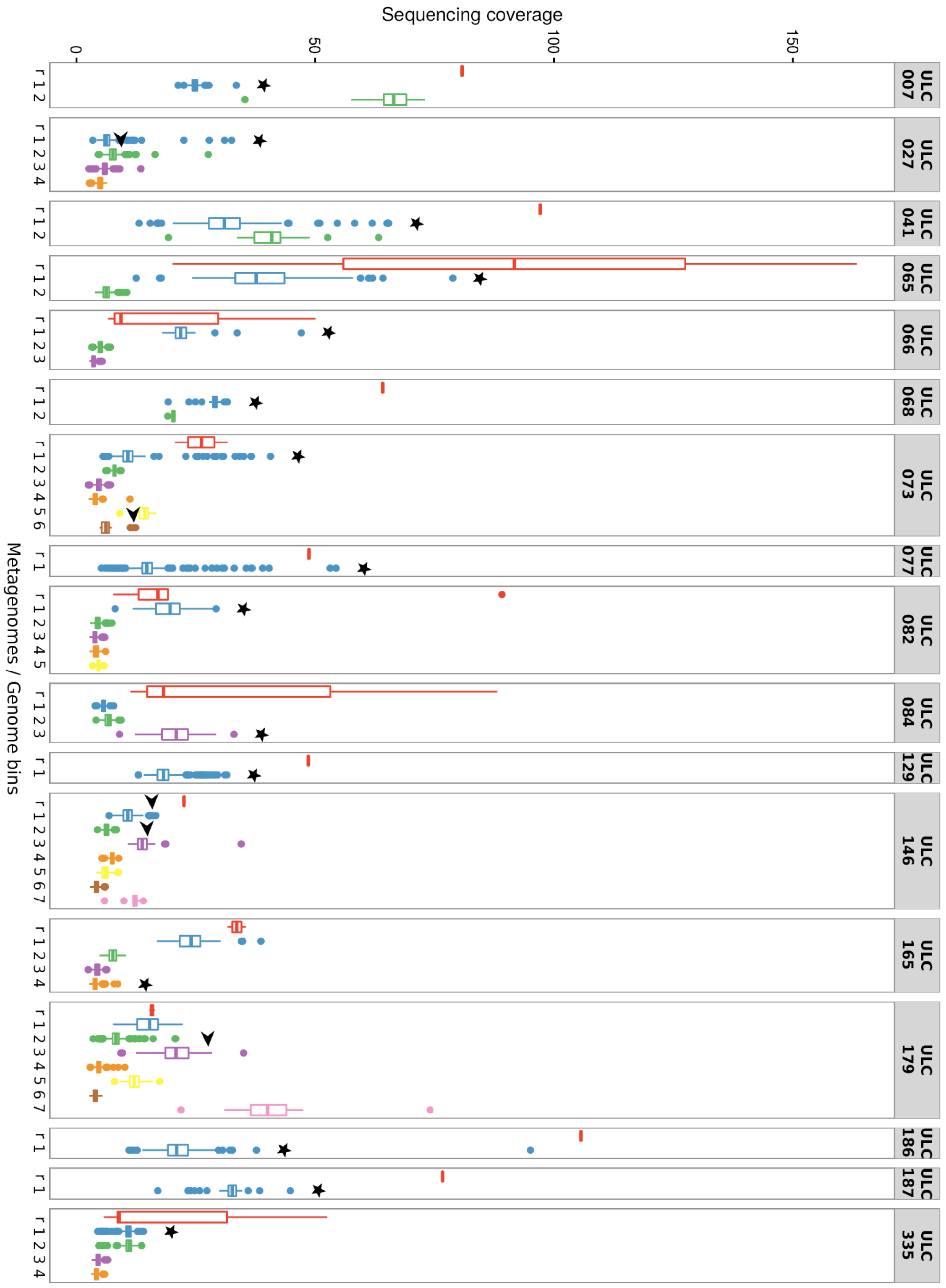
Sequencing coverage
0
50
100
150
Metagenomes / Genome bins
ULC 007
ULC 027
ULC 041
ULC 065
ULC 066
ULC 068
ULC 073
ULC 077
ULC 082
ULC 084
ULC 129
ULC 146
ULC 165
ULC 179
ULC 186
ULC 187
ULC 335

**Figure S1**

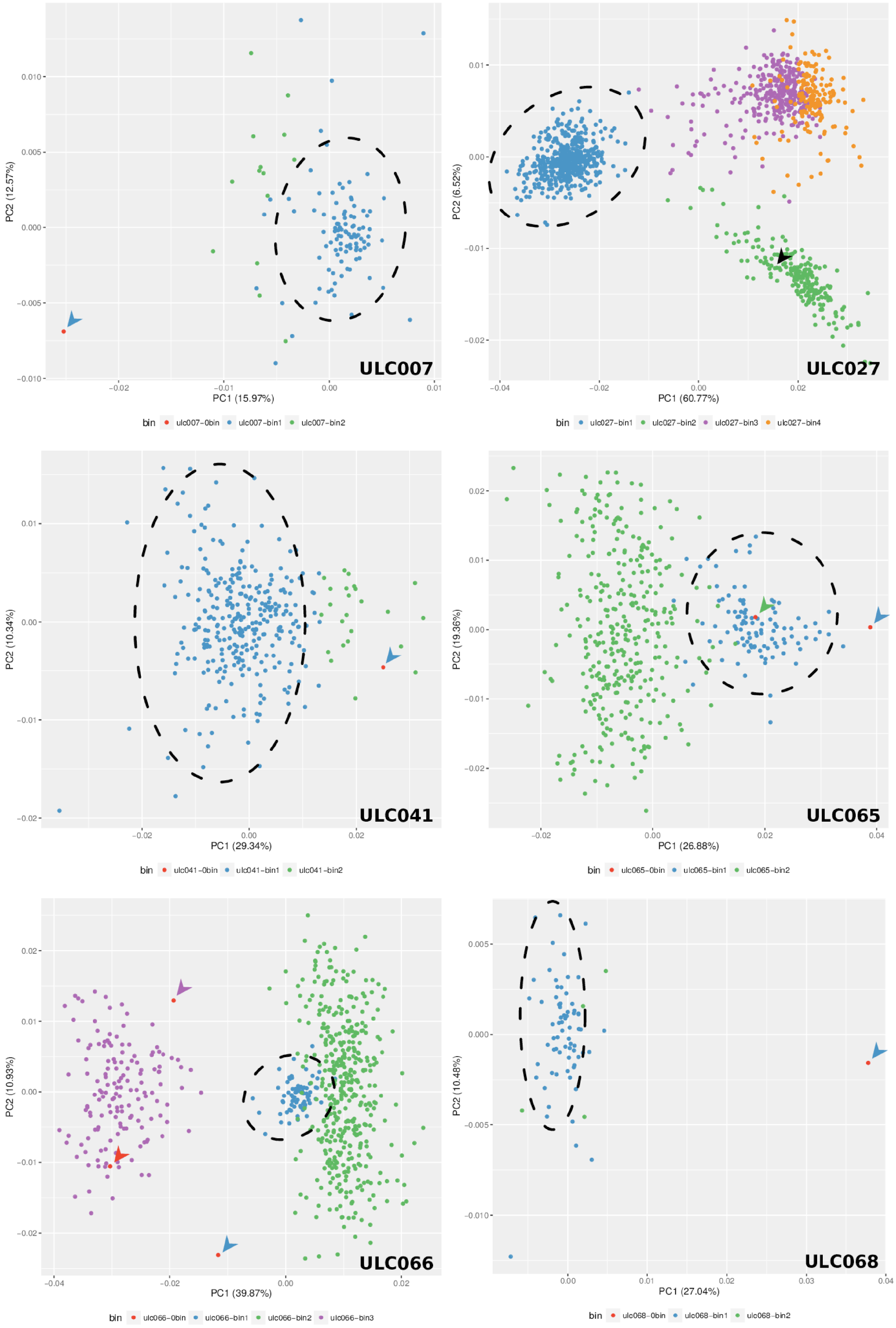
ULC007
PC1 (15.97%)
PC2 (12.57%)
bin ulc007-0bin ulc007-bin1 ulc007-bin2
ULC027
PC1 (60.77%)
PC2 (6.52%)
bin ulc027-bin1 ulc027-bin2 ulc027-bin3 ulc027-bin4
ULC041
PC1 (29.34%)
PC2 (10.34%)
bin ulc041-0bin ulc041-bin1 ulc041-bin2
ULC065
PC1 (26.88%)
PC2 (19.36%)
bin ulc065-0bin ulc065-bin1 ulc065-bin2
ULC066
PC1 (39.87%)
PC2 (10.93%)
bin ulc066-0bin ulc066-bin1 ulc066-bin2 ulc066-bin3
ULC068
PC1 (27.04%)
PC2 (10.48%)
bin ulc068-0bin ulc068-bin1 ulc068-bin2

**Figure S2**

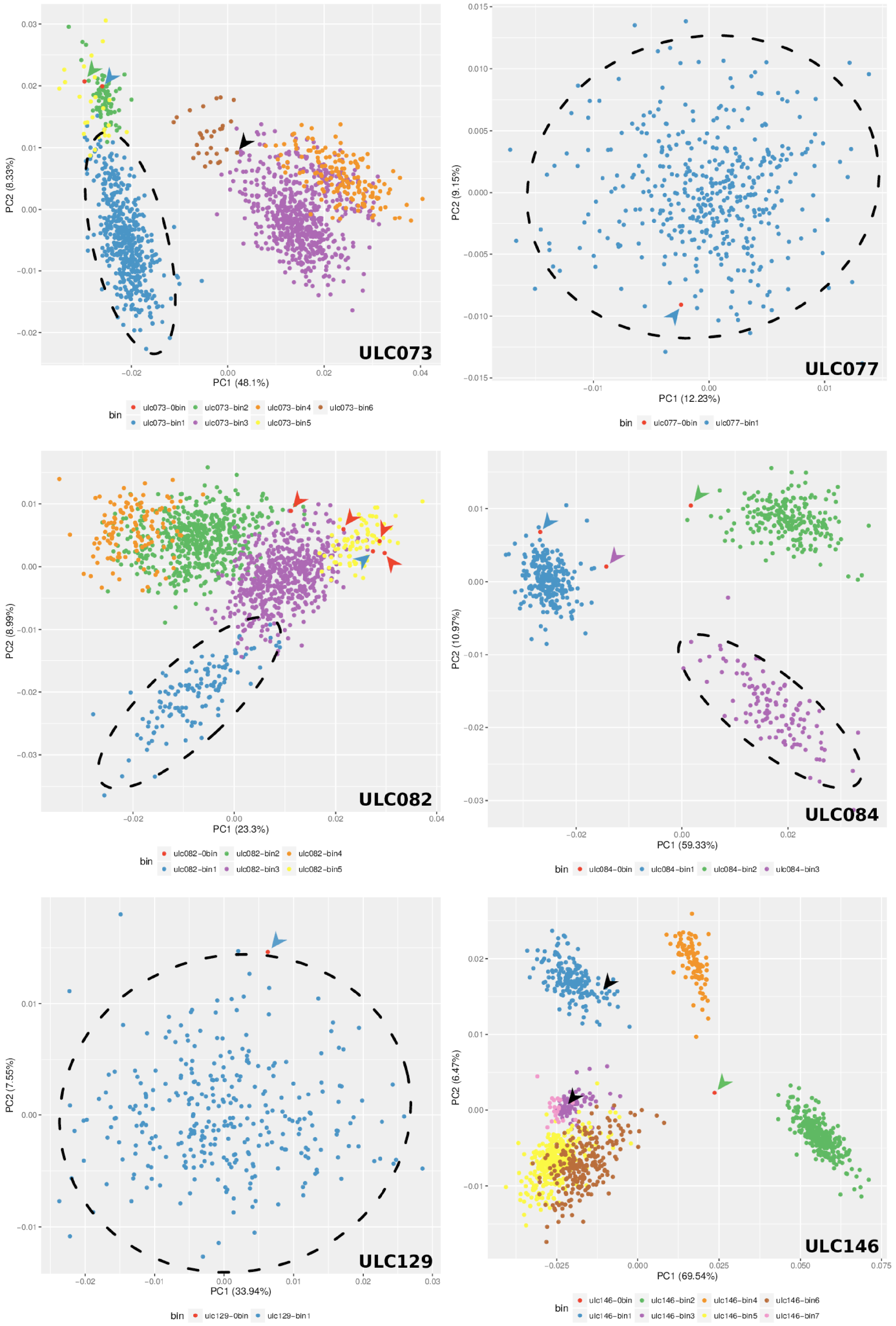


**Figure S2 (continued)**

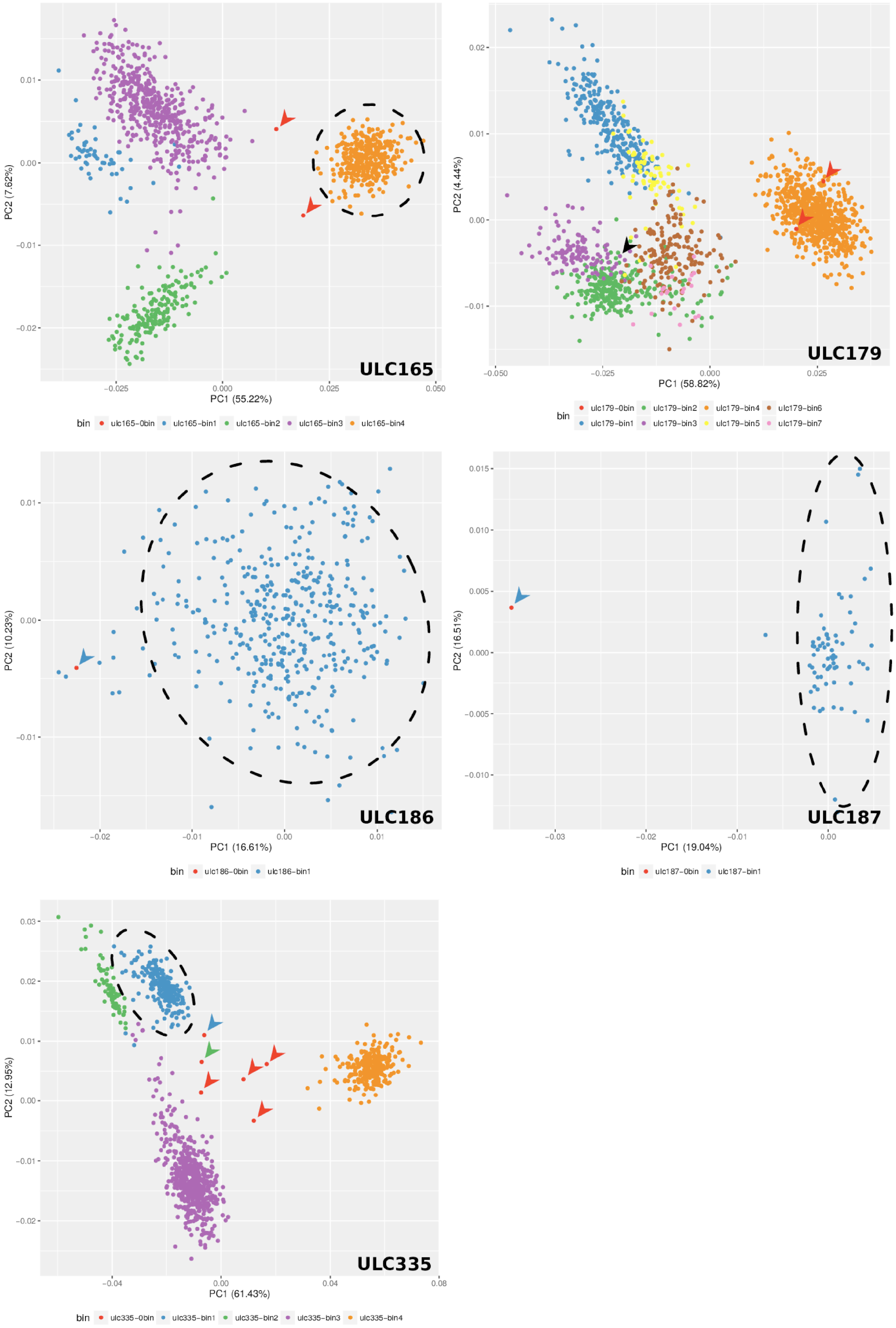


**Figure S2 (continued)**

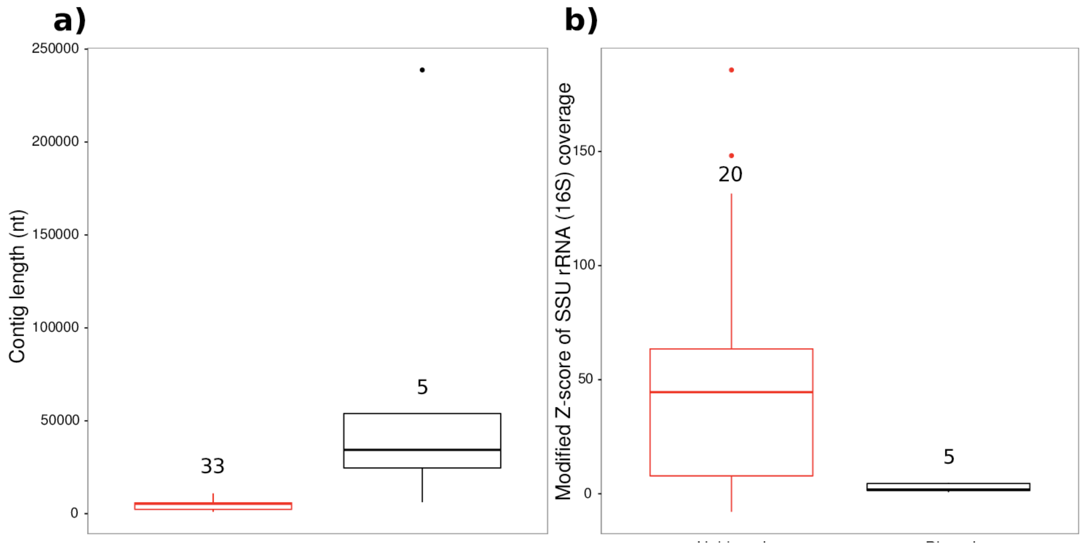
a)
250000
200000
150000
100000
50000
0
Contig length (nt)
33
5
b)
Modified Z-score of SSU rRNA (16S) coverage
150
100
50
0
20
5

**Figure S3**

### Supplemental Note 2

Our phylogenomic analysis of the cyanobacterial microbiome has shown that five sets of foreign bins (11 bins in total) appear indistinguishable in the tree (**Figure 2**). This can be explained by two causes: 1) association of the same or very closely related microbiome taxa to the Cyanobacteria, resulting in redundancy across multiple cyanobacterial cultures or 2) cross-contamination of cultures during strain propagation or of genomic DNA during nucleic acid extraction. In this supplemental note, we investigated the issue through all-vs-all BLASTN analyses on whole genome bins, using first the standard Average Nucleotide Identity (ANI) and then a newly developed metric, dubbed the Identical Genome Fraction (IGF), designed to better take into account the differential coverage of the bins during their comparison.

#### Methods

With the aim of not missing any possible case of redundancy or cross-contamination, we included an additional bin (called nobin) to each metagenome, containing the scaffolds rejected by MetaBAT during the binning process. ANI was computed with the Python script average_nucleotide_identity.py from the pyani software package (1) on genome bins split into overlapping fragments of 1020 nt, whereas IGF (introduced below) was computed with a custom Perl script (available upon request) on genome bins split into non-overlapping fragments (= pseudo-reads) of 250 nt. In both cases, BLASTN (2) was used to blast each split genome bin against all unsplit genome bins, using options inspired by (3) : -dust no -xdrop_ungap 150 -xdrop_gap 150 -xdrop_gap_final 150 -evalue 1e-15 -max_target_seqs 1.

To compute a neighbor-joining tree with the neighbor program of the PHYLIP package (4, 5), we first built a reduced version of the IGF matrix by selecting only the genome bins involved in any pair with an IGF value ≥75%. This IGF tree was then made comparable with the small phylogenomic tree of the co-cultivated organisms by pruning the latter tree with the prune-tree.pl script (part of the Bio-MUST-Core software package; D. Baurain; https://metacpan.org/release/Bio-MUST-Core). Details about sampling sites were retrieved from the BCCM/ULC web site (http://bccm.belspo.be/catalogues).

#### Results and Discussion

We identified 20 bins (including the 11 bins discussed above) sharing a very high ANI (≥98%) with another bin (**Table S3**), thus well over the species threshold (94%) defined in Goris et al. (2007) (3). However, ANI was not suitable for this study because it is computed only from homologous fragments (BLASTN identity ≥30%) (3), which can collectively cover only a (very) small part of the genomes (**Table S4**).

Hence, among the 20 high ANI matches, only 7 were computed on ≥80% of the genome bins (**Table S4**) (median = 52%, IQR = 68%). Moreover, ANI does not take into account any potential difference in completeness between the metagenomic bins under comparison. This is an important concern because it complicates the interpretation of ANI values. In contrast, the similarity distance of Mauve (6) would be a more suitable metric, because it is defined as the number of aligned nucleotides (estimated with nucmer; (7)) between the two genomes divided by their average size. Yet, when comparing two genome bins with a large difference in completeness, such use of the average size results in a mechanically reduced value of the similarity distance. As no tool was adapted to our purpose, i.e., to estimate bin similarity based on all fragments of two genome bins with possibly different completeness values, we developed the Identical Genome Fraction (IGF), a metric inspired from both the ANI and Mauve similarity distance. IGF is defined as the ratio between the number of identical fragments between two bins (preferring the BLASTN direction yielding the highest estimate) and the number of fragments in the smallest of the two bins: $IGF = max(A \cap B, B \cap A) / min(A, B)$. By default, two fragments are considered as identical when perfectly aligning (100% identity) over their full length (100% coverage).

We identified 13 bins (included in the 20 ANI matches) displaying a conspicuously high IGF (≥75%) with another bin (**Table S5**). The maximum IGF observed between any two bins (ULC146-bin4/ULC073-bin2) was 92.4%. We tested different identity thresholds and even with a threshold set at 25% (instead of 100%), no perfect match could be detected between our genome bins (at the exception of self-matches). This suggests that foreign bin redundancy is not the result of cross-contamination between cultures. As expected, the 11 bins identified in the microbiome phylogenomic tree are all comprised in these 13 high-IGF bins. ULC073-bin3 (included in the tree) and ULC066-bin3 (not in the tree, see main text) are the two additional bins with IGF ≥75%. **Figure S4** shows that, contrary to **Figure 2**, the 13 genome bins yield distinct terminal branches in the neighbor-joining tree based on the IGF matrix. This increased resolution is not surprising, considering that IGF is computed from full genome bins and not only from ribosomal proteins. Nevertheless, the bin clusters present in **Figure S4** all correspond to bin clusters observed in **Figure 2**, indicating that the two approaches are congruent for our purpose. Interestingly, all bin clusters comprise strains from different sampling sites, at the exception of the cluster composed of ULC146-bin1 and ULC165-bin3, which were both sampled in Antarctic Sør Rondane Mountains. Finally, no bin pair involving artificial bins of metagenomic contigs rejected by MetaBAT (nobins) was identified.

#### Conclusion

Our analysis failed to bring evidence for cross-contamination between BCCM/ULC cyanobacterial cultures. The 11 foreign genome bins that appear indistinguishable in the microbiome phylogenomic tree thus seem to correspond to very similar, but distinct, taxa present in multiple cultures despite distant sampling sites. This result suggests that these 11 genome bins belong to associated organisms in close interaction with Cyanobacteria, independently of their geographical location, even when Cyanobacteria were sampled from the opposite Earth poles. This is reminiscent of the Baas-Becking statement ‘Everything is everywhere, but the environment selects’ (8), where the ecological selection would mostly be defined by the presence of Cyanobacteria. However, the observation that the IGF metric can differentiate closely related bins indicates that evolutionary processes are at play, processes that could be due to long-term co-evolution.

Ideally, IGF should be repetitively estimated from multiple samples of the same culture to define a lower bound for identical organisms. Indeed, independent samples of the same organism would not necessarily yield an IGF of 100% due to slight variations during sequencing and assembly. Since we only had one sample of each culture in our experimental design, this could not be tested. For the same reason, we were not able to use MetaGen, a reference-free learning program not based on coverage (9). This is somewhat unfortunate as this very recent metagenomic tool reportedly performs better than MetaBAT when given multiple samples (≥ 10) of the same mixtures (9).

### Figures and Tables

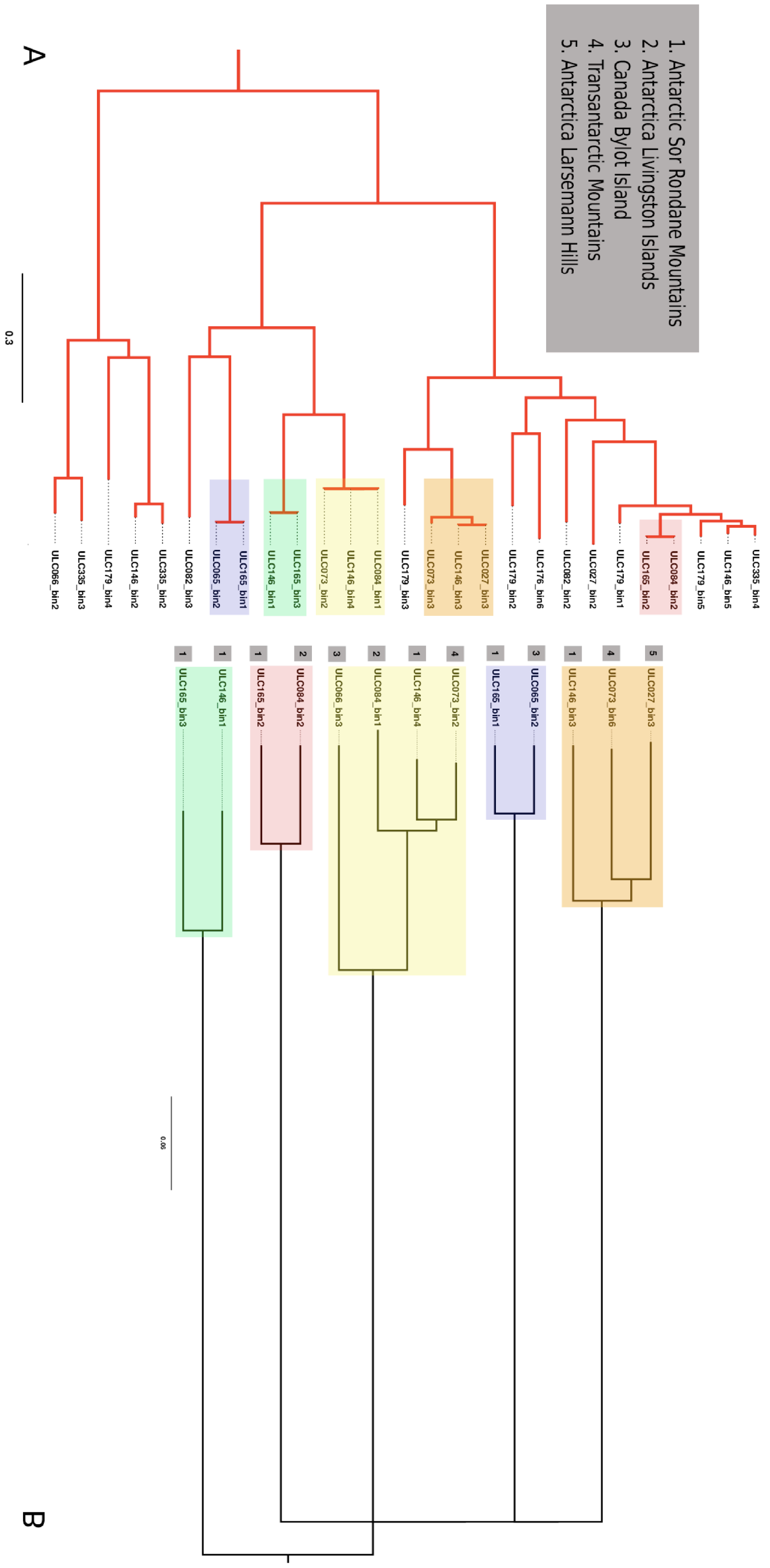


**Figure S4**

**Figure S4: Mirror comparison between a reduced version of the microbiome phylogenomic tree and an IGF neighbor-joining tree.** The phylogenomic tree was inferred under the CAT+$\Gamma_4$ model from a supermatrix made of 53 ribosomal genes. The 27 genome bins (**A**) were pruned out of this tree. The second tree (**B**) was inferred using neighbor-joining, based on a reduced version of the IGF matrix. Sampling information was collected from the BCCM/ULC website.

**Table S3: Average Nucleotide Identity identity matrix as computed by pyani.**

**Table S4: Average Nucleotide Identity coverage matrix as computed by pyani.**

**Table S5: Identical Genome Fraction matrix (IGF).**

| | ULC335-bin1 | ULC335-bin2 | ULC335-bin3 | ULC335-bin4 | ULC335-nobin | ULC007-bin1 | ULC007-bin2 | ULC007-nobin |
|---|---|---|---|---|---|---|---|---|
| **ULC335-bin1** | [redacted] | 0.942 | 0.704 | 0.723 | 0.956 | 0.743 | 0.741 | 0.739 |
| **ULC335-bin2** | 0.769 | [redacted] | 0.729 | 0.000 | 0.815 | 0.716 | 0.000 | 0.000 |
| **ULC335-bin3** | 0.704 | 0.730 | [redacted] | 0.714 | 0.787 | 0.723 | 0.000 | 0.000 |
| **ULC335-bin4** | 0.723 | 0.000 | 0.714 | [redacted] | 0.775 | 0.726 | 0.000 | 0.000 |
| **ULC335-nobin** | 0.912 | 0.890 | 0.818 | 0.774 | [redacted] | 0.736 | 0.696 | 0.812 |
| **ULC007-bin1** | 0.743 | 0.716 | 0.720 | 0.727 | 0.739 | [redacted] | 0.798 | 0.904 |
| **ULC007-bin2** | 0.753 | 0.000 | 0.000 | 0.000 | 0.000 | 0.786 | [redacted] | 0.843 |
| **ULC007-nobin** | 0.745 | 0.000 | 0.000 | 0.000 | 0.854 | 0.930 | 0.858 | [redacted] |
| **ULC027-bin1** | 0.738 | 0.725 | 0.714 | 0.745 | 0.735 | 0.748 | 0.782 | 0.737 |
| **ULC027-bin2** | 0.707 | 0.748 | 0.717 | 0.771 | 0.760 | 0.727 | 0.000 | 0.817 |
| **ULC027-bin3** | 0.712 | 0.717 | 0.723 | 0.767 | 0.786 | 0.734 | 0.000 | [redacted] |
| **ULC027-bin4** | 0.000 | 0.000 | 0.710 | 0.762 | 0.773 | 0.738 | 0.000 | 0.000 |
| **ULC027-nobin** | 0.735 | 0.718 | 0.720 | 0.779 | 0.784 | 0.738 | 0.704 | 0.851 |
| **ULC041-bin1** | 0.737 | 0.701 | 0.732 | 0.739 | 0.747 | 0.749 | 0.806 | 0.865 |
| **ULC041-bin2** | 0.725 | 0.000 | 0.780 | 0.727 | 0.742 | 0.747 | 0.760 | [redacted] |
| **ULC041-nobin** | 0.743 | 0.000 | 0.000 | 0.000 | 0.839 | 0.762 | 0.000 | 0.837 |
| **ULC065-bin1** | 0.726 | 0.000 | 0.729 | 0.750 | 0.741 | 0.737 | 0.810 | 0.951 |
| **ULC065-bin2** | 0.713 | 0.701 | 0.726 | 0.767 | 0.764 | 0.721 | 0.694 | 0.000 |
| **ULC065-nobin** | 0.734 | 0.741 | 0.724 | 0.790 | 0.805 | 0.753 | 0.809 | 0.845 |
| **ULC066-bin1** | 0.742 | 0.730 | 0.711 | 0.720 | 0.730 | 0.742 | 0.777 | 0.934 |
| **ULC066-bin2** | 0.727 | 0.727 | 0.742 | 0.706 | 0.751 | 0.735 | 0.000 | [redacted] |
| **ULC066-bin3** | 0.716 | 0.000 | 0.721 | 0.754 | 0.745 | 0.722 | 0.000 | 0.000 |
| **ULC066-nobin** | 0.725 | 0.724 | 0.747 | 0.786 | 0.788 | 0.730 | 0.687 | 0.862 |
| **ULC068-bin1** | 0.741 | 0.715 | 0.722 | 0.707 | 0.734 | 0.742 | 0.760 | 0.831 |
| **ULC068-bin2** | 0.769 | 0.000 | 0.717 | 0.000 | 0.779 | 0.739 | 0.000 | 0.758 |
| **ULC068-nobin** | 0.746 | 0.790 | 0.732 | 0.799 | 0.790 | 0.734 | 0.000 | 0.819 |
| **ULC073-bin1** | 0.736 | 0.725 | 0.727 | 0.746 | 0.745 | 0.751 | 0.695 | 0.871 |
| **ULC073-bin2** | 0.726 | 0.719 | 0.726 | 0.732 | 0.753 | 0.712 | 0.000 | 0.997 |
| **ULC073-bin3** | 0.722 | 0.708 | 0.719 | 0.769 | 0.782 | 0.724 | 0.000 | [redacted] |
| **ULC073-bin4** | 0.000 | 0.701 | 0.000 | 0.772 | 0.772 | 0.729 | 0.735 | 0.000 |
| **ULC073-bin5** | 0.746 | 0.000 | 0.000 | 0.000 | 0.730 | 0.755 | 0.761 | 0.740 |
| **ULC073-bin6** | 0.000 | 0.000 | 0.000 | 0.774 | 0.803 | 0.000 | 0.000 | 0.801 |
| **ULC073-nobin** | 0.736 | 0.716 | 0.723 | 0.773 | 0.796 | 0.749 | 0.719 | 0.856 |
| **ULC077-bin1** | 0.742 | 0.720 | 0.714 | 0.724 | 0.726 | 0.759 | 0.761 | 0.918 |
| **ULC077-nobin** | 0.737 | 0.725 | 0.727 | 0.792 | 0.794 | 0.773 | 0.797 | 0.911 |
| **ULC082-bin1** | 0.726 | 0.000 | 0.718 | 0.755 | 0.740 | 0.742 | 0.817 | 0.782 |
| **ULC082-bin2** | 0.711 | 0.739 | 0.719 | 0.860 | 0.824 | 0.729 | 0.000 | 0.000 |
| **ULC082-bin3** | 0.719 | 0.713 | 0.738 | 0.761 | 0.782 | 0.722 | 0.000 | 0.000 |
| **ULC082-bin4** | 0.000 | 0.718 | 0.000 | 0.904 | 0.884 | 0.717 | 0.000 | 0.000 |
| **ULC082-bin5** | 0.000 | 0.000 | 0.000 | 0.695 | 0.769 | 0.000 | 0.000 | 0.000 |
| **ULC082-nobin** | 0.719 | 0.711 | 0.736 | 0.858 | 0.837 | 0.725 | 0.698 | 0.822 |
| **ULC084-bin1** | 0.719 | 0.706 | 0.719 | 0.743 | 0.753 | 0.728 | 0.690 | 0.000 |
| **ULC084-bin2** | 0.724 | 0.702 | 0.711 | 0.810 | 0.798 | 0.725 | 0.000 | 0.709 |
| **ULC084-bin3** | 0.728 | 0.743 | 0.713 | 0.752 | 0.745 | 0.741 | 0.758 | 0.776 |
| **ULC084-nobin** | 0.720 | 0.728 | 0.721 | 0.779 | 0.799 | 0.741 | 0.786 | 0.842 |
| **ULC129-bin1** | 0.736 | 0.724 | 0.714 | 0.722 | 0.737 | 0.746 | 0.753 | 0.756 |
| **ULC129-nobin** | 0.742 | 0.741 | 0.744 | 0.772 | 0.801 | 0.750 | 0.727 | 0.821 |
| **ULC146-bin1** | 0.701 | 0.693 | 0.729 | 0.750 | 0.842 | 0.721 | 0.000 | 0.782 |
| **ULC146-bin2** | 0.721 | 0.773 | 0.726 | 0.738 | 0.729 | 0.725 | 0.000 | 0.000 |
| **ULC146-bin3** | 0.717 | 0.000 | 0.716 | 0.775 | 0.784 | 0.717 | 0.000 | 0.818 |
| **ULC146-bin4** | 0.716 | 0.717 | 0.730 | 0.742 | 0.751 | 0.722 | 0.000 | 0.000 |
| **ULC146-bin5** | 0.702 | 0.720 | 0.721 | 0.853 | 0.847 | 0.723 | 0.000 | 0.802 |
| **ULC146-bin6** | 0.000 | 0.000 | 0.000 | 0.774 | 0.770 | 0.737 | 0.000 | 0.000 |
| **ULC146-bin7** | 0.000 | 0.000 | 0.747 | 0.747 | 0.776 | 0.726 | 0.000 | 0.000 |
| **ULC146-nobin** | 0.748 | 0.762 | 0.725 | 0.805 | 0.802 | 0.749 | 0.774 | 0.831 |
| **ULC165-bin1** | 0.704 | 0.725 | 0.718 | 0.765 | 0.760 | 0.715 | 0.695 | 0.000 |
| **ULC165-bin2** | 0.722 | 0.703 | 0.706 | 0.811 | 0.799 | 0.725 | 0.000 | 0.692 |
| **ULC165-bin3** | 0.000 | 0.000 | 0.749 | 0.751 | 0.842 | 0.734 | 0.000 | 0.000 |
| **ULC165-bin4** | 0.749 | 0.709 | 0.733 | 0.690 | 0.747 | 0.758 | 0.786 | 0.800 |
| **ULC165-nobin** | 0.744 | 0.764 | 0.726 | 0.778 | 0.821 | 0.758 | 0.772 | 0.801 |
| **ULC179-bin1** | 0.712 | 0.717 | 0.721 | 0.827 | 0.818 | 0.726 | 0.000 | 0.000 |
| **ULC179-bin2** | 0.706 | 0.000 | 0.719 | 0.783 | 0.780 | 0.720 | 0.000 | 0.737 |
| **ULC179-bin3** | 0.717 | 0.700 | 0.713 | 0.767 | 0.763 | 0.731 | 0.000 | 0.815 |
| **ULC179-bin4** | 0.727 | 0.726 | 0.740 | 0.749 | 0.735 | 0.728 | 0.694 | 0.000 |
| **ULC179-bin5** | 0.693 | 0.000 | 0.717 | 0.832 | 0.820 | 0.727 | 0.000 | 0.000 |
| **ULC179-bin6** | 0.000 | 0.000 | 0.725 | 0.762 | 0.762 | 0.718 | 0.000 | 0.000 |
| **ULC179-bin7** | 0.000 | 0.000 | 0.000 | 0.758 | 0.756 | 0.000 | 0.000 | 0.000 |
| **ULC179-nobin** | 0.749 | 0.724 | 0.744 | 0.799 | 0.783 | 0.747 | 0.780 | 0.822 |
| **ULC186-bin1** | 0.738 | 0.723 | 0.727 | 0.744 | 0.746 | 0.751 | 0.794 | 0.776 |
| **ULC186-nobin** | 0.761 | 0.760 | 0.740 | 0.777 | 0.908 | 0.748 | 0.788 | 0.867 |
| **ULC187-bin1** | 0.747 | 0.729 | 0.717 | 0.718 | 0.744 | 0.741 | 0.701 | 0.778 |
| **ULC187-nobin** | 0.740 | 0.720 | 0.715 | 0.805 | 0.797 | 0.740 | 0.716 | 0.857 |

| | ULC027-bin1 | ULC027-bin2 | ULC027-bin3 | ULC027-bin4 | ULC027-nobin | ULC041-bin1 | ULC041-bin2 | ULC041-nobin |
|---|---|---|---|---|---|---|---|---|
| **ULC335-bin1** | 0.737 | 0.707 | 0.712 | 0.000 | 0.734 | 0.737 | 0.725 | 0.746 |
| **ULC335-bin2** | 0.725 | 0.748 | 0.000 | 0.000 | 0.719 | 0.701 | 0.000 | 0.000 |
| **ULC335-bin3** | 0.717 | 0.717 | 0.723 | 0.710 | 0.722 | 0.731 | 0.780 | 0.000 |
| **ULC335-bin4** | 0.750 | 0.772 | 0.770 | 0.759 | 0.782 | 0.743 | 0.727 | 0.000 |
| **ULC335-nobin** | 0.737 | 0.763 | 0.784 | 0.777 | 0.783 | 0.750 | 0.736 | 0.809 |
| **ULC007-bin1** | 0.748 | 0.727 | 0.732 | 0.738 | 0.734 | 0.749 | 0.746 | 0.754 |
| **ULC007-bin2** | 0.778 | 0.000 | 0.000 | 0.000 | 0.709 | 0.789 | 0.760 | 0.000 |
| **ULC007-nobin** | 0.793 | 0.854 | 0.928 | 0.000 | 0.854 | 0.871 | 0.989 | 0.837 |
| **ULC027-bin1** | | 0.734 | 0.730 | 0.706 | 0.859 | 0.761 | 0.760 | 0.749 |
| **ULC027-bin2** | 0.734 | | 0.758 | 0.746 | 0.772 | 0.736 | 0.727 | 0.769 |
| **ULC027-bin3** | 0.733 | 0.758 | | 0.771 | 0.791 | 0.735 | 0.716 | 0.711 |
| **ULC027-bin4** | 0.709 | 0.750 | 0.764 | | 0.784 | 0.728 | 0.000 | 0.000 |
| **ULC027-nobin** | 0.892 | 0.770 | 0.792 | 0.782 | | 0.777 | 0.829 | 0.866 |
| **ULC041-bin1** | 0.761 | 0.736 | 0.737 | 0.728 | 0.775 | | 0.883 | 0.929 |
| **ULC041-bin2** | 0.760 | 0.727 | 0.716 | 0.000 | 0.840 | 0.858 | | 0.878 |
| **ULC041-nobin** | 0.747 | 0.769 | 0.711 | 0.000 | 0.882 | 0.959 | 0.948 | |
| **ULC065-bin1** | 0.737 | 0.739 | 0.742 | 0.755 | 0.739 | 0.759 | 0.739 | 0.806 |
| **ULC065-bin2** | 0.719 | 0.753 | 0.750 | 0.745 | 0.752 | 0.743 | 0.724 | 0.741 |
| **ULC065-nobin** | 0.738 | 0.766 | 0.959 | 0.951 | 0.924 | 0.745 | 0.763 | 0.836 |
| **ULC066-bin1** | 0.738 | 0.726 | 0.723 | 0.000 | 0.739 | 0.741 | 0.741 | 0.876 |
| **ULC066-bin2** | 0.708 | 0.725 | 0.712 | 0.000 | 0.719 | 0.712 | 0.000 | 0.000 |
| **ULC066-bin3** | 0.722 | 0.729 | 0.716 | 0.000 | 0.989 | 0.731 | 0.000 | 0.000 |
| **ULC066-nobin** | 0.735 | 0.977 | 0.949 | 0.941 | 0.920 | 0.742 | 0.733 | 0.851 |
| **ULC068-bin1** | 0.739 | 0.722 | 0.729 | 0.717 | 0.757 | 0.740 | 0.767 | 0.769 |
| **ULC068-bin2** | 0.709 | 0.000 | 0.000 | 0.000 | 0.736 | 0.736 | 0.000 | 0.000 |
| **ULC068-nobin** | 0.733 | 0.899 | 0.776 | 0.769 | 0.923 | 0.751 | 0.755 | 0.818 |
| **ULC073-bin1** | 0.758 | 0.729 | 0.732 | 0.727 | 0.764 | 0.986 | 0.967 | 0.961 |
| **ULC073-bin2** | 0.727 | 0.735 | 0.743 | 0.733 | 0.988 | 0.729 | 0.000 | |
| **ULC073-bin3** | 0.725 | 0.754 | 0.987 | 0.985 | 0.893 | 0.732 | 0.718 | 0.724 |
| **ULC073-bin4** | 0.742 | 0.753 | 0.797 | 0.867 | 0.956 | 0.742 | 0.000 | |
| **ULC073-bin5** | 0.775 | 0.724 | 0.711 | 0.000 | 0.884 | 0.978 | 0.977 | 0.959 |
| **ULC073-bin6** | 0.700 | 0.818 | 0.995 | 0.859 | 0.926 | 0.749 | 0.000 | 0.000 |
| **ULC073-nobin** | 0.748 | 0.763 | 0.887 | 0.918 | 0.957 | 0.954 | 0.950 | 0.969 |
| **ULC077-bin1** | 0.747 | 0.724 | 0.721 | 0.727 | 0.745 | 0.750 | 0.749 | 0.774 |
| **ULC077-nobin** | 0.746 | 0.774 | 0.905 | 0.901 | 0.946 | 0.757 | 0.760 | 0.839 |
| **ULC082-bin1** | 0.739 | 0.739 | 0.741 | 0.728 | 0.751 | 0.757 | 0.742 | 0.761 |
| **ULC082-bin2** | 0.729 | 0.758 | 0.761 | 0.755 | 0.767 | 0.747 | 0.764 | 0.000 |
| **ULC082-bin3** | 0.735 | 0.747 | 0.752 | 0.737 | 0.755 | 0.740 | 0.000 | 0.723 |
| **ULC082-bin4** | 0.726 | 0.769 | 0.772 | 0.757 | 0.775 | 0.759 | 0.709 | 0.000 |
| **ULC082-bin5** | 0.000 | 0.710 | 0.730 | 0.000 | 0.762 | 0.704 | 0.000 | 0.000 |
| **ULC082-nobin** | 0.726 | 0.762 | 0.763 | 0.750 | 0.797 | 0.742 | 0.753 | 0.811 |
| **ULC084-bin1** | 0.731 | 0.733 | 0.729 | 0.721 | 0.986 | 0.725 | 0.000 | |
| **ULC084-bin2** | 0.732 | 0.759 | 0.761 | 0.751 | 0.761 | 0.746 | 0.732 | 0.000 |
| **ULC084-bin3** | 0.742 | 0.740 | 0.740 | 0.724 | 0.749 | 0.757 | 0.729 | 0.780 |
| **ULC084-nobin** | 0.755 | 0.775 | 0.987 | 0.985 | 0.880 | 0.738 | 0.722 | 0.853 |
| **ULC129-bin1** | 0.788 | 0.728 | 0.729 | 0.733 | 0.795 | 0.754 | 0.750 | 0.743 |
| **ULC129-nobin** | 0.821 | 0.985 | 0.907 | 0.909 | 0.942 | 0.783 | 0.790 | 0.848 |
| **ULC146-bin1** | 0.739 | 0.746 | 0.751 | 0.740 | 0.880 | 0.735 | 0.722 | 0.798 |
| **ULC146-bin2** | 0.719 | 0.763 | 0.717 | 0.000 | 0.719 | 0.701 | 0.000 | 0.000 |
| **ULC146-bin3** | 0.728 | 0.759 | 0.996 | 0.992 | 0.878 | 0.734 | 0.723 | 0.982 |
| **ULC146-bin4** | 0.728 | 0.730 | 0.733 | 0.723 | 0.987 | 0.739 | 0.702 | 0.000 |
| **ULC146-bin5** | 0.727 | 0.768 | 0.769 | 0.733 | 0.774 | 0.731 | 0.722 | 0.775 |
| **ULC146-bin6** | 0.730 | 0.756 | 0.828 | 0.777 | 0.981 | 0.739 | 0.000 | 0.000 |
| **ULC146-bin7** | 0.000 | 0.749 | 0.993 | 0.995 | 0.892 | 0.000 | 0.685 | 0.719 |
| **ULC146-nobin** | 0.743 | 0.766 | 0.806 | 0.805 | 0.954 | 0.745 | 0.723 | 0.824 |
| **ULC165-bin1** | 0.724 | 0.749 | 0.754 | 0.747 | 0.750 | 0.747 | 0.756 | 0.737 |
| **ULC165-bin2** | 0.733 | 0.757 | 0.760 | 0.749 | 0.762 | 0.738 | 0.732 | 0.000 |
| **ULC165-bin3** | 0.730 | 0.754 | 0.754 | 0.745 | 0.877 | 0.743 | 0.760 | 0.835 |
| **ULC165-bin4** | 0.744 | 0.725 | 0.717 | 0.000 | 0.746 | 0.746 | 0.751 | 0.000 |
| **ULC165-nobin** | 0.751 | 0.765 | 0.856 | 0.837 | 0.884 | 0.749 | 0.772 | 0.821 |
| **ULC179-bin1** | 0.724 | 0.761 | 0.766 | 0.745 | 0.775 | 0.746 | 0.707 | 0.720 |
| **ULC179-bin2** | 0.731 | 0.764 | 0.766 | 0.758 | 0.816 | 0.740 | 0.730 | 0.722 |
| **ULC179-bin3** | 0.730 | 0.759 | 0.776 | 0.771 | 0.771 | 0.731 | 0.746 | 0.799 |
| **ULC179-bin4** | 0.721 | 0.730 | 0.743 | 0.734 | 0.732 | 0.736 | 0.727 | 0.000 |
| **ULC179-bin5** | 0.726 | 0.762 | 0.763 | 0.751 | 0.780 | 0.732 | 0.000 | 0.000 |
| **ULC179-bin6** | 0.740 | 0.755 | 0.770 | 0.746 | 0.768 | 0.727 | 0.000 | 0.710 |
| **ULC179-bin7** | 0.000 | 0.761 | 0.765 | 0.000 | 0.755 | 0.000 | 0.000 | 0.000 |
| **ULC179-nobin** | 0.745 | 0.769 | 0.768 | 0.747 | 0.785 | 0.743 | 0.738 | 0.824 |
| **ULC186-bin1** | 0.759 | 0.741 | 0.726 | 0.726 | 0.747 | 0.842 | 0.832 | 0.854 |
| **ULC186-nobin** | 0.750 | 0.768 | 0.760 | 0.744 | 0.786 | 0.846 | 0.858 | 0.888 |
| **ULC187-bin1** | 0.744 | 0.727 | 0.723 | 0.000 | 0.735 | 0.737 | 0.739 | 0.767 |
| **ULC187-nobin** | 0.762 | 0.778 | 0.975 | 0.973 | 0.928 | 0.751 | 0.728 | 0.844 |

| | ULC065-bin1 | ULC065-bin2 | ULC065-nobin | ULC066-bin1 | ULC066-bin2 | ULC066-bin3 | ULC066-nobin | ULC068-bin1 |
|---|---|---|---|---|---|---|---|---|
| **ULC335-bin1** | 0.726 | 0.713 | 0.730 | 0.743 | 0.727 | 0.716 | 0.726 | 0.741 |
| **ULC335-bin2** | 0.000 | 0.000 | 0.743 | 0.727 | 0.727 | 0.000 | 0.724 | 0.711 |
| **ULC335-bin3** | 0.730 | 0.733 | 0.722 | 0.711 | 0.743 | 0.721 | 0.748 | 0.723 |
| **ULC335-bin4** | 0.751 | 0.770 | 0.791 | 0.723 | 0.706 | 0.754 | 0.789 | 0.707 |
| **ULC335-nobin** | 0.745 | 0.766 | 0.805 | 0.726 | 0.753 | 0.747 | 0.786 | 0.736 |
| **ULC007-bin1** | 0.737 | 0.719 | 0.753 | 0.742 | 0.727 | 0.719 | 0.732 | 0.742 |
| **ULC007-bin2** | 0.832 | 0.694 | 0.704 | 0.777 | 0.000 | 0.000 | 0.000 | 0.762 |
| **ULC007-nobin** | 0.963 | 0.000 | 0.893 | 0.942 | [redacted] | 0.000 | 0.918 | 0.859 |
| **ULC027-bin1** | 0.737 | 0.718 | 0.731 | 0.738 | 0.709 | 0.722 | 0.730 | 0.739 |
| **ULC027-bin2** | 0.740 | 0.753 | 0.762 | 0.728 | 0.725 | 0.715 | 0.989 | 0.727 |
| **ULC027-bin3** | 0.742 | 0.750 | 0.969 | 0.726 | 0.712 | 0.716 | 0.960 | 0.731 |
| **ULC027-bin4** | 0.755 | 0.752 | 0.968 | 0.000 | 0.000 | 0.000 | 0.956 | 0.717 |
| **ULC027-nobin** | 0.739 | 0.756 | 0.921 | 0.740 | 0.722 | 0.982 | 0.923 | 0.754 |
| **ULC041-bin1** | 0.758 | 0.744 | 0.740 | 0.741 | 0.712 | 0.729 | 0.744 | 0.740 |
| **ULC041-bin2** | 0.743 | 0.724 | 0.776 | 0.741 | 0.000 | 0.000 | 0.733 | 0.773 |
| **ULC041-nobin** | 0.809 | 0.741 | 0.909 | 0.871 | 0.000 | 0.000 | 0.902 | 0.772 |
| **ULC065-bin1** | [redacted] | 0.743 | 0.762 | 0.730 | 0.713 | 0.710 | 0.751 | 0.731 |
| **ULC065-bin2** | 0.742 | [redacted] | 0.762 | 0.699 | 0.740 | 0.748 | 0.756 | 0.726 |
| **ULC065-nobin** | 0.801 | 0.767 | [redacted] | 0.806 | 0.730 | 0.745 | 0.935 | 0.760 |
| **ULC066-bin1** | 0.729 | 0.699 | 0.816 | [redacted] | 0.710 | 0.728 | 0.788 | 0.824 |
| **ULC066-bin2** | 0.713 | 0.740 | 0.736 | 0.710 | [redacted] | 0.000 | 0.824 | 0.712 |
| **ULC066-bin3** | 0.715 | 0.748 | 0.744 | 0.728 | 0.000 | [redacted] | 0.793 | 0.727 |
| **ULC066-nobin** | 0.751 | 0.758 | 0.936 | 0.825 | 0.853 | 0.789 | [redacted] | 0.800 |
| **ULC068-bin1** | 0.732 | 0.724 | 0.757 | 0.825 | 0.712 | 0.721 | 0.800 | [redacted] |
| **ULC068-bin2** | 0.000 | 0.000 | 0.000 | 0.828 | 0.000 | 0.000 | 0.000 | 0.841 |
| **ULC068-nobin** | 0.754 | 0.977 | 0.934 | 0.835 | 0.734 | 0.748 | 0.913 | 0.915 |
| **ULC073-bin1** | 0.761 | 0.738 | 0.747 | 0.737 | 0.726 | 0.728 | 0.740 | 0.737 |
| **ULC073-bin2** | 0.734 | 0.742 | 0.749 | 0.733 | 0.726 | 0.993 | 0.995 | 0.721 |
| **ULC073-bin3** | 0.740 | 0.754 | 0.961 | 0.720 | 0.724 | 0.734 | 0.945 | 0.726 |
| **ULC073-bin4** | 0.747 | 0.759 | 0.785 | 0.000 | 0.740 | 0.776 | 0.783 | 0.720 |
| **ULC073-bin5** | 0.769 | 0.000 | 0.739 | 0.740 | 0.000 | [redacted] | 0.927 | 0.773 |
| **ULC073-bin6** | 0.726 | 0.000 | 0.987 | 0.686 | 0.703 | 0.000 | 0.978 | 0.000 |
| **ULC073-nobin** | 0.751 | 0.756 | 0.921 | 0.767 | 0.745 | 0.868 | 0.894 | 0.736 |
| **ULC077-bin1** | 0.736 | 0.727 | 0.744 | 0.747 | 0.716 | 0.708 | 0.748 | 0.744 |
| **ULC077-nobin** | 0.750 | 0.987 | 0.927 | 0.795 | 0.713 | 0.983 | 0.973 | 0.751 |
| **ULC082-bin1** | 0.821 | 0.741 | 0.755 | 0.736 | 0.722 | 0.725 | 0.742 | 0.737 |
| **ULC082-bin2** | 0.737 | 0.750 | 0.820 | 0.734 | 0.722 | 0.771 | 0.795 | 0.721 |
| **ULC082-bin3** | 0.740 | 0.763 | 0.766 | 0.721 | 0.724 | 0.723 | 0.756 | 0.723 |
| **ULC082-bin4** | 0.758 | 0.759 | 0.786 | 0.686 | 0.000 | 0.702 | 0.786 | 0.696 |
| **ULC082-bin5** | 0.000 | 0.873 | 0.742 | 0.000 | 0.000 | 0.000 | 0.732 | 0.000 |
| **ULC082-nobin** | 0.766 | 0.761 | 0.808 | 0.726 | 0.730 | 0.752 | 0.783 | 0.724 |
| **ULC084-bin1** | 0.731 | 0.739 | 0.743 | 0.731 | 0.716 | 0.992 | 0.993 | 0.740 |
| **ULC084-bin2** | 0.749 | 0.755 | 0.912 | 0.726 | 0.722 | 0.706 | 0.769 | 0.719 |
| **ULC084-bin3** | 0.823 | 0.744 | 0.782 | 0.729 | 0.719 | 0.729 | 0.750 | 0.735 |
| **ULC084-nobin** | 0.783 | 0.758 | 0.918 | 0.726 | 0.725 | 0.957 | 0.944 | 0.726 |
| **ULC129-bin1** | 0.743 | 0.732 | 0.731 | 0.737 | 0.725 | 0.709 | 0.730 | 0.738 |
| **ULC129-nobin** | 0.747 | 0.759 | 0.909 | 0.737 | 0.743 | 0.981 | 0.979 | 0.737 |
| **ULC146-bin1** | 0.743 | 0.760 | 0.952 | 0.723 | 0.709 | 0.748 | 0.810 | 0.725 |
| **ULC146-bin2** | 0.710 | 0.707 | 0.000 | 0.735 | 0.731 | 0.000 | 0.731 | 0.710 |
| **ULC146-bin3** | 0.739 | 0.751 | 0.966 | 0.717 | 0.727 | 0.757 | 0.958 | 0.727 |
| **ULC146-bin4** | 0.727 | 0.736 | 0.744 | 0.715 | 0.721 | 0.993 | 0.995 | 0.724 |
| **ULC146-bin5** | 0.742 | 0.754 | 0.782 | 0.710 | 0.713 | 0.715 | 0.776 | 0.715 |
| **ULC146-bin6** | 0.744 | 0.748 | 0.780 | 0.717 | 0.710 | 0.713 | 0.775 | 0.724 |
| **ULC146-bin7** | 0.737 | 0.740 | 0.972 | 0.000 | 0.000 | 0.000 | 0.960 | 0.708 |
| **ULC146-nobin** | 0.746 | 0.759 | 0.888 | 0.743 | 0.736 | 0.891 | 0.873 | 0.744 |
| **ULC165-bin1** | 0.743 | 0.998 | 0.941 | 0.707 | 0.742 | 0.722 | 0.753 | 0.717 |
| **ULC165-bin2** | 0.746 | 0.755 | 0.909 | 0.730 | 0.708 | 0.729 | 0.772 | 0.710 |
| **ULC165-bin3** | 0.744 | 0.762 | 0.954 | 0.743 | 0.750 | 0.742 | 0.797 | 0.734 |
| **ULC165-bin4** | 0.735 | 0.733 | 0.768 | 0.749 | 0.727 | 0.698 | 0.731 | 0.741 |
| **ULC165-nobin** | 0.746 | 0.804 | 0.900 | 0.746 | 0.722 | 0.753 | 0.854 | 0.748 |
| **ULC179-bin1** | 0.741 | 0.755 | 0.780 | 0.710 | 0.704 | 0.732 | 0.774 | 0.708 |
| **ULC179-bin2** | 0.741 | 0.755 | 0.768 | 0.720 | 0.724 | 0.746 | 0.783 | 0.728 |
| **ULC179-bin3** | 0.734 | 0.755 | 0.768 | 0.725 | 0.729 | 0.724 | 0.769 | 0.722 |
| **ULC179-bin4** | 0.718 | 0.734 | 0.732 | 0.720 | 0.724 | 0.736 | 0.734 | 0.726 |
| **ULC179-bin5** | 0.741 | 0.754 | 0.777 | 0.697 | 0.000 | 0.747 | 0.776 | 0.717 |
| **ULC179-bin6** | 0.733 | 0.766 | 0.772 | 0.714 | 0.000 | 0.723 | 0.766 | 0.735 |
| **ULC179-bin7** | 0.000 | 0.000 | 0.750 | 0.000 | 0.000 | 0.000 | 0.754 | 0.000 |
| **ULC179-nobin** | 0.750 | 0.756 | 0.780 | 0.752 | 0.737 | 0.716 | 0.784 | 0.747 |
| **ULC186-bin1** | 0.756 | 0.736 | 0.743 | 0.739 | 0.713 | 0.732 | 0.737 | 0.739 |
| **ULC186-nobin** | 0.752 | 0.764 | 0.808 | 0.743 | 0.739 | 0.758 | 0.797 | 0.732 |
| **ULC187-bin1** | 0.732 | 0.709 | 0.729 | 0.799 | 0.734 | 0.731 | 0.739 | 0.797 |
| **ULC187-nobin** | 0.745 | 0.764 | 0.946 | 0.818 | 0.739 | 0.828 | 0.912 | 0.807 |

| | ULC068-bin2 | ULC068-nobin | ULC073-bin1 | ULC073-bin2 | ULC073-bin3 | ULC073-bin4 | ULC073-bin5 | ULC073-bin6 |
|---|---|---|---|---|---|---|---|---|
| **ULC335-bin1** | 0.769 | 0.739 | 0.736 | 0.725 | 0.722 | 0.689 | 0.746 | 0.000 |
| **ULC335-bin2** | 0.000 | 0.782 | 0.750 | 0.716 | 0.708 | 0.701 | 0.000 | 0.000 |
| **ULC335-bin3** | 0.717 | 0.727 | 0.728 | 0.726 | 0.719 | 0.000 | 0.000 | 0.000 |
| **ULC335-bin4** | 0.000 | 0.795 | 0.753 | 0.733 | 0.773 | 0.770 | 0.000 | 0.774 |
| **ULC335-nobin** | 0.778 | 0.784 | 0.748 | 0.757 | 0.779 | 0.771 | 0.730 | 0.799 |
| **ULC007-bin1** | 0.739 | 0.739 | 0.752 | 0.714 | 0.724 | 0.729 | 0.751 | 0.000 |
| **ULC007-bin2** | 0.000 | 0.000 | 0.706 | 0.000 | 0.000 | 0.735 | 0.761 | 0.000 |
| **ULC007-nobin** | 0.756 | 0.833 | 0.886 | 0.998 | 0.928 | 0.000 | 0.741 | 0.801 |
| **ULC027-bin1** | 0.709 | 0.734 | 0.758 | 0.724 | 0.720 | 0.742 | 0.770 | 0.700 |
| **ULC027-bin2** | 0.000 | 0.895 | 0.729 | 0.736 | 0.756 | 0.753 | 0.724 | 0.818 |
| **ULC027-bin3** | 0.000 | 0.776 | 0.733 | 0.742 | 0.992 | 0.795 | 0.711 | 0.985 |
| **ULC027-bin4** | 0.000 | 0.774 | 0.730 | 0.730 | 0.988 | 0.870 | 0.000 | 0.858 |
| **ULC027-nobin** | 0.738 | 0.909 | 0.763 | 0.985 | 0.882 | 0.938 | 0.891 | 0.896 |
| **ULC041-bin1** | 0.736 | 0.746 | 0.985 | 0.726 | 0.734 | 0.738 | 0.968 | 0.749 |
| **ULC041-bin2** | 0.000 | 0.743 | 0.975 | 0.000 | 0.718 | 0.000 | 0.976 | 0.000 |
| **ULC041-nobin** | 0.000 | 0.844 | 0.955 | | 0.724 | | 0.951 | 0.000 |
| **ULC065-bin1** | 0.000 | 0.752 | 0.761 | 0.734 | 0.741 | 0.746 | 0.769 | 0.726 |
| **ULC065-bin2** | 0.000 | 0.977 | 0.741 | 0.742 | 0.752 | 0.754 | 0.000 | 0.000 |
| **ULC065-nobin** | 0.000 | 0.921 | 0.745 | 0.751 | 0.955 | 0.778 | 0.751 | 0.981 |
| **ULC066-bin1** | 0.827 | 0.817 | 0.737 | 0.727 | 0.718 | 0.000 | 0.740 | 0.686 |
| **ULC066-bin2** | 0.000 | 0.736 | 0.726 | 0.726 | 0.724 | 0.740 | 0.000 | 0.703 |
| **ULC066-bin3** | 0.000 | 0.743 | 0.728 | 0.994 | 0.733 | 0.776 | | 0.000 |
| **ULC066-nobin** | 0.000 | 0.897 | 0.740 | 0.992 | 0.936 | 0.778 | 0.930 | 0.971 |
| **ULC068-bin1** | 0.844 | 0.836 | 0.737 | 0.719 | 0.726 | 0.720 | 0.769 | 0.000 |
| **ULC068-bin2** | | 0.750 | 0.000 | 0.708 | 0.000 | 0.000 | 0.000 | 0.000 |
| **ULC068-nobin** | 0.835 | | 0.742 | 0.753 | 0.772 | 0.763 | 0.724 | 0.832 |
| **ULC073-bin1** | 0.000 | 0.741 | | 0.728 | 0.734 | 0.748 | 0.895 | 0.769 |
| **ULC073-bin2** | 0.708 | 0.752 | 0.728 | | 0.726 | 0.749 | 0.717 | 0.000 |
| **ULC073-bin3** | 0.000 | 0.768 | 0.733 | 0.727 | | 0.800 | 0.709 | 0.807 |
| **ULC073-bin4** | 0.000 | 0.765 | 0.746 | 0.749 | 0.801 | | 0.000 | 0.000 |
| **ULC073-bin5** | 0.000 | 0.736 | 0.869 | 0.726 | 0.709 | 0.000 | | 0.000 |
| **ULC073-bin6** | 0.000 | 0.834 | 0.769 | 0.000 | 0.816 | 0.000 | 0.000 | |
| **ULC073-nobin** | 0.738 | 0.908 | 0.861 | 0.780 | 0.796 | 0.828 | 0.952 | 0.818 |
| **ULC077-bin1** | 0.745 | 0.768 | 0.749 | 0.731 | 0.728 | 0.728 | 0.772 | 0.000 |
| **ULC077-nobin** | 0.727 | 0.961 | 0.752 | 0.989 | 0.900 | 0.803 | 0.973 | 0.960 |
| **ULC082-bin1** | 0.000 | 0.744 | 0.756 | 0.726 | 0.739 | 0.732 | 0.762 | 0.000 |
| **ULC082-bin2** | 0.746 | 0.777 | 0.744 | 0.742 | 0.759 | 0.752 | 0.000 | 0.782 |
| **ULC082-bin3** | 0.000 | 0.755 | 0.741 | 0.750 | 0.749 | 0.747 | 0.000 | 0.767 |
| **ULC082-bin4** | 0.000 | 0.785 | 0.730 | 0.736 | 0.766 | 0.745 | 0.000 | 0.781 |
| **ULC082-bin5** | 0.000 | 0.733 | 0.000 | 0.730 | 0.710 | 0.000 | 0.000 | 0.000 |
| **ULC082-nobin** | 0.774 | 0.784 | 0.743 | 0.753 | 0.760 | 0.764 | 0.770 | 0.801 |
| **ULC084-bin1** | 0.719 | 0.749 | 0.733 | 0.998 | 0.725 | 0.718 | 0.731 | 0.726 |
| **ULC084-bin2** | 0.000 | 0.885 | 0.736 | 0.727 | 0.756 | 0.752 | 0.000 | 0.720 |
| **ULC084-bin3** | 0.720 | 0.742 | 0.757 | 0.737 | 0.737 | 0.768 | 0.754 | 0.000 |
| **ULC084-nobin** | 0.788 | 0.798 | 0.740 | 0.973 | 0.984 | 0.790 | 0.977 | 0.985 |
| **ULC129-bin1** | 0.768 | 0.754 | 0.753 | 0.725 | 0.736 | 0.729 | 0.752 | 0.708 |
| **ULC129-nobin** | 0.744 | 0.911 | 0.797 | 0.991 | 0.900 | 0.868 | 0.921 | 0.962 |
| **ULC146-bin1** | 0.000 | 0.791 | 0.740 | 0.751 | 0.751 | 0.749 | 0.736 | 0.761 |
| **ULC146-bin2** | 0.000 | 0.775 | 0.683 | 0.729 | 0.714 | 0.000 | 0.000 | 0.692 |
| **ULC146-bin3** | 0.772 | 0.773 | 0.738 | 0.733 | 0.992 | 0.792 | 0.000 | 0.984 |
| **ULC146-bin4** | 0.708 | 0.758 | 0.727 | 0.999 | 0.726 | 0.749 | 0.911 | 0.000 |
| **ULC146-bin5** | 0.000 | 0.774 | 0.740 | 0.733 | 0.760 | 0.753 | 0.000 | 0.793 |
| **ULC146-bin6** | 0.724 | 0.758 | 0.739 | 0.759 | 0.820 | 0.841 | 0.000 | 0.772 |
| **ULC146-bin7** | 0.000 | 0.761 | 0.703 | 0.735 | 0.993 | 0.770 | 0.730 | 0.830 |
| **ULC146-nobin** | 0.744 | 0.896 | 0.744 | 0.870 | 0.843 | 0.919 | 0.944 | 0.928 |
| **ULC165-bin1** | 0.000 | 0.982 | 0.744 | 0.741 | 0.751 | 0.738 | 0.000 | 0.732 |
| **ULC165-bin2** | 0.000 | 0.889 | 0.734 | 0.734 | 0.760 | 0.753 | 0.691 | 0.760 |
| **ULC165-bin3** | 0.000 | 0.800 | 0.730 | 0.755 | 0.753 | 0.742 | 0.727 | 0.750 |
| **ULC165-bin4** | 0.708 | 0.758 | 0.755 | 0.734 | 0.000 | 0.000 | 0.774 | 0.000 |
| **ULC165-nobin** | 0.754 | 0.830 | 0.749 | 0.750 | 0.859 | 0.878 | 0.753 | 0.940 |
| **ULC179-bin1** | 0.000 | 0.774 | 0.744 | 0.737 | 0.761 | 0.758 | 0.734 | 0.754 |
| **ULC179-bin2** | 0.000 | 0.767 | 0.741 | 0.732 | 0.763 | 0.759 | 0.721 | 0.747 |
| **ULC179-bin3** | 0.755 | 0.767 | 0.738 | 0.731 | 0.777 | 0.775 | 0.000 | 0.794 |
| **ULC179-bin4** | 0.000 | 0.726 | 0.733 | 0.735 | 0.737 | 0.763 | 0.725 | 0.000 |
| **ULC179-bin5** | 0.000 | 0.773 | 0.733 | 0.749 | 0.760 | 0.763 | 0.000 | 0.791 |
| **ULC179-bin6** | 0.000 | 0.762 | 0.734 | 0.737 | 0.765 | 0.760 | 0.000 | 0.754 |
| **ULC179-bin7** | 0.000 | 0.783 | 0.000 | 0.725 | 0.743 | 0.789 | 0.000 | 0.777 |
| **ULC179-nobin** | 0.761 | 0.783 | 0.747 | 0.748 | 0.762 | 0.773 | 0.737 | 0.819 |
| **ULC186-bin1** | 0.718 | 0.742 | 0.843 | 0.728 | 0.729 | 0.746 | 0.848 | 0.749 |
| **ULC186-nobin** | 0.811 | 0.807 | 0.847 | 0.770 | 0.773 | 0.736 | 0.854 | 0.836 |
| **ULC187-bin1** | 0.877 | 0.784 | 0.737 | 0.727 | 0.728 | 0.705 | 0.741 | 0.000 |
| **ULC187-nobin** | 0.827 | 0.902 | 0.747 | 0.823 | 0.973 | 0.926 | 0.827 | 0.986 |

| | ULC073-nobin | ULC077-bin1 | ULC077-nobin | ULC082-bin1 | ULC082-bin2 | ULC082-bin3 | ULC082-bin4 | ULC082-bin5 |
|---|---|---|---|---|---|---|---|---|
| **ULC335-bin1** | 0.735 | 0.742 | 0.740 | 0.727 | 0.710 | 0.719 | 0.000 | 0.000 |
| **ULC335-bin2** | 0.684 | 0.720 | 0.725 | 0.000 | 0.739 | 0.713 | 0.718 | 0.000 |
| **ULC335-bin3** | 0.723 | 0.712 | 0.722 | 0.718 | 0.721 | 0.739 | 0.695 | 0.000 |
| **ULC335-bin4** | 0.771 | 0.724 | 0.791 | 0.755 | 0.866 | 0.761 | 0.905 | 0.695 |
| **ULC335-nobin** | 0.794 | 0.726 | 0.790 | 0.743 | 0.819 | 0.781 | 0.872 | 0.763 |
| **ULC007-bin1** | 0.752 | 0.759 | 0.762 | 0.741 | 0.723 | 0.725 | 0.717 | 0.000 |
| **ULC007-bin2** | 0.719 | 0.784 | 0.824 | 0.842 | 0.000 | 0.000 | 0.000 | 0.000 |
| **ULC007-nobin** | 0.887 | 0.856 | 0.874 | 0.807 | 0.933 | 0.000 | 0.000 | 0.000 |
| **ULC027-bin1** | 0.745 | 0.747 | 0.745 | 0.738 | 0.729 | 0.734 | 0.726 | 0.000 |
| **ULC027-bin2** | 0.757 | 0.723 | 0.767 | 0.737 | 0.755 | 0.750 | 0.766 | 0.710 |
| **ULC027-bin3** | 0.908 | 0.720 | 0.916 | 0.739 | 0.760 | 0.749 | 0.769 | 0.730 |
| **ULC027-bin4** | 0.942 | 0.727 | 0.897 | 0.731 | 0.756 | 0.738 | 0.747 | 0.000 |
| **ULC027-nobin** | 0.955 | 0.747 | 0.940 | 0.752 | 0.767 | 0.758 | 0.772 | 0.720 |
| **ULC041-bin1** | 0.951 | 0.750 | 0.749 | 0.756 | 0.740 | 0.737 | 0.757 | 0.704 |
| **ULC041-bin2** | 0.956 | 0.749 | 0.763 | 0.742 | 0.764 | 0.000 | 0.709 | 0.000 |
| **ULC041-nobin** | 0.973 | 0.778 | 0.862 | 0.770 | 0.000 | 0.723 | 0.000 | 0.000 |
| **ULC065-bin1** | 0.751 | 0.736 | 0.746 | 0.821 | 0.738 | 0.740 | 0.753 | 0.000 |
| **ULC065-bin2** | 0.753 | 0.727 | 0.991 | 0.744 | 0.747 | 0.760 | 0.753 | 0.698 |
| **ULC065-nobin** | 0.922 | 0.743 | 0.919 | 0.756 | 0.818 | 0.770 | 0.779 | 0.727 |
| **ULC066-bin1** | 0.768 | 0.747 | 0.799 | 0.733 | 0.725 | 0.722 | 0.689 | 0.000 |
| **ULC066-bin2** | 0.736 | 0.713 | 0.696 | 0.722 | 0.709 | 0.727 | 0.000 | 0.000 |
| **ULC066-bin3** | 0.867 | 0.708 | 0.990 | 0.736 | 0.767 | 0.725 | 0.702 | 0.000 |
| **ULC066-nobin** | 0.893 | 0.741 | 0.966 | 0.744 | 0.793 | 0.754 | 0.782 | 0.732 |
| **ULC068-bin1** | 0.730 | 0.744 | 0.754 | 0.732 | 0.721 | 0.725 | 0.696 | 0.000 |
| **ULC068-bin2** | 0.738 | 0.745 | 0.727 | 0.000 | 0.746 | 0.000 | 0.000 | 0.000 |
| **ULC068-nobin** | 0.914 | 0.738 | 0.965 | 0.747 | 0.779 | 0.759 | 0.787 | 0.733 |
| **ULC073-bin1** | 0.832 | 0.749 | 0.745 | 0.756 | 0.742 | 0.739 | 0.724 | 0.000 |
| **ULC073-bin2** | 0.762 | 0.733 | 0.993 | 0.726 | 0.740 | 0.749 | 0.736 | 0.734 |
| **ULC073-bin3** | 0.791 | 0.738 | 0.897 | 0.739 | 0.756 | 0.744 | 0.763 | 0.710 |
| **ULC073-bin4** | 0.827 | 0.725 | 0.805 | 0.742 | 0.756 | 0.747 | 0.745 | 0.000 |
| **ULC073-bin5** | 0.902 | 0.772 | 0.972 | 0.770 | 0.000 | 0.000 | 0.000 | 0.000 |
| **ULC073-bin6** | 0.810 | 0.000 | 0.966 | 0.000 | 0.760 | 0.767 | 0.781 | 0.000 |
| **ULC073-nobin** | | 0.740 | 0.883 | 0.747 | 0.762 | 0.763 | 0.770 | 0.813 |
| **ULC077-bin1** | 0.739 | | 0.928 | 0.738 | 0.733 | 0.734 | 0.720 | 0.000 |
| **ULC077-nobin** | 0.890 | 0.963 | | 0.750 | 0.778 | 0.762 | 0.812 | 0.710 |
| **ULC082-bin1** | 0.743 | 0.738 | 0.750 | | 0.750 | 0.734 | 0.733 | |
| **ULC082-bin2** | 0.760 | 0.734 | 0.775 | 0.746 | | 0.744 | 0.761 | 0.000 |
| **ULC082-bin3** | 0.764 | 0.731 | 0.765 | 0.733 | 0.741 | | 0.740 | 0.729 |
| **ULC082-bin4** | 0.764 | 0.729 | 0.817 | 0.735 | 0.760 | 0.740 | | 0.000 |
| **ULC082-bin5** | 0.802 | 0.000 | 0.710 | 0.000 | 0.000 | 0.734 | 0.000 | |
| **ULC082-nobin** | 0.792 | 0.739 | 0.788 | 0.757 | 0.783 | 0.781 | 0.803 | 0.796 |
| **ULC084-bin1** | 0.871 | 0.734 | 0.993 | 0.724 | 0.734 | 0.745 | 0.725 | 0.742 |
| **ULC084-bin2** | 0.756 | 0.727 | 0.958 | 0.743 | 0.800 | 0.740 | 0.835 | 0.696 |
| **ULC084-bin3** | 0.756 | 0.738 | 0.746 | 0.986 | 0.756 | 0.736 | 0.754 | 0.000 |
| **ULC084-nobin** | 0.915 | 0.745 | 0.946 | 0.923 | 0.832 | 0.766 | 0.772 | 0.730 |
| **ULC129-bin1** | 0.752 | 0.744 | 0.745 | 0.743 | 0.726 | 0.734 | 0.693 | 0.000 |
| **ULC129-nobin** | 0.924 | 0.737 | 0.969 | 0.752 | 0.778 | 0.769 | 0.760 | 0.719 |
| **ULC146-bin1** | 0.950 | 0.726 | 0.782 | 0.745 | 0.743 | 0.785 | 0.745 | 0.738 |
| **ULC146-bin2** | 0.000 | 0.719 | 0.717 | 0.000 | 0.747 | 0.723 | 0.716 | 0.000 |
| **ULC146-bin3** | 0.926 | 0.713 | 0.904 | 0.737 | 0.753 | 0.752 | 0.759 | 0.726 |
| **ULC146-bin4** | 0.899 | 0.718 | 0.993 | 0.721 | 0.739 | 0.741 | 0.739 | 0.731 |
| **ULC146-bin5** | 0.763 | 0.723 | 0.792 | 0.737 | 0.827 | 0.744 | 0.841 | 0.730 |
| **ULC146-bin6** | 0.964 | 0.721 | 0.793 | 0.751 | 0.749 | 0.754 | 0.755 | 0.750 |
| **ULC146-bin7** | 0.933 | 0.723 | 0.923 | 0.735 | 0.747 | 0.746 | 0.760 | 0.000 |
| **ULC146-nobin** | 0.945 | 0.751 | 0.877 | 0.749 | 0.771 | 0.751 | 0.792 | 0.785 |
| **ULC165-bin1** | 0.750 | 0.726 | 0.993 | 0.738 | 0.740 | 0.759 | 0.756 | 0.730 |
| **ULC165-bin2** | 0.757 | 0.729 | 0.958 | 0.745 | 0.810 | 0.743 | 0.821 | 0.000 |
| **ULC165-bin3** | 0.951 | 0.727 | 0.783 | 0.740 | 0.747 | 0.809 | 0.741 | 0.733 |
| **ULC165-bin4** | 0.745 | 0.757 | 0.736 | 0.737 | 0.000 | 0.720 | 0.685 | 0.000 |
| **ULC165-nobin** | 0.900 | 0.758 | 0.822 | 0.747 | 0.764 | 0.764 | 0.767 | 0.728 |
| **ULC179-bin1** | 0.759 | 0.724 | 0.788 | 0.742 | 0.799 | 0.748 | 0.822 | 0.720 |
| **ULC179-bin2** | 0.768 | 0.729 | 0.767 | 0.741 | 0.757 | 0.741 | 0.766 | 0.731 |
| **ULC179-bin3** | 0.773 | 0.720 | 0.763 | 0.736 | 0.756 | 0.737 | 0.761 | 0.000 |
| **ULC179-bin4** | 0.734 | 0.732 | 0.742 | 0.728 | 0.726 | 0.744 | 0.736 | 0.000 |
| **ULC179-bin5** | 0.756 | 0.718 | 0.785 | 0.736 | 0.811 | 0.742 | 0.829 | 0.000 |
| **ULC179-bin6** | 0.765 | 0.722 | 0.759 | 0.722 | 0.758 | 0.739 | 0.745 | 0.000 |
| **ULC179-bin7** | 0.754 | 0.731 | 0.785 | 0.752 | 0.757 | 0.000 | 0.820 | 0.000 |
| **ULC179-nobin** | 0.778 | 0.744 | 0.787 | 0.746 | 0.763 | 0.758 | 0.778 | 0.724 |
| **ULC186-bin1** | 0.821 | 0.751 | 0.756 | 0.759 | 0.741 | 0.730 | 0.742 | 0.742 |
| **ULC186-nobin** | 0.826 | 0.746 | 0.813 | 0.769 | 0.761 | 0.800 | 0.767 | 0.000 |
| **ULC187-bin1** | 0.725 | 0.742 | 0.743 | 0.734 | 0.734 | 0.725 | 0.695 | 0.000 |
| **ULC187-nobin** | 0.938 | 0.741 | 0.898 | 0.751 | 0.786 | 0.754 | 0.805 | 0.000 |

| | ULC082-nobin | ULC084-bin1 | ULC084-bin2 | ULC084-bin3 | ULC084-nobin | ULC129-bin1 | ULC129-nobin | ULC146-bin1 |
|---|---|---|---|---|---|---|---|---|
| **ULC335-bin1** | 0.716 | 0.717 | 0.724 | 0.728 | 0.717 | 0.736 | 0.744 | 0.701 |
| **ULC335-bin2** | 0.711 | 0.706 | 0.702 | 0.743 | 0.725 | 0.727 | 0.736 | 0.693 |
| **ULC335-bin3** | 0.734 | 0.718 | 0.711 | 0.713 | 0.721 | 0.714 | 0.735 | 0.729 |
| **ULC335-bin4** | 0.880 | 0.741 | 0.811 | 0.755 | 0.775 | 0.722 | 0.766 | 0.754 |
| **ULC335-nobin** | 0.841 | 0.756 | 0.795 | 0.745 | 0.795 | 0.733 | 0.795 | 0.841 |
| **ULC007-bin1** | 0.722 | 0.727 | 0.726 | 0.742 | 0.744 | 0.745 | 0.748 | 0.723 |
| **ULC007-bin2** | 0.698 | 0.690 | 0.000 | 0.761 | 0.000 | 0.762 | 0.727 | 0.000 |
| **ULC007-nobin** | 0.904 | 0.000 | 0.709 | 0.803 | 0.903 | 0.755 | 0.869 | 0.782 |
| **ULC027-bin1** | 0.725 | 0.730 | 0.730 | 0.742 | 0.750 | 0.788 | 0.817 | 0.739 |
| **ULC027-bin2** | 0.759 | 0.732 | 0.759 | 0.742 | 0.770 | 0.730 | 0.988 | 0.747 |
| **ULC027-bin3** | 0.762 | 0.729 | 0.761 | 0.740 | 0.988 | 0.730 | 0.911 | 0.752 |
| **ULC027-bin4** | 0.754 | 0.725 | 0.752 | 0.732 | 0.989 | 0.733 | 0.910 | 0.742 |
| **ULC027-nobin** | 0.802 | 0.984 | 0.765 | 0.752 | 0.869 | 0.798 | 0.935 | 0.876 |
| **ULC041-bin1** | 0.741 | 0.728 | 0.746 | 0.757 | 0.736 | 0.754 | 0.787 | 0.735 |
| **ULC041-bin2** | 0.762 | 0.000 | 0.742 | 0.729 | 0.711 | 0.753 | 0.790 | 0.716 |
| **ULC041-nobin** | 0.852 | | 0.000 | 0.783 | 0.893 | 0.752 | 0.898 | 0.798 |
| **ULC065-bin1** | 0.774 | 0.731 | 0.750 | 0.823 | 0.786 | 0.742 | 0.740 | 0.744 |
| **ULC065-bin2** | 0.761 | 0.740 | 0.755 | 0.746 | 0.746 | 0.735 | 0.749 | 0.762 |
| **ULC065-nobin** | 0.812 | 0.745 | 0.903 | 0.772 | 0.913 | 0.734 | 0.903 | 0.953 |
| **ULC066-bin1** | 0.722 | 0.733 | 0.727 | 0.729 | 0.723 | 0.738 | 0.731 | 0.723 |
| **ULC066-bin2** | 0.722 | 0.715 | 0.722 | 0.719 | 0.724 | 0.725 | 0.740 | 0.709 |
| **ULC066-bin3** | 0.750 | 0.993 | 0.747 | 0.729 | 0.968 | 0.714 | 0.987 | 0.750 |
| **ULC066-nobin** | 0.785 | 0.990 | 0.773 | 0.748 | 0.936 | 0.731 | 0.972 | 0.809 |
| **ULC068-bin1** | 0.728 | 0.740 | 0.719 | 0.735 | 0.724 | 0.738 | 0.733 | 0.727 |
| **ULC068-bin2** | 0.774 | 0.719 | 0.000 | 0.720 | 0.788 | 0.768 | 0.744 | 0.000 |
| **ULC068-nobin** | 0.789 | 0.757 | 0.882 | 0.745 | 0.795 | 0.744 | 0.916 | 0.800 |
| **ULC073-bin1** | 0.742 | 0.733 | 0.736 | 0.757 | 0.738 | 0.752 | 0.797 | 0.738 |
| **ULC073-bin2** | 0.752 | 0.997 | 0.726 | 0.741 | 0.972 | 0.729 | 0.993 | 0.751 |
| **ULC073-bin3** | 0.759 | 0.726 | 0.756 | 0.739 | 0.981 | 0.737 | 0.904 | 0.753 |
| **ULC073-bin4** | 0.763 | 0.718 | 0.751 | 0.766 | 0.794 | 0.729 | 0.876 | 0.750 |
| **ULC073-bin5** | 0.777 | 0.731 | 0.000 | 0.760 | 0.976 | 0.754 | 0.901 | 0.736 |
| **ULC073-bin6** | 0.789 | 0.726 | 0.714 | 0.000 | 0.987 | 0.708 | 0.965 | 0.761 |
| **ULC073-nobin** | 0.797 | 0.858 | 0.760 | 0.754 | 0.897 | 0.749 | 0.918 | 0.953 |
| **ULC077-bin1** | 0.739 | 0.733 | 0.729 | 0.738 | 0.732 | 0.744 | 0.733 | 0.727 |
| **ULC077-nobin** | 0.794 | 0.988 | 0.955 | 0.751 | 0.950 | 0.746 | 0.967 | 0.788 |
| **ULC082-bin1** | 0.764 | 0.725 | 0.748 | 0.986 | 0.929 | 0.743 | 0.746 | 0.744 |
| **ULC082-bin2** | 0.783 | 0.739 | 0.801 | 0.755 | 0.826 | 0.727 | 0.774 | 0.746 |
| **ULC082-bin3** | 0.782 | 0.747 | 0.740 | 0.737 | 0.762 | 0.736 | 0.764 | 0.789 |
| **ULC082-bin4** | 0.805 | 0.725 | 0.837 | 0.758 | 0.773 | 0.693 | 0.767 | 0.746 |
| **ULC082-bin5** | 0.788 | 0.742 | 0.696 | 0.000 | 0.730 | 0.873 | 0.773 | 0.744 |
| **ULC082-nobin** | | 0.752 | 0.807 | 0.841 | 0.812 | 0.730 | 0.792 | 0.840 |
| **ULC084-bin1** | 0.751 | | 0.732 | 0.734 | 0.794 | 0.726 | 0.992 | 0.752 |
| **ULC084-bin2** | 0.808 | 0.731 | | 0.745 | 0.769 | 0.738 | 0.760 | 0.745 |
| **ULC084-bin3** | 0.867 | 0.735 | 0.745 | | 0.787 | 0.740 | 0.757 | 0.747 |
| **ULC084-nobin** | 0.824 | 0.787 | 0.772 | 0.791 | | 0.741 | 0.917 | 0.788 |
| **ULC129-bin1** | 0.725 | 0.725 | 0.738 | 0.741 | 0.738 | | 0.863 | 0.736 |
| **ULC129-nobin** | 0.799 | 0.989 | 0.765 | 0.755 | 0.922 | 0.899 | | 0.943 |
| **ULC146-bin1** | 0.844 | 0.751 | 0.744 | 0.746 | 0.781 | 0.737 | 0.941 | |
| **ULC146-bin2** | 0.719 | 0.723 | 0.709 | 0.713 | 0.719 | 0.712 | 0.743 | 0.716 |
| **ULC146-bin3** | 0.763 | 0.734 | 0.756 | 0.744 | 0.988 | 0.733 | 0.914 | 0.753 |
| **ULC146-bin4** | 0.750 | 0.997 | 0.744 | 0.732 | 0.970 | 0.730 | 0.992 | 0.752 |
| **ULC146-bin5** | 0.836 | 0.731 | 0.807 | 0.745 | 0.774 | 0.722 | 0.760 | 0.749 |
| **ULC146-bin6** | 0.752 | 0.747 | 0.750 | 0.748 | 0.784 | 0.710 | 0.793 | 0.748 |
| **ULC146-bin7** | 0.753 | 0.723 | 0.766 | 0.740 | 0.988 | 0.000 | 0.923 | 0.765 |
| **ULC146-nobin** | 0.837 | 0.848 | 0.781 | 0.747 | 0.850 | 0.749 | 0.891 | 0.769 |
| **ULC165-bin1** | 0.759 | 0.740 | 0.753 | 0.740 | 0.748 | 0.723 | 0.749 | 0.760 |
| **ULC165-bin2** | 0.810 | 0.731 | 0.998 | 0.740 | 0.892 | 0.731 | 0.756 | 0.751 |
| **ULC165-bin3** | 0.846 | 0.750 | 0.746 | 0.735 | 0.789 | 0.735 | 0.935 | 0.996 |
| **ULC165-bin4** | 0.785 | 0.723 | 0.728 | 0.733 | 0.754 | 0.751 | 0.764 | 0.718 |
| **ULC165-nobin** | 0.838 | 0.750 | 0.804 | 0.745 | 0.870 | 0.749 | 0.880 | 0.985 |
| **ULC179-bin1** | 0.817 | 0.732 | 0.805 | 0.751 | 0.772 | 0.731 | 0.761 | 0.748 |
| **ULC179-bin2** | 0.822 | 0.730 | 0.764 | 0.740 | 0.771 | 0.728 | 0.767 | 0.750 |
| **ULC179-bin3** | 0.759 | 0.729 | 0.760 | 0.742 | 0.776 | 0.731 | 0.769 | 0.751 |
| **ULC179-bin4** | 0.739 | 0.729 | 0.731 | 0.728 | 0.739 | 0.732 | 0.740 | 0.739 |
| **ULC179-bin5** | 0.820 | 0.734 | 0.807 | 0.744 | 0.761 | 0.717 | 0.759 | 0.753 |
| **ULC179-bin6** | 0.762 | 0.729 | 0.752 | 0.725 | 0.778 | 0.727 | 0.764 | 0.747 |
| **ULC179-bin7** | 0.777 | 0.691 | 0.780 | 0.000 | 0.787 | 0.728 | 0.728 | 0.750 |
| **ULC179-nobin** | 0.789 | 0.745 | 0.777 | 0.748 | 0.796 | 0.738 | 0.780 | 0.771 |
| **ULC186-bin1** | 0.742 | 0.738 | 0.734 | 0.756 | 0.759 | 0.755 | 0.755 | 0.737 |
| **ULC186-nobin** | 0.810 | 0.772 | 0.757 | 0.765 | 0.839 | 0.752 | 0.827 | 0.816 |
| **ULC187-bin1** | 0.765 | 0.727 | 0.724 | 0.731 | 0.766 | 0.735 | 0.741 | 0.730 |
| **ULC187-nobin** | 0.805 | 0.820 | 0.954 | 0.751 | 0.958 | 0.749 | 0.899 | 0.765 |

| | ULC146-bin2 | ULC146-bin3 | ULC146-bin4 | ULC146-bin5 | ULC146-bin6 | ULC146-bin7 | ULC146-nobin | ULC165-bin1 |
|---|---|---|---|---|---|---|---|---|
| **ULC335-bin1** | 0.734 | 0.717 | 0.716 | 0.704 | 0.000 | 0.000 | 0.744 | 0.704 |
| **ULC335-bin2** | 0.773 | 0.000 | 0.717 | 0.709 | 0.000 | 0.000 | 0.762 | 0.725 |
| **ULC335-bin3** | 0.727 | 0.716 | 0.730 | 0.722 | 0.000 | 0.747 | 0.724 | 0.724 |
| **ULC335-bin4** | 0.738 | 0.775 | 0.742 | 0.854 | 0.772 | 0.746 | 0.817 | 0.768 |
| **ULC335-nobin** | 0.733 | 0.783 | 0.754 | 0.838 | 0.771 | 0.779 | 0.802 | 0.763 |
| **ULC007-bin1** | 0.725 | 0.717 | 0.719 | 0.721 | 0.737 | 0.726 | 0.745 | 0.715 |
| **ULC007-bin2** | 0.000 | 0.000 | 0.000 | 0.000 | 0.000 | 0.000 | 0.758 | 0.695 |
| **ULC007-nobin** | 0.000 | 0.822 | | 0.722 | 0.000 | 0.000 | 0.833 | 0.000 |
| **ULC027-bin1** | 0.719 | 0.727 | 0.723 | 0.726 | 0.730 | 0.000 | 0.736 | 0.723 |
| **ULC027-bin2** | 0.763 | 0.757 | 0.730 | 0.766 | 0.753 | 0.758 | 0.761 | 0.749 |
| **ULC027-bin3** | 0.717 | 0.997 | 0.733 | 0.767 | 0.828 | 0.991 | 0.810 | 0.754 |
| **ULC027-bin4** | 0.000 | 0.995 | 0.723 | 0.734 | 0.778 | 0.995 | 0.806 | 0.751 |
| **ULC027-nobin** | 0.728 | 0.861 | 0.985 | 0.774 | 0.959 | 0.875 | 0.951 | 0.756 |
| **ULC041-bin1** | 0.701 | 0.736 | 0.738 | 0.732 | 0.738 | 0.000 | 0.744 | 0.745 |
| **ULC041-bin2** | 0.000 | 0.723 | 0.702 | 0.740 | 0.000 | 0.685 | 0.721 | 0.756 |
| **ULC041-nobin** | 0.000 | | 0.000 | 0.722 | 0.000 | 0.719 | 0.823 | 0.753 |
| **ULC065-bin1** | 0.710 | 0.741 | 0.724 | 0.741 | 0.742 | 0.737 | 0.742 | 0.745 |
| **ULC065-bin2** | 0.707 | 0.752 | 0.736 | 0.752 | 0.745 | 0.740 | 0.755 | 0.998 |
| **ULC065-nobin** | 0.729 | 0.958 | 0.750 | 0.785 | 0.778 | 0.960 | 0.888 | 0.926 |
| **ULC066-bin1** | 0.735 | 0.717 | 0.715 | 0.701 | 0.717 | 0.000 | 0.739 | 0.707 |
| **ULC066-bin2** | 0.732 | 0.727 | 0.722 | 0.713 | 0.710 | 0.000 | 0.729 | 0.742 |
| **ULC066-bin3** | 0.000 | 0.757 | 0.994 | 0.748 | 0.713 | 0.000 | 0.914 | 0.725 |
| **ULC066-nobin** | 0.732 | 0.947 | 0.991 | 0.776 | 0.777 | 0.948 | 0.872 | 0.754 |
| **ULC068-bin1** | 0.710 | 0.727 | 0.723 | 0.713 | 0.724 | 0.708 | 0.740 | 0.717 |
| **ULC068-bin2** | 0.000 | 0.772 | 0.708 | 0.000 | 0.724 | 0.000 | 0.744 | 0.000 |
| **ULC068-nobin** | 0.779 | 0.774 | 0.756 | 0.780 | 0.764 | 0.767 | 0.908 | 0.981 |
| **ULC073-bin1** | 0.683 | 0.737 | 0.726 | 0.736 | 0.743 | 0.700 | 0.742 | 0.743 |
| **ULC073-bin2** | 0.729 | 0.733 | 0.998 | 0.729 | 0.759 | 0.732 | 0.887 | 0.740 |
| **ULC073-bin3** | 0.714 | 0.990 | 0.728 | 0.756 | 0.818 | 0.991 | 0.845 | 0.752 |
| **ULC073-bin4** | 0.000 | 0.796 | 0.749 | 0.754 | 0.841 | 0.784 | 0.925 | 0.745 |
| **ULC073-bin5** | 0.000 | 0.000 | 0.732 | 0.000 | 0.000 | 0.730 | 0.966 | 0.000 |
| **ULC073-bin6** | 0.000 | 0.990 | 0.000 | 0.765 | 0.781 | 0.849 | 0.943 | 0.732 |
| **ULC073-nobin** | 0.828 | 0.907 | 0.886 | 0.763 | 0.951 | 0.911 | 0.942 | 0.754 |
| **ULC077-bin1** | 0.719 | 0.713 | 0.721 | 0.718 | 0.721 | 0.723 | 0.746 | 0.727 |
| **ULC077-nobin** | 0.717 | 0.905 | 0.991 | 0.790 | 0.800 | 0.907 | 0.888 | 0.988 |
| **ULC082-bin1** | 0.000 | 0.741 | 0.720 | 0.733 | 0.751 | 0.732 | 0.752 | 0.741 |
| **ULC082-bin2** | 0.747 | 0.758 | 0.739 | 0.826 | 0.751 | 0.749 | 0.768 | 0.745 |
| **ULC082-bin3** | 0.724 | 0.754 | 0.743 | 0.745 | 0.753 | 0.746 | 0.757 | 0.761 |
| **ULC082-bin4** | 0.716 | 0.765 | 0.744 | 0.845 | 0.755 | 0.760 | 0.799 | 0.761 |
| **ULC082-bin5** | 0.000 | 0.726 | 0.728 | 0.731 | 0.744 | 0.000 | 0.845 | 0.766 |
| **ULC082-nobin** | 0.716 | 0.763 | 0.752 | 0.826 | 0.753 | 0.756 | 0.832 | 0.759 |
| **ULC084-bin1** | 0.723 | 0.734 | 0.997 | 0.730 | 0.747 | 0.723 | 0.867 | 0.741 |
| **ULC084-bin2** | 0.709 | 0.757 | 0.744 | 0.806 | 0.747 | 0.763 | 0.782 | 0.753 |
| **ULC084-bin3** | 0.713 | 0.747 | 0.733 | 0.745 | 0.748 | 0.738 | 0.747 | 0.743 |
| **ULC084-nobin** | 0.716 | 0.988 | 0.969 | 0.776 | 0.790 | 0.984 | 0.862 | 0.756 |
| **ULC129-bin1** | 0.712 | 0.733 | 0.731 | 0.721 | 0.710 | 0.694 | 0.743 | 0.724 |
| **ULC129-nobin** | 0.748 | 0.909 | 0.989 | 0.766 | 0.792 | 0.917 | 0.901 | 0.756 |
| **ULC146-bin1** | 0.716 | 0.750 | 0.752 | 0.745 | 0.744 | 0.765 | 0.773 | 0.759 |
| **ULC146-bin2** | | 0.715 | 0.707 | 0.712 | 0.000 | 0.000 | 0.802 | 0.705 |
| **ULC146-bin3** | 0.715 | | 0.737 | 0.759 | 0.788 | 0.797 | 0.786 | 0.750 |
| **ULC146-bin4** | 0.707 | 0.738 | | 0.734 | 0.736 | 0.715 | 0.790 | 0.742 |
| **ULC146-bin5** | 0.712 | 0.762 | 0.737 | | 0.757 | 0.760 | 0.774 | 0.759 |
| **ULC146-bin6** | 0.000 | 0.791 | 0.736 | 0.758 | | 0.765 | 0.810 | 0.743 |
| **ULC146-bin7** | 0.000 | 0.825 | 0.715 | 0.763 | 0.765 | | 0.780 | 0.753 |
| **ULC146-nobin** | 0.835 | 0.788 | 0.795 | 0.777 | 0.815 | 0.779 | | 0.756 |
| **ULC165-bin1** | 0.705 | 0.751 | 0.743 | 0.757 | 0.738 | 0.749 | 0.754 | |
| **ULC165-bin2** | 0.734 | 0.756 | 0.731 | 0.804 | 0.750 | 0.759 | 0.785 | 0.748 |
| **ULC165-bin3** | 0.699 | 0.748 | 0.752 | 0.750 | 0.751 | 0.763 | 0.795 | 0.760 |
| **ULC165-bin4** | 0.714 | 0.728 | 0.740 | 0.000 | 0.732 | 0.000 | 0.797 | 0.735 |
| **ULC165-nobin** | 0.747 | 0.849 | 0.751 | 0.773 | 0.886 | 0.839 | 0.873 | 0.774 |
| **ULC179-bin1** | 0.719 | 0.758 | 0.731 | 0.820 | 0.757 | 0.756 | 0.795 | 0.754 |
| **ULC179-bin2** | 0.706 | 0.764 | 0.734 | 0.774 | 0.760 | 0.754 | 0.907 | 0.749 |
| **ULC179-bin3** | 0.716 | 0.774 | 0.733 | 0.760 | 0.769 | 0.764 | 0.773 | 0.754 |
| **ULC179-bin4** | 0.724 | 0.732 | 0.727 | 0.730 | 0.719 | 0.752 | 0.741 | 0.730 |
| **ULC179-bin5** | 0.732 | 0.754 | 0.742 | 0.826 | 0.750 | 0.751 | 0.802 | 0.747 |
| **ULC179-bin6** | 0.000 | 0.760 | 0.739 | 0.762 | 0.746 | 0.744 | 0.843 | 0.763 |
| **ULC179-bin7** | 0.000 | 0.752 | 0.000 | 0.759 | 0.740 | 0.768 | 0.824 | 0.721 |
| **ULC179-nobin** | 0.728 | 0.770 | 0.747 | 0.783 | 0.769 | 0.751 | 0.860 | 0.758 |
| **ULC186-bin1** | 0.717 | 0.732 | 0.732 | 0.742 | 0.740 | 0.717 | 0.746 | 0.738 |
| **ULC186-nobin** | 0.751 | 0.777 | 0.765 | 0.765 | 0.805 | 0.760 | 0.824 | 0.768 |
| **ULC187-bin1** | 0.721 | 0.724 | 0.727 | 0.704 | 0.730 | 0.704 | 0.741 | 0.729 |
| **ULC187-nobin** | 0.759 | 0.975 | 0.819 | 0.800 | 0.836 | 0.980 | 0.900 | 0.762 |

| | ULC165-bin2 | ULC165-bin3 | ULC165-bin4 | ULC165-nobin | ULC179-bin1 | ULC179-bin2 | ULC179-bin3 | ULC179-bin4 |
|---|---|---|---|---|---|---|---|---|
| **ULC335-bin1** | 0.722 | 0.000 | 0.747 | 0.742 | 0.715 | 0.706 | 0.714 | 0.725 |
| **ULC335-bin2** | 0.703 | 0.000 | 0.709 | 0.766 | 0.717 | 0.000 | 0.700 | 0.726 |
| **ULC335-bin3** | 0.710 | 0.749 | 0.717 | 0.726 | 0.721 | 0.719 | 0.713 | 0.740 |
| **ULC335-bin4** | 0.813 | 0.754 | 0.690 | 0.781 | 0.832 | 0.786 | 0.770 | 0.749 |
| **ULC335-nobin** | 0.796 | 0.840 | 0.740 | 0.822 | 0.816 | 0.778 | 0.766 | 0.741 |
| **ULC007-bin1** | 0.725 | 0.728 | 0.755 | 0.755 | 0.724 | 0.721 | 0.728 | 0.723 |
| **ULC007-bin2** | 0.000 | 0.000 | 0.776 | 0.777 | 0.000 | 0.000 | 0.000 | 0.000 |
| **ULC007-nobin** | 0.692 | 0.000 | 0.808 | 0.793 | 0.000 | 0.737 | 0.815 | 0.000 |
| **ULC027-bin1** | 0.733 | 0.723 | 0.743 | 0.749 | 0.719 | 0.730 | 0.729 | 0.720 |
| **ULC027-bin2** | 0.757 | 0.755 | 0.701 | 0.762 | 0.761 | 0.765 | 0.760 | 0.730 |
| **ULC027-bin3** | 0.760 | 0.753 | 0.717 | 0.865 | 0.766 | 0.765 | 0.776 | 0.739 |
| **ULC027-bin4** | 0.753 | 0.745 | 0.000 | 0.836 | 0.747 | 0.765 | 0.773 | 0.734 |
| **ULC027-nobin** | 0.766 | 0.865 | 0.761 | 0.876 | 0.777 | 0.815 | 0.775 | 0.738 |
| **ULC041-bin1** | 0.738 | 0.741 | 0.746 | 0.747 | 0.746 | 0.739 | 0.729 | 0.736 |
| **ULC041-bin2** | 0.740 | 0.760 | 0.751 | 0.778 | 0.737 | 0.730 | 0.746 | 0.727 |
| **ULC041-nobin** | 0.000 | 0.835 | 0.000 | 0.818 | 0.720 | 0.721 | 0.799 | 0.000 |
| **ULC065-bin1** | 0.746 | 0.742 | 0.731 | 0.744 | 0.743 | 0.742 | 0.734 | 0.719 |
| **ULC065-bin2** | 0.755 | 0.762 | 0.733 | 0.810 | 0.756 | 0.755 | 0.756 | 0.735 |
| **ULC065-nobin** | 0.897 | 0.944 | 0.765 | 0.907 | 0.780 | 0.774 | 0.774 | 0.739 |
| **ULC066-bin1** | 0.730 | 0.728 | 0.745 | 0.744 | 0.711 | 0.720 | 0.723 | 0.718 |
| **ULC066-bin2** | 0.708 | 0.750 | 0.727 | 0.713 | 0.704 | 0.724 | 0.742 | 0.726 |
| **ULC066-bin3** | 0.744 | 0.744 | 0.698 | 0.747 | 0.744 | 0.753 | 0.730 | 0.736 |
| **ULC066-nobin** | 0.773 | 0.797 | 0.724 | 0.849 | 0.778 | 0.788 | 0.775 | 0.739 |
| **ULC068-bin1** | 0.710 | 0.734 | 0.738 | 0.746 | 0.711 | 0.725 | 0.720 | 0.726 |
| **ULC068-bin2** | 0.000 | 0.000 | 0.708 | 0.763 | 0.000 | 0.000 | 0.755 | 0.000 |
| **ULC068-nobin** | 0.881 | 0.803 | 0.742 | 0.830 | 0.782 | 0.776 | 0.772 | 0.733 |
| **ULC073-bin1** | 0.735 | 0.727 | 0.753 | 0.749 | 0.742 | 0.741 | 0.737 | 0.733 |
| **ULC073-bin2** | 0.734 | 0.755 | 0.740 | 0.751 | 0.736 | 0.731 | 0.733 | 0.737 |
| **ULC073-bin3** | 0.759 | 0.751 | 0.000 | 0.865 | 0.761 | 0.765 | 0.779 | 0.739 |
| **ULC073-bin4** | 0.754 | 0.742 | 0.000 | 0.889 | 0.760 | 0.765 | 0.774 | 0.763 |
| **ULC073-bin5** | 0.000 | 0.727 | 0.770 | 0.750 | 0.734 | 0.721 | 0.000 | 0.730 |
| **ULC073-bin6** | 0.760 | 0.750 | 0.000 | 0.948 | 0.756 | 0.747 | 0.791 | 0.000 |
| **ULC073-nobin** | 0.761 | 0.953 | 0.757 | 0.902 | 0.762 | 0.770 | 0.775 | 0.737 |
| **ULC077-bin1** | 0.728 | 0.725 | 0.753 | 0.754 | 0.726 | 0.732 | 0.720 | 0.732 |
| **ULC077-nobin** | 0.956 | 0.785 | 0.745 | 0.820 | 0.792 | 0.770 | 0.769 | 0.740 |
| **ULC082-bin1** | 0.745 | 0.738 | 0.736 | 0.747 | 0.740 | 0.743 | 0.734 | 0.728 |
| **ULC082-bin2** | 0.809 | 0.749 | 0.000 | 0.762 | 0.802 | 0.762 | 0.758 | 0.724 |
| **ULC082-bin3** | 0.745 | 0.809 | 0.706 | 0.766 | 0.749 | 0.744 | 0.740 | 0.745 |
| **ULC082-bin4** | 0.823 | 0.741 | 0.685 | 0.768 | 0.827 | 0.772 | 0.766 | 0.752 |
| **ULC082-bin5** | 0.000 | 0.735 | 0.000 | 0.768 | 0.725 | 0.731 | 0.000 | 0.873 |
| **ULC082-nobin** | 0.808 | 0.837 | 0.782 | 0.833 | 0.816 | 0.814 | 0.761 | 0.742 |
| **ULC084-bin1** | 0.731 | 0.750 | 0.723 | 0.750 | 0.729 | 0.732 | 0.728 | 0.729 |
| **ULC084-bin2** | 0.998 | 0.743 | 0.696 | 0.810 | 0.808 | 0.764 | 0.760 | 0.733 |
| **ULC084-bin3** | 0.739 | 0.731 | 0.733 | 0.744 | 0.750 | 0.742 | 0.741 | 0.729 |
| **ULC084-nobin** | 0.888 | 0.794 | 0.754 | 0.878 | 0.777 | 0.778 | 0.782 | 0.745 |
| **ULC129-bin1** | 0.730 | 0.732 | 0.748 | 0.746 | 0.737 | 0.724 | 0.729 | 0.732 |
| **ULC129-nobin** | 0.764 | 0.935 | 0.762 | 0.891 | 0.768 | 0.776 | 0.775 | 0.745 |
| **ULC146-bin1** | 0.750 | 0.992 | 0.715 | 0.989 | 0.747 | 0.750 | 0.750 | 0.737 |
| **ULC146-bin2** | 0.734 | 0.699 | 0.714 | 0.747 | 0.717 | 0.706 | 0.716 | 0.723 |
| **ULC146-bin3** | 0.756 | 0.746 | 0.728 | 0.854 | 0.758 | 0.764 | 0.775 | 0.729 |
| **ULC146-bin4** | 0.731 | 0.751 | 0.730 | 0.751 | 0.732 | 0.733 | 0.733 | 0.727 |
| **ULC146-bin5** | 0.805 | 0.750 | 0.000 | 0.777 | 0.825 | 0.776 | 0.763 | 0.730 |
| **ULC146-bin6** | 0.750 | 0.743 | 0.000 | 0.891 | 0.760 | 0.762 | 0.773 | 0.719 |
| **ULC146-bin7** | 0.761 | 0.763 | 0.000 | 0.853 | 0.757 | 0.760 | 0.767 | 0.752 |
| **ULC146-nobin** | 0.783 | 0.781 | 0.790 | 0.869 | 0.793 | 0.900 | 0.778 | 0.749 |
| **ULC165-bin1** | 0.748 | 0.757 | 0.735 | 0.761 | 0.753 | 0.750 | 0.754 | 0.729 |
| **ULC165-bin2** | | 0.746 | 0.735 | 0.767 | 0.807 | 0.764 | 0.758 | 0.736 |
| **ULC165-bin3** | 0.748 | | 0.714 | 0.778 | 0.748 | 0.749 | 0.755 | 0.728 |
| **ULC165-bin4** | 0.743 | 0.714 | | 0.801 | 0.764 | 0.738 | 0.738 | 0.726 |
| **ULC165-nobin** | 0.767 | 0.776 | 0.828 | | 0.769 | 0.813 | 0.781 | 0.733 |
| **ULC179-bin1** | 0.805 | 0.747 | 0.764 | 0.767 | | 0.766 | 0.759 | 0.738 |
| **ULC179-bin2** | 0.765 | 0.747 | 0.719 | 0.814 | 0.768 | | 0.761 | 0.729 |
| **ULC179-bin3** | 0.757 | 0.751 | 0.738 | 0.777 | 0.760 | 0.762 | | 0.736 |
| **ULC179-bin4** | 0.736 | 0.728 | 0.726 | 0.736 | 0.740 | 0.729 | 0.736 | |
| **ULC179-bin5** | 0.798 | 0.746 | 0.000 | 0.766 | 0.815 | 0.773 | 0.762 | 0.742 |
| **ULC179-bin6** | 0.752 | 0.752 | 0.000 | 0.763 | 0.768 | 0.769 | 0.758 | 0.745 |
| **ULC179-bin7** | 0.762 | 0.774 | 0.000 | 0.771 | 0.768 | 0.745 | 0.845 | 0.750 |
| **ULC179-nobin** | 0.781 | 0.763 | 0.750 | 0.790 | 0.777 | 0.785 | 0.787 | 0.808 |
| **ULC186-bin1** | 0.737 | 0.740 | 0.751 | 0.750 | 0.739 | 0.735 | 0.736 | 0.740 |
| **ULC186-nobin** | 0.764 | 0.811 | 0.752 | 0.822 | 0.759 | 0.771 | 0.765 | 0.747 |
| **ULC187-bin1** | 0.723 | 0.731 | 0.743 | 0.739 | 0.723 | 0.718 | 0.716 | 0.726 |
| **ULC187-nobin** | 0.953 | 0.762 | 0.755 | 0.864 | 0.796 | 0.784 | 0.786 | 0.738 |

| | ULC179-bin5 | ULC179-bin6 | ULC179-bin7 | ULC179-nobin | ULC186-bin1 | ULC186-nobin | ULC187-bin1 | ULC187-nobin |
|---|---|---|---|---|---|---|---|---|
| **ULC335-bin1** | 0.693 | 0.000 | 0.000 | 0.746 | 0.738 | 0.861 | 0.748 | 0.734 |
| **ULC335-bin2** | 0.000 | 0.000 | 0.000 | 0.722 | 0.723 | 0.749 | 0.730 | 0.718 |
| **ULC335-bin3** | 0.717 | 0.725 | 0.000 | 0.733 | 0.726 | 0.725 | 0.717 | 0.712 |
| **ULC335-bin4** | 0.835 | 0.765 | 0.746 | 0.789 | 0.750 | 0.769 | 0.718 | 0.804 |
| **ULC335-nobin** | 0.811 | 0.759 | 0.760 | 0.778 | 0.746 | 0.885 | 0.743 | 0.790 |
| **ULC007-bin1** | 0.727 | 0.718 | 0.000 | 0.743 | 0.750 | 0.749 | 0.741 | 0.759 |
| **ULC007-bin2** | 0.000 | 0.000 | 0.000 | 0.774 | 0.774 | 0.786 | 0.701 | 0.000 |
| **ULC007-nobin** | 0.000 | 0.000 | 0.000 | 0.835 | 0.791 | 0.906 | 0.788 | 0.892 |
| **ULC027-bin1** | 0.726 | 0.740 | 0.703 | 0.743 | 0.758 | 0.748 | 0.744 | 0.761 |
| **ULC027-bin2** | 0.762 | 0.756 | 0.761 | 0.758 | 0.742 | 0.759 | 0.725 | 0.771 |
| **ULC027-bin3** | 0.763 | 0.769 | 0.766 | 0.763 | 0.727 | 0.754 | 0.723 | 0.977 |
| **ULC027-bin4** | 0.753 | 0.759 | 0.000 | 0.754 | 0.731 | 0.737 | 0.000 | 0.970 |
| **ULC027-nobin** | 0.779 | 0.772 | 0.755 | 0.781 | 0.747 | 0.785 | 0.739 | 0.912 |
| **ULC041-bin1** | 0.731 | 0.725 | 0.000 | 0.742 | 0.842 | 0.838 | 0.737 | 0.743 |
| **ULC041-bin2** | 0.000 | 0.000 | 0.000 | 0.746 | 0.839 | 0.851 | 0.739 | 0.727 |
| **ULC041-nobin** | 0.000 | 0.710 | 0.000 | 0.839 | 0.862 | 0.881 | 0.771 | 0.879 |
| **ULC065-bin1** | 0.738 | 0.735 | 0.000 | 0.746 | 0.756 | 0.750 | 0.732 | 0.742 |
| **ULC065-bin2** | 0.752 | 0.763 | 0.000 | 0.751 | 0.735 | 0.762 | 0.706 | 0.758 |
| **ULC065-nobin** | 0.775 | 0.768 | 0.759 | 0.778 | 0.744 | 0.804 | 0.730 | 0.932 |
| **ULC066-bin1** | 0.697 | 0.714 | 0.000 | 0.742 | 0.738 | 0.743 | 0.798 | 0.798 |
| **ULC066-bin2** | 0.000 | 0.000 | 0.000 | 0.733 | 0.713 | 0.734 | 0.734 | 0.747 |
| **ULC066-bin3** | 0.756 | 0.723 | 0.000 | 0.715 | 0.732 | 0.740 | 0.727 | 0.825 |
| **ULC066-nobin** | 0.776 | 0.773 | 0.752 | 0.780 | 0.739 | 0.793 | 0.746 | 0.899 |
| **ULC068-bin1** | 0.717 | 0.731 | 0.000 | 0.741 | 0.737 | 0.732 | 0.797 | 0.804 |
| **ULC068-bin2** | 0.000 | 0.000 | 0.000 | 0.749 | 0.718 | 0.811 | 0.886 | 0.833 |
| **ULC068-nobin** | 0.778 | 0.767 | 0.796 | 0.783 | 0.745 | 0.808 | 0.786 | 0.900 |
| **ULC073-bin1** | 0.732 | 0.734 | 0.000 | 0.744 | 0.843 | 0.841 | 0.737 | 0.742 |
| **ULC073-bin2** | 0.749 | 0.737 | 0.725 | 0.742 | 0.731 | 0.766 | 0.728 | 0.819 |
| **ULC073-bin3** | 0.760 | 0.763 | 0.741 | 0.758 | 0.729 | 0.755 | 0.730 | 0.970 |
| **ULC073-bin4** | 0.761 | 0.760 | 0.789 | 0.774 | 0.746 | 0.713 | 0.705 | 0.925 |
| **ULC073-bin5** | 0.000 | 0.000 | 0.000 | 0.731 | 0.852 | 0.829 | 0.741 | 0.000 |
| **ULC073-bin6** | 0.791 | 0.754 | 0.777 | 0.795 | 0.749 | 0.836 | 0.000 | 0.991 |
| **ULC073-nobin** | 0.759 | 0.764 | 0.753 | 0.775 | 0.824 | 0.815 | 0.727 | 0.920 |
| **ULC077-bin1** | 0.716 | 0.716 | 0.731 | 0.739 | 0.751 | 0.748 | 0.742 | 0.737 |
| **ULC077-nobin** | 0.786 | 0.766 | 0.774 | 0.784 | 0.755 | 0.802 | 0.742 | 0.894 |
| **ULC082-bin1** | 0.736 | 0.722 | 0.752 | 0.748 | 0.758 | 0.785 | 0.734 | 0.757 |
| **ULC082-bin2** | 0.811 | 0.756 | 0.757 | 0.758 | 0.740 | 0.762 | 0.737 | 0.777 |
| **ULC082-bin3** | 0.745 | 0.742 | 0.000 | 0.750 | 0.731 | 0.789 | 0.723 | 0.751 |
| **ULC082-bin4** | 0.834 | 0.745 | 0.820 | 0.778 | 0.734 | 0.727 | 0.695 | 0.801 |
| **ULC082-bin5** | 0.000 | 0.000 | 0.000 | 0.730 | 0.742 | 0.000 | 0.000 | 0.000 |
| **ULC082-nobin** | 0.815 | 0.759 | 0.779 | 0.782 | 0.744 | 0.800 | 0.762 | 0.798 |
| **ULC084-bin1** | 0.734 | 0.729 | 0.691 | 0.739 | 0.739 | 0.768 | 0.729 | 0.816 |
| **ULC084-bin2** | 0.806 | 0.749 | 0.778 | 0.772 | 0.734 | 0.753 | 0.724 | 0.956 |
| **ULC084-bin3** | 0.744 | 0.725 | 0.000 | 0.745 | 0.756 | 0.763 | 0.731 | 0.745 |
| **ULC084-nobin** | 0.767 | 0.785 | 0.774 | 0.794 | 0.756 | 0.831 | 0.763 | 0.955 |
| **ULC129-bin1** | 0.715 | 0.727 | 0.728 | 0.736 | 0.755 | 0.748 | 0.735 | 0.746 |
| **ULC129-nobin** | 0.770 | 0.767 | 0.760 | 0.779 | 0.757 | 0.808 | 0.744 | 0.892 |
| **ULC146-bin1** | 0.748 | 0.744 | 0.750 | 0.765 | 0.736 | 0.806 | 0.729 | 0.756 |
| **ULC146-bin2** | 0.732 | 0.000 | 0.000 | 0.728 | 0.717 | 0.738 | 0.722 | 0.764 |
| **ULC146-bin3** | 0.754 | 0.760 | 0.742 | 0.764 | 0.730 | 0.779 | 0.726 | 0.973 |
| **ULC146-bin4** | 0.740 | 0.739 | 0.000 | 0.745 | 0.731 | 0.758 | 0.725 | 0.817 |
| **ULC146-bin5** | 0.827 | 0.763 | 0.759 | 0.781 | 0.747 | 0.768 | 0.706 | 0.795 |
| **ULC146-bin6** | 0.751 | 0.746 | 0.728 | 0.766 | 0.736 | 0.774 | 0.742 | 0.834 |
| **ULC146-bin7** | 0.747 | 0.744 | 0.768 | 0.747 | 0.726 | 0.759 | 0.711 | 0.974 |
| **ULC146-nobin** | 0.803 | 0.830 | 0.812 | 0.853 | 0.746 | 0.812 | 0.748 | 0.891 |
| **ULC165-bin1** | 0.746 | 0.763 | 0.721 | 0.754 | 0.738 | 0.751 | 0.729 | 0.754 |
| **ULC165-bin2** | 0.797 | 0.753 | 0.762 | 0.779 | 0.737 | 0.762 | 0.722 | 0.955 |
| **ULC165-bin3** | 0.744 | 0.751 | 0.758 | 0.764 | 0.740 | 0.803 | 0.731 | 0.757 |
| **ULC165-bin4** | 0.000 | 0.000 | 0.000 | 0.745 | 0.754 | 0.743 | 0.745 | 0.723 |
| **ULC165-nobin** | 0.770 | 0.773 | 0.771 | 0.784 | 0.751 | 0.805 | 0.742 | 0.852 |
| **ULC179-bin1** | 0.814 | 0.765 | 0.756 | 0.771 | 0.738 | 0.757 | 0.720 | 0.791 |
| **ULC179-bin2** | 0.774 | 0.765 | 0.735 | 0.779 | 0.732 | 0.764 | 0.717 | 0.775 |
| **ULC179-bin3** | 0.760 | 0.760 | 0.846 | 0.773 | 0.736 | 0.756 | 0.716 | 0.779 |
| **ULC179-bin4** | 0.742 | 0.729 | 0.750 | 0.782 | 0.741 | 0.746 | 0.726 | 0.740 |
| **ULC179-bin5** | | 0.766 | 0.765 | 0.778 | 0.723 | 0.753 | 0.711 | 0.789 |
| **ULC179-bin6** | 0.768 | | 0.768 | 0.785 | 0.741 | 0.765 | 0.705 | 0.773 |
| **ULC179-bin7** | 0.767 | 0.768 | | 0.815 | 0.000 | 0.000 | 0.757 | 0.730 |
| **ULC179-nobin** | 0.778 | 0.786 | 0.833 | | 0.741 | 0.798 | 0.744 | 0.790 |
| **ULC186-bin1** | 0.725 | 0.736 | 0.000 | 0.742 | | 0.879 | 0.741 | 0.797 |
| **ULC186-nobin** | 0.756 | 0.770 | 0.000 | 0.792 | 0.923 | | 0.766 | 0.841 |
| **ULC187-bin1** | 0.708 | 0.705 | 0.757 | 0.738 | 0.741 | 0.761 | | 0.870 |
| **ULC187-nobin** | 0.793 | 0.787 | 0.734 | 0.790 | 0.785 | 0.830 | 0.889 | |

| | ULC335-bin1 | ULC335-bin2 | ULC335-bin3 | ULC335-bin4 | ULC335-nobin | ULC007-bin1 | ULC007-bin2 | ULC007-nobin |
|---|---|---|---|---|---|---|---|---|
| **ULC335-bin1** | | 0.002 | 0.000 | 0.000 | 0.008 | 0.040 | 0.000 | 0.001 |
| **ULC335-bin2** | 0.001 | | 0.004 | | 0.004 | 0.000 | | |
| **ULC335-bin3** | 0.000 | 0.004 | | 0.000 | 0.001 | 0.001 | | |
| **ULC335-bin4** | 0.000 | | 0.001 | | 0.027 | 0.003 | | |
| **ULC335-nobin** | 0.002 | 0.002 | 0.000 | 0.005 | | 0.001 | 0.000 | 0.001 |
| **ULC007-bin1** | 0.035 | 0.000 | 0.001 | 0.001 | 0.002 | | 0.001 | 0.003 |
| **ULC007-bin2** | 0.004 | | | | | 0.015 | | 0.008 |
| **ULC007-nobin** | 0.012 | | | | 0.018 | 0.071 | 0.016 | |
| **ULC027-bin1** | 0.023 | 0.000 | 0.000 | 0.000 | 0.003 | 0.039 | 0.001 | 0.000 |
| **ULC027-bin2** | 0.000 | 0.000 | 0.001 | 0.015 | 0.031 | 0.001 | | 0.000 |
| **ULC027-bin3** | 0.000 | 0.000 | 0.000 | 0.010 | 0.046 | 0.001 | | 0.000 |
| **ULC027-bin4** | | | 0.000 | 0.007 | 0.042 | 0.001 | | |
| **ULC027-nobin** | 0.001 | 0.000 | 0.000 | 0.007 | 0.032 | 0.002 | 0.000 | 0.001 |
| **ULC041-bin1** | 0.023 | 0.000 | 0.001 | 0.001 | 0.005 | 0.044 | 0.000 | 0.001 |
| **ULC041-bin2** | 0.006 | | 0.000 | 0.000 | 0.002 | 0.019 | 0.004 | 0.000 |
| **ULC041-nobin** | 0.004 | | | | 0.010 | 0.013 | | 0.024 |
| **ULC065-bin1** | 0.007 | | 0.001 | 0.002 | 0.010 | 0.017 | 0.000 | 0.000 |
| **ULC065-bin2** | 0.000 | 0.000 | 0.001 | 0.006 | 0.028 | 0.001 | 0.000 | |
| **ULC065-nobin** | 0.000 | 0.000 | 0.000 | 0.010 | 0.059 | 0.001 | 0.000 | 0.001 |
| **ULC066-bin1** | 0.026 | 0.000 | 0.001 | 0.000 | 0.001 | 0.030 | 0.001 | 0.001 |
| **ULC066-bin2** | 0.001 | 0.011 | 0.033 | 0.000 | 0.014 | 0.000 | | 0.000 |
| **ULC066-bin3** | 0.001 | | 0.001 | 0.002 | 0.030 | 0.001 | | |
| **ULC066-nobin** | 0.001 | 0.001 | 0.002 | 0.008 | 0.034 | 0.001 | 0.000 | 0.001 |
| **ULC068-bin1** | 0.028 | 0.000 | 0.001 | 0.000 | 0.001 | 0.036 | 0.001 | 0.001 |
| **ULC068-bin2** | 0.007 | | 0.001 | | 0.006 | 0.005 | | 0.006 |
| **ULC068-nobin** | 0.001 | 0.008 | 0.001 | 0.012 | 0.042 | 0.002 | | 0.001 |
| **ULC073-bin1** | 0.023 | 0.000 | 0.001 | 0.001 | 0.006 | 0.041 | 0.000 | 0.001 |
| **ULC073-bin2** | 0.001 | 0.001 | 0.001 | 0.001 | 0.025 | 0.001 | | 0.000 |
| **ULC073-bin3** | 0.000 | 0.000 | 0.000 | 0.011 | 0.049 | 0.001 | | 0.000 |
| **ULC073-bin4** | | 0.000 | | 0.010 | 0.035 | 0.001 | 0.000 | |
| **ULC073-bin5** | 0.011 | | | | 0.002 | 0.023 | 0.006 | 0.001 |
| **ULC073-bin6** | | | | 0.001 | 0.014 | | | 0.003 |
| **ULC073-nobin** | 0.001 | 0.000 | 0.000 | 0.008 | 0.042 | 0.002 | 0.000 | 0.001 |
| **ULC077-bin1** | 0.035 | 0.000 | 0.000 | 0.000 | 0.002 | 0.078 | 0.002 | 0.002 |
| **ULC077-nobin** | 0.002 | 0.000 | 0.001 | 0.008 | 0.034 | 0.003 | 0.002 | 0.005 |
| **ULC082-bin1** | 0.010 | | 0.001 | 0.003 | 0.009 | 0.019 | 0.001 | 0.001 |
| **ULC082-bin2** | 0.001 | 0.000 | 0.001 | 0.028 | 0.086 | 0.001 | | |
| **ULC082-bin3** | 0.001 | 0.000 | 0.001 | 0.004 | 0.029 | 0.002 | | |
| **ULC082-bin4** | | 0.001 | | 0.117 | 0.206 | 0.002 | | |
| **ULC082-bin5** | | | | 0.001 | 0.002 | | | |
| **ULC082-nobin** | 0.000 | 0.000 | 0.000 | 0.024 | 0.087 | 0.001 | 0.000 | 0.001 |
| **ULC084-bin1** | 0.002 | 0.000 | 0.001 | 0.002 | 0.026 | 0.001 | 0.000 | |
| **ULC084-bin2** | 0.000 | 0.000 | 0.001 | 0.060 | 0.108 | 0.001 | | 0.000 |
| **ULC084-bin3** | 0.009 | 0.000 | 0.000 | 0.002 | 0.008 | 0.015 | 0.001 | 0.001 |
| **ULC084-nobin** | 0.000 | 0.000 | 0.001 | 0.007 | 0.052 | 0.001 | 0.000 | 0.001 |
| **ULC129-bin1** | 0.023 | 0.000 | 0.000 | 0.000 | 0.003 | 0.035 | 0.001 | 0.001 |
| **ULC129-nobin** | 0.001 | 0.002 | 0.002 | 0.005 | 0.029 | 0.002 | 0.000 | 0.001 |
| **ULC146-bin1** | 0.000 | 0.000 | 0.001 | 0.005 | 0.140 | 0.001 | | 0.000 |
| **ULC146-bin2** | 0.000 | 0.097 | 0.004 | 0.000 | 0.003 | 0.000 | | |
| **ULC146-bin3** | 0.000 | | 0.000 | 0.011 | 0.050 | 0.001 | | 0.000 |
| **ULC146-bin4** | 0.001 | 0.000 | 0.001 | 0.001 | 0.024 | 0.001 | | |
| **ULC146-bin5** | 0.000 | 0.000 | 0.001 | 0.097 | 0.156 | 0.001 | | 0.000 |
| **ULC146-bin6** | | | | 0.008 | 0.033 | 0.000 | | |
| **ULC146-bin7** | | | 0.000 | 0.011 | 0.057 | 0.000 | | |
| **ULC146-nobin** | 0.006 | 0.002 | 0.001 | 0.009 | 0.034 | 0.009 | 0.000 | 0.001 |
| **ULC165-bin1** | 0.000 | 0.000 | 0.001 | 0.007 | 0.030 | 0.001 | 0.000 | |
| **ULC165-bin2** | 0.000 | 0.000 | 0.000 | 0.059 | 0.108 | 0.001 | | 0.000 |
| **ULC165-bin3** | | | 0.001 | 0.005 | 0.143 | 0.001 | | |
| **ULC165-bin4** | 0.034 | 0.000 | 0.000 | 0.000 | 0.001 | 0.071 | 0.002 | 0.001 |
| **ULC165-nobin** | 0.010 | 0.000 | 0.000 | 0.005 | 0.047 | 0.021 | 0.001 | 0.001 |
| **ULC179-bin1** | 0.000 | 0.000 | 0.000 | 0.058 | 0.114 | 0.001 | | |
| **ULC179-bin2** | 0.000 | | 0.000 | 0.009 | 0.033 | 0.001 | | 0.000 |
| **ULC179-bin3** | 0.000 | 0.000 | 0.000 | 0.011 | 0.039 | 0.001 | | 0.000 |
| **ULC179-bin4** | 0.000 | 0.004 | 0.004 | 0.001 | 0.004 | 0.001 | 0.000 | |
| **ULC179-bin5** | 0.000 | | 0.000 | 0.050 | 0.113 | 0.001 | | |
| **ULC179-bin6** | | | 0.001 | 0.013 | 0.032 | 0.001 | | |
| **ULC179-bin7** | | | | 0.002 | 0.009 | | | |
| **ULC179-nobin** | 0.003 | 0.001 | 0.001 | 0.005 | 0.020 | 0.005 | 0.000 | 0.001 |
| **ULC186-bin1** | 0.022 | 0.000 | 0.000 | 0.001 | 0.005 | 0.042 | 0.001 | 0.001 |
| **ULC186-nobin** | 0.003 | 0.006 | 0.002 | 0.002 | 0.048 | 0.007 | 0.001 | 0.002 |
| **ULC187-bin1** | 0.031 | 0.001 | 0.001 | 0.000 | 0.001 | 0.031 | 0.000 | 0.001 |
| **ULC187-nobin** | 0.000 | 0.000 | 0.001 | 0.015 | 0.047 | 0.001 | 0.000 | 0.002 |

| | **ULC027-bin1** | **ULC027-bin2** | **ULC027-bin3** | **ULC027-bin4** | **ULC027-nobin** | **ULC041-bin1** | **ULC041-bin2** | **ULC041-nobin** |
|---|---|---|---|---|---|---|---|---|
| **ULC335-bin1** | 0.024 | 0.000 | 0.000 | | 0.002 | 0.023 | 0.001 | 0.000 |
| **ULC335-bin2** | 0.000 | 0.000 | | | 0.001 | 0.000 | | |
| **ULC335-bin3** | 0.001 | 0.001 | 0.000 | 0.000 | 0.001 | 0.001 | 0.000 | |
| **ULC335-bin4** | 0.001 | 0.050 | 0.026 | 0.006 | 0.041 | 0.003 | 0.000 | |
| **ULC335-nobin** | 0.001 | 0.016 | 0.018 | 0.006 | 0.032 | 0.003 | 0.000 | 0.001 |
| **ULC007-bin1** | 0.036 | 0.001 | 0.001 | 0.000 | 0.003 | 0.039 | 0.002 | 0.001 |
| **ULC007-bin2** | 0.009 | | | | 0.003 | 0.005 | 0.008 | |
| **ULC007-nobin** | 0.004 | 0.004 | 0.001 | | 0.028 | 0.021 | 0.001 | 0.034 |
| **ULC027-bin1** | | 0.001 | 0.001 | 0.000 | 0.009 | 0.054 | 0.003 | 0.001 |
| **ULC027-bin2** | 0.001 | | 0.016 | 0.004 | 0.027 | 0.002 | 0.000 | 0.000 |
| **ULC027-bin3** | 0.001 | 0.021 | | 0.002 | 0.076 | 0.003 | 0.000 | 0.000 |
| **ULC027-bin4** | 0.000 | 0.017 | 0.004 | | 0.062 | 0.002 | | |
| **ULC027-nobin** | 0.006 | 0.014 | 0.029 | 0.008 | | 0.004 | 0.001 | 0.001 |
| **ULC041-bin1** | 0.055 | 0.002 | 0.002 | 0.000 | 0.009 | | 0.003 | 0.003 |
| **ULC041-bin2** | 0.026 | 0.002 | 0.000 | | 0.016 | 0.017 | | 0.007 |
| **ULC041-nobin** | 0.017 | 0.005 | 0.002 | | 0.012 | 0.074 | 0.037 | |
| **ULC065-bin1** | 0.017 | 0.003 | 0.003 | 0.001 | 0.007 | 0.032 | 0.001 | 0.000 |
| **ULC065-bin2** | 0.001 | 0.012 | 0.011 | 0.003 | 0.019 | 0.005 | 0.000 | 0.000 |
| **ULC065-nobin** | 0.001 | 0.017 | 0.068 | 0.017 | 0.100 | 0.004 | 0.001 | 0.001 |
| **ULC066-bin1** | 0.018 | 0.001 | 0.000 | | 0.002 | 0.014 | 0.001 | 0.001 |
| **ULC066-bin2** | 0.001 | 0.000 | 0.000 | | 0.001 | 0.000 | | |
| **ULC066-bin3** | 0.002 | 0.002 | 0.005 | | 0.116 | 0.004 | | |
| **ULC066-nobin** | 0.001 | 0.115 | 0.045 | 0.011 | 0.075 | 0.003 | 0.000 | 0.001 |
| **ULC068-bin1** | 0.020 | 0.001 | 0.000 | 0.000 | 0.004 | 0.018 | 0.004 | 0.001 |
| **ULC068-bin2** | 0.005 | | | | 0.005 | 0.001 | | |
| **ULC068-nobin** | 0.002 | 0.040 | 0.016 | 0.004 | 0.085 | 0.004 | 0.000 | 0.001 |
| **ULC073-bin1** | 0.054 | 0.002 | 0.002 | 0.000 | 0.008 | 0.451 | 0.022 | 0.012 |
| **ULC073-bin2** | 0.001 | 0.003 | 0.004 | 0.000 | 0.110 | 0.002 | | 0.000 |
| **ULC073-bin3** | 0.001 | 0.023 | 0.303 | 0.083 | 0.137 | 0.004 | 0.000 | 0.000 |
| **ULC073-bin4** | 0.001 | 0.020 | 0.044 | 0.025 | 0.254 | 0.003 | | 0.000 |
| **ULC073-bin5** | 0.026 | 0.005 | 0.001 | | 0.025 | 0.108 | 0.201 | 0.048 |
| **ULC073-bin6** | 0.001 | 0.005 | 0.287 | 0.006 | 0.062 | 0.001 | | |
| **ULC073-nobin** | 0.002 | 0.016 | 0.062 | 0.027 | 0.220 | 0.021 | 0.002 | 0.006 |
| **ULC077-bin1** | 0.032 | 0.001 | 0.000 | 0.000 | 0.003 | 0.036 | 0.002 | 0.001 |
| **ULC077-nobin** | 0.003 | 0.010 | 0.020 | 0.004 | 0.070 | 0.004 | 0.000 | 0.001 |
| **ULC082-bin1** | 0.018 | 0.004 | 0.003 | 0.001 | 0.007 | 0.037 | 0.001 | 0.001 |
| **ULC082-bin2** | 0.002 | 0.020 | 0.018 | 0.004 | 0.035 | 0.003 | 0.000 | |
| **ULC082-bin3** | 0.002 | 0.004 | 0.006 | 0.001 | 0.013 | 0.004 | | 0.000 |
| **ULC082-bin4** | 0.001 | 0.040 | 0.028 | 0.006 | 0.041 | 0.005 | 0.001 | |
| **ULC082-bin5** | | 0.001 | 0.001 | | 0.002 | 0.001 | | |
| **ULC082-nobin** | 0.001 | 0.015 | 0.010 | 0.003 | 0.026 | 0.003 | 0.000 | 0.000 |
| **ULC084-bin1** | 0.002 | 0.004 | 0.003 | 0.001 | 0.108 | 0.003 | | 0.000 |
| **ULC084-bin2** | 0.002 | 0.036 | 0.026 | 0.007 | 0.036 | 0.003 | 0.001 | |
| **ULC084-bin3** | 0.017 | 0.004 | 0.004 | 0.001 | 0.008 | 0.035 | 0.001 | 0.001 |
| **ULC084-nobin** | 0.001 | 0.016 | 0.137 | 0.039 | 0.074 | 0.003 | 0.000 | 0.001 |
| **ULC129-bin1** | 0.128 | 0.001 | 0.001 | 0.000 | 0.014 | 0.051 | 0.002 | 0.001 |
| **ULC129-nobin** | 0.006 | 0.099 | 0.020 | 0.004 | 0.068 | 0.004 | 0.000 | 0.001 |
| **ULC146-bin1** | 0.001 | 0.008 | 0.006 | 0.002 | 0.032 | 0.003 | 0.000 | 0.000 |
| **ULC146-bin2** | 0.000 | 0.000 | 0.000 | | 0.000 | 0.000 | | |
| **ULC146-bin3** | 0.001 | 0.022 | 0.353 | 0.093 | 0.118 | 0.004 | 0.000 | 0.000 |
| **ULC146-bin4** | 0.001 | 0.003 | 0.004 | 0.000 | 0.110 | 0.002 | 0.000 | |
| **ULC146-bin5** | 0.002 | 0.043 | 0.023 | 0.004 | 0.040 | 0.004 | 0.000 | 0.000 |
| **ULC146-bin6** | 0.001 | 0.017 | 0.054 | 0.014 | 0.330 | 0.001 | | |
| **ULC146-bin7** | | 0.010 | 0.187 | 0.219 | 0.130 | | 0.001 | 0.000 |
| **ULC146-nobin** | 0.004 | 0.014 | 0.024 | 0.008 | 0.143 | 0.006 | 0.000 | 0.001 |
| **ULC165-bin1** | 0.002 | 0.011 | 0.008 | 0.003 | 0.017 | 0.004 | 0.001 | 0.000 |
| **ULC165-bin2** | 0.001 | 0.035 | 0.018 | 0.006 | 0.033 | 0.003 | 0.001 | |
| **ULC165-bin3** | 0.001 | 0.006 | 0.009 | 0.003 | 0.036 | 0.004 | 0.000 | 0.000 |
| **ULC165-bin4** | 0.034 | 0.000 | 0.000 | | 0.003 | 0.043 | 0.002 | |
| **ULC165-nobin** | 0.012 | 0.010 | 0.025 | 0.007 | 0.045 | 0.015 | 0.001 | 0.001 |
| **ULC179-bin1** | 0.001 | 0.036 | 0.020 | 0.006 | 0.036 | 0.004 | 0.000 | 0.000 |
| **ULC179-bin2** | 0.001 | 0.021 | 0.012 | 0.004 | 0.054 | 0.004 | 0.000 | 0.000 |
| **ULC179-bin3** | 0.001 | 0.026 | 0.045 | 0.018 | 0.060 | 0.003 | 0.000 | 0.000 |
| **ULC179-bin4** | 0.001 | 0.001 | 0.001 | 0.000 | 0.002 | 0.001 | 0.000 | |
| **ULC179-bin5** | 0.001 | 0.039 | 0.021 | 0.005 | 0.040 | 0.003 | | |
| **ULC179-bin6** | 0.001 | 0.019 | 0.016 | 0.006 | 0.040 | 0.004 | | 0.000 |
| **ULC179-bin7** | | 0.001 | 0.003 | | 0.014 | | | |
| **ULC179-nobin** | 0.003 | 0.010 | 0.008 | 0.001 | 0.022 | 0.003 | 0.000 | 0.001 |
| **ULC186-bin1** | 0.048 | 0.002 | 0.001 | 0.000 | 0.006 | 0.234 | 0.007 | 0.006 |
| **ULC186-nobin** | 0.009 | 0.005 | 0.004 | 0.001 | 0.018 | 0.040 | 0.003 | 0.005 |
| **ULC187-bin1** | 0.022 | 0.000 | 0.000 | | 0.002 | 0.019 | 0.001 | 0.000 |
| **ULC187-nobin** | 0.002 | 0.021 | 0.129 | 0.032 | 0.138 | 0.003 | 0.000 | 0.001 |

| | ULC065-bin1 | ULC065-bin2 | ULC065-nobin | ULC066-bin1 | ULC066-bin2 | ULC066-bin3 | ULC066-nobin | ULC068-bin1 |
|---|---|---|---|---|---|---|---|---|
| **ULC335-bin1** | 0.005 | 0.000 | 0.001 | 0.030 | 0.000 | 0.000 | 0.001 | 0.028 |
| **ULC335-bin2** | | | 0.000 | 0.001 | 0.009 | | 0.002 | 0.001 |
| **ULC335-bin3** | 0.000 | 0.001 | 0.001 | 0.001 | 0.031 | 0.000 | 0.007 | 0.001 |
| **ULC335-bin4** | 0.006 | 0.015 | 0.053 | 0.001 | 0.001 | 0.001 | 0.048 | 0.001 |
| **ULC335-nobin** | 0.004 | 0.011 | 0.046 | 0.001 | 0.004 | 0.002 | 0.033 | 0.001 |
| **ULC007-bin1** | 0.010 | 0.000 | 0.001 | 0.031 | 0.000 | 0.000 | 0.001 | 0.031 |
| **ULC007-bin2** | 0.002 | 0.001 | 0.001 | 0.015 | | | | 0.022 |
| **ULC007-nobin** | 0.007 | | 0.016 | 0.034 | 0.003 | | 0.019 | 0.034 |
| **ULC027-bin1** | 0.011 | 0.001 | 0.002 | 0.020 | 0.000 | 0.000 | 0.002 | 0.019 |
| **ULC027-bin2** | 0.002 | 0.009 | 0.024 | 0.001 | 0.000 | 0.000 | 0.256 | 0.000 |
| **ULC027-bin3** | 0.003 | 0.011 | 0.157 | 0.001 | 0.000 | 0.001 | 0.127 | 0.001 |
| **ULC027-bin4** | 0.003 | 0.010 | 0.106 | | | | 0.092 | 0.001 |
| **ULC027-nobin** | 0.003 | 0.008 | 0.079 | 0.001 | 0.000 | 0.008 | 0.077 | 0.002 |
| **ULC041-bin1** | 0.021 | 0.003 | 0.005 | 0.016 | 0.000 | 0.001 | 0.004 | 0.018 |
| **ULC041-bin2** | 0.005 | 0.001 | 0.006 | 0.009 | | | 0.003 | 0.025 |
| **ULC041-nobin** | 0.003 | 0.001 | 0.008 | 0.013 | | | 0.010 | 0.009 |
| **ULC065-bin1** | | 0.006 | 0.007 | 0.009 | 0.000 | 0.000 | 0.006 | 0.010 |
| **ULC065-bin2** | 0.006 | | 0.019 | 0.000 | 0.000 | 0.003 | 0.022 | 0.001 |
| **ULC065-nobin** | 0.004 | 0.009 | | 0.001 | 0.000 | 0.001 | 0.087 | 0.001 |
| **ULC066-bin1** | 0.005 | 0.000 | 0.001 | | 0.000 | 0.000 | 0.002 | 0.215 |
| **ULC066-bin2** | 0.000 | 0.000 | 0.000 | 0.001 | | | 0.002 | 0.001 |
| **ULC066-bin3** | 0.001 | 0.013 | 0.009 | 0.001 | | | 0.011 | 0.001 |
| **ULC066-nobin** | 0.003 | 0.009 | 0.070 | 0.001 | 0.001 | 0.001 | | 0.001 |
| **ULC068-bin1** | 0.007 | 0.001 | 0.001 | 0.251 | 0.001 | 0.000 | 0.003 | |
| **ULC068-bin2** | | | | 0.044 | | | | 0.045 |
| **ULC068-nobin** | 0.004 | 0.110 | 0.105 | 0.009 | 0.000 | 0.001 | 0.073 | 0.005 |
| **ULC073-bin1** | 0.022 | 0.003 | 0.004 | 0.017 | 0.000 | 0.000 | 0.004 | 0.017 |
| **ULC073-bin2** | 0.002 | 0.009 | 0.009 | 0.001 | 0.001 | 0.073 | 0.303 | 0.001 |
| **ULC073-bin3** | 0.004 | 0.010 | 0.123 | 0.000 | 0.000 | 0.001 | 0.102 | 0.000 |
| **ULC073-bin4** | 0.003 | 0.011 | 0.036 | | 0.000 | 0.001 | 0.034 | 0.000 |
| **ULC073-bin5** | 0.007 | | 0.001 | 0.017 | | 0.006 | 0.015 | 0.022 |
| **ULC073-bin6** | 0.001 | | 0.198 | 0.001 | 0.001 | | 0.181 | |
| **ULC073-nobin** | 0.003 | 0.009 | 0.079 | 0.001 | 0.000 | 0.001 | 0.057 | 0.001 |
| **ULC077-bin1** | 0.009 | 0.001 | 0.001 | 0.033 | 0.000 | 0.000 | 0.002 | 0.030 |
| **ULC077-nobin** | 0.004 | 0.129 | 0.046 | 0.002 | 0.000 | 0.029 | 0.147 | 0.002 |
| **ULC082-bin1** | 0.232 | 0.005 | 0.007 | 0.011 | 0.000 | 0.000 | 0.007 | 0.007 |
| **ULC082-bin2** | 0.004 | 0.010 | 0.132 | 0.000 | 0.000 | 0.001 | 0.051 | 0.001 |
| **ULC082-bin3** | 0.003 | 0.016 | 0.013 | 0.001 | 0.001 | 0.002 | 0.016 | 0.001 |
| **ULC082-bin4** | 0.006 | 0.021 | 0.048 | 0.000 | | 0.001 | 0.051 | 0.000 |
| **ULC082-bin5** | | 0.000 | 0.001 | | | | 0.003 | |
| **ULC082-nobin** | 0.004 | 0.013 | 0.042 | 0.000 | 0.000 | 0.002 | 0.026 | 0.001 |
| **ULC084-bin1** | 0.002 | 0.011 | 0.009 | 0.001 | 0.001 | 0.085 | 0.329 | 0.001 |
| **ULC084-bin2** | 0.003 | 0.013 | 0.071 | 0.001 | 0.000 | 0.000 | 0.037 | 0.001 |
| **ULC084-bin3** | 0.225 | 0.006 | 0.009 | 0.008 | 0.000 | 0.000 | 0.007 | 0.009 |
| **ULC084-nobin** | 0.008 | 0.007 | 0.105 | 0.001 | 0.000 | 0.007 | 0.102 | 0.001 |
| **ULC129-bin1** | 0.012 | 0.001 | 0.001 | 0.019 | 0.000 | 0.000 | 0.002 | 0.017 |
| **ULC129-nobin** | 0.002 | 0.007 | 0.042 | 0.001 | 0.001 | 0.017 | 0.142 | 0.001 |
| **ULC146-bin1** | 0.005 | 0.018 | 0.071 | 0.000 | 0.000 | 0.005 | 0.028 | 0.000 |
| **ULC146-bin2** | 0.000 | 0.000 | | 0.001 | 0.007 | | 0.001 | 0.001 |
| **ULC146-bin3** | 0.003 | 0.012 | 0.144 | 0.000 | 0.000 | 0.001 | 0.120 | 0.000 |
| **ULC146-bin4** | 0.002 | 0.009 | 0.009 | 0.001 | 0.001 | 0.083 | 0.321 | 0.001 |
| **ULC146-bin5** | 0.004 | 0.016 | 0.047 | 0.000 | 0.000 | 0.000 | 0.045 | 0.001 |
| **ULC146-bin6** | 0.003 | 0.007 | 0.031 | 0.000 | 0.000 | 0.000 | 0.034 | 0.001 |
| **ULC146-bin7** | 0.002 | 0.007 | 0.125 | | | | 0.103 | 0.000 |
| **ULC146-nobin** | 0.003 | 0.006 | 0.052 | 0.004 | 0.001 | 0.001 | 0.043 | 0.004 |
| **ULC165-bin1** | 0.006 | 0.415 | 0.076 | 0.000 | 0.000 | 0.001 | 0.020 | 0.001 |
| **ULC165-bin2** | 0.004 | 0.013 | 0.072 | 0.000 | 0.000 | 0.001 | 0.038 | 0.000 |
| **ULC165-bin3** | 0.004 | 0.019 | 0.086 | 0.000 | 0.000 | 0.004 | 0.033 | 0.000 |
| **ULC165-bin4** | 0.008 | 0.000 | 0.001 | 0.032 | 0.000 | 0.000 | 0.002 | 0.034 |
| **ULC165-nobin** | 0.005 | 0.010 | 0.040 | 0.009 | 0.000 | 0.001 | 0.031 | 0.009 |
| **ULC179-bin1** | 0.004 | 0.013 | 0.040 | 0.000 | 0.000 | 0.001 | 0.039 | 0.001 |
| **ULC179-bin2** | 0.003 | 0.009 | 0.036 | 0.001 | 0.000 | 0.000 | 0.038 | 0.001 |
| **ULC179-bin3** | 0.004 | 0.015 | 0.039 | 0.001 | 0.000 | 0.000 | 0.038 | 0.001 |
| **ULC179-bin4** | 0.001 | 0.002 | 0.002 | 0.000 | 0.004 | 0.000 | 0.002 | 0.001 |
| **ULC179-bin5** | 0.004 | 0.015 | 0.040 | 0.001 | | 0.001 | 0.037 | 0.000 |
| **ULC179-bin6** | 0.003 | 0.009 | 0.043 | 0.000 | | 0.000 | 0.033 | 0.001 |
| **ULC179-bin7** | | | 0.009 | | | | 0.009 | |
| **ULC179-nobin** | 0.003 | 0.005 | 0.021 | 0.002 | 0.001 | 0.000 | 0.018 | 0.003 |
| **ULC186-bin1** | 0.022 | 0.003 | 0.004 | 0.019 | 0.000 | 0.000 | 0.004 | 0.018 |
| **ULC186-nobin** | 0.004 | 0.007 | 0.020 | 0.003 | 0.004 | 0.001 | 0.015 | 0.003 |
| **ULC187-bin1** | 0.008 | 0.000 | 0.001 | 0.201 | 0.000 | 0.000 | 0.002 | 0.187 |
| **ULC187-nobin** | 0.003 | 0.011 | 0.116 | 0.005 | 0.000 | 0.005 | 0.101 | 0.003 |

| | **ULC068-bin2** | **ULC068-nobin** | **ULC073-bin1** | **ULC073-bin2** | **ULC073-bin3** | **ULC073-bin4** | **ULC073-bin5** | **ULC073-bin6** |
|---|---|---|---|---|---|---|---|---|
| **ULC335-bin1** | 0.000 | 0.001 | 0.023 | 0.001 | 0.000 | 0.000 | 0.001 | |
| **ULC335-bin2** | | 0.007 | 0.000 | 0.001 | 0.000 | 0.000 | | |
| **ULC335-bin3** | 0.000 | 0.001 | 0.001 | 0.001 | 0.000 | | | |
| **ULC335-bin4** | | 0.027 | 0.004 | 0.004 | 0.031 | 0.006 | | 0.000 |
| **ULC335-nobin** | 0.000 | 0.015 | 0.004 | 0.011 | 0.021 | 0.003 | 0.000 | 0.001 |
| **ULC007-bin1** | 0.000 | 0.002 | 0.037 | 0.001 | 0.001 | 0.000 | 0.001 | |
| **ULC007-bin2** | | | 0.001 | | | 0.001 | 0.006 | |
| **ULC007-nobin** | 0.007 | 0.011 | 0.013 | 0.002 | 0.001 | | 0.001 | 0.004 |
| **ULC027-bin1** | 0.000 | 0.001 | 0.053 | 0.001 | 0.001 | 0.000 | 0.002 | 0.000 |
| **ULC027-bin2** | | 0.028 | 0.003 | 0.003 | 0.019 | 0.003 | 0.000 | 0.000 |
| **ULC027-bin3** | | 0.014 | 0.003 | 0.004 | 0.352 | 0.009 | 0.000 | 0.031 |
| **ULC027-bin4** | | 0.009 | 0.002 | 0.002 | 0.279 | 0.015 | | 0.002 |
| **ULC027-nobin** | 0.000 | 0.029 | 0.004 | 0.044 | 0.057 | 0.019 | 0.001 | 0.002 |
| **ULC041-bin1** | 0.000 | 0.002 | 0.440 | 0.001 | 0.003 | 0.000 | 0.007 | 0.000 |
| **ULC041-bin2** | | 0.002 | 0.182 | | 0.000 | | 0.111 | |
| **ULC041-nobin** | | 0.006 | 0.179 | 0.001 | 0.001 | 0.001 | 0.049 | |
| **ULC065-bin1** | | 0.004 | 0.033 | 0.002 | 0.004 | 0.001 | 0.001 | 0.000 |
| **ULC065-bin2** | | 0.109 | 0.004 | 0.011 | 0.011 | 0.002 | | |
| **ULC065-nobin** | | 0.045 | 0.003 | 0.005 | 0.063 | 0.004 | 0.000 | 0.009 |
| **ULC066-bin1** | 0.002 | 0.006 | 0.015 | 0.001 | 0.000 | | 0.001 | 0.000 |
| **ULC066-bin2** | | 0.000 | 0.000 | 0.001 | 0.000 | 0.000 | | 0.000 |
| **ULC066-bin3** | | 0.005 | 0.004 | 0.461 | 0.006 | 0.001 | 0.003 | |
| **ULC066-nobin** | | 0.026 | 0.003 | 0.113 | 0.043 | 0.003 | 0.000 | 0.006 |
| **ULC068-bin1** | 0.002 | 0.003 | 0.017 | 0.001 | 0.000 | 0.000 | 0.001 | |
| **ULC068-bin2** | | 0.002 | | 0.001 | | | | |
| **ULC068-nobin** | 0.000 | | 0.004 | 0.007 | 0.017 | 0.003 | 0.000 | 0.001 |
| **ULC073-bin1** | | 0.003 | | 0.002 | 0.002 | 0.000 | 0.002 | 0.000 |
| **ULC073-bin2** | 0.000 | 0.005 | 0.003 | | 0.003 | 0.000 | 0.000 | |
| **ULC073-bin3** | | 0.013 | 0.002 | 0.004 | | 0.011 | 0.000 | 0.002 |
| **ULC073-bin4** | | 0.014 | 0.002 | 0.002 | 0.063 | | | |
| **ULC073-bin5** | | 0.003 | 0.015 | 0.001 | 0.001 | | | |
| **ULC073-bin6** | | 0.012 | 0.001 | | 0.021 | | | |
| **ULC073-nobin** | 0.000 | 0.026 | 0.004 | 0.005 | 0.040 | 0.007 | 0.001 | 0.001 |
| **ULC077-bin1** | 0.000 | 0.001 | 0.037 | 0.001 | 0.000 | 0.000 | 0.001 | |
| **ULC077-nobin** | 0.000 | 0.048 | 0.004 | 0.182 | 0.018 | 0.002 | 0.001 | 0.003 |
| **ULC082-bin1** | | 0.004 | 0.035 | 0.002 | 0.004 | 0.001 | 0.001 | |
| **ULC082-bin2** | 0.000 | 0.021 | 0.004 | 0.004 | 0.017 | 0.003 | | 0.000 |
| **ULC082-bin3** | | 0.008 | 0.003 | 0.012 | 0.006 | 0.001 | | 0.000 |
| **ULC082-bin4** | | 0.026 | 0.003 | 0.008 | 0.032 | 0.008 | | 0.001 |
| **ULC082-bin5** | | 0.002 | | 0.003 | 0.001 | | | |
| **ULC082-nobin** | 0.000 | 0.015 | 0.003 | 0.008 | 0.011 | 0.002 | 0.000 | 0.001 |
| **ULC084-bin1** | 0.000 | 0.005 | 0.003 | 0.502 | 0.004 | 0.000 | 0.000 | 0.000 |
| **ULC084-bin2** | | 0.032 | 0.003 | 0.003 | 0.024 | 0.003 | | 0.000 |
| **ULC084-bin3** | 0.000 | 0.005 | 0.039 | 0.002 | 0.003 | 0.000 | 0.002 | |
| **ULC084-nobin** | 0.000 | 0.017 | 0.003 | 0.049 | 0.133 | 0.005 | 0.002 | 0.018 |
| **ULC129-bin1** | 0.000 | 0.001 | 0.048 | 0.001 | 0.001 | 0.000 | 0.001 | 0.000 |
| **ULC129-nobin** | 0.000 | 0.020 | 0.004 | 0.097 | 0.020 | 0.003 | 0.001 | 0.003 |
| **ULC146-bin1** | | 0.015 | 0.004 | 0.023 | 0.008 | 0.001 | 0.000 | 0.000 |
| **ULC146-bin2** | | 0.006 | 0.000 | 0.000 | 0.000 | | | 0.000 |
| **ULC146-bin3** | 0.000 | 0.016 | 0.003 | 0.004 | 0.344 | 0.009 | | 0.021 |
| **ULC146-bin4** | 0.000 | 0.005 | 0.002 | 0.456 | 0.003 | 0.000 | 0.000 | |
| **ULC146-bin5** | | 0.023 | 0.004 | 0.003 | 0.024 | 0.003 | | 0.001 |
| **ULC146-bin6** | 0.001 | 0.010 | 0.002 | 0.000 | 0.077 | 0.020 | | 0.001 |
| **ULC146-bin7** | | 0.008 | 0.002 | 0.003 | 0.308 | 0.009 | 0.000 | 0.000 |
| **ULC146-nobin** | 0.000 | 0.023 | 0.005 | 0.005 | 0.034 | 0.014 | 0.001 | 0.003 |
| **ULC165-bin1** | | 0.105 | 0.004 | 0.009 | 0.011 | 0.002 | | 0.000 |
| **ULC165-bin2** | | 0.030 | 0.004 | 0.004 | 0.023 | 0.003 | 0.000 | 0.001 |
| **ULC165-bin3** | | 0.016 | 0.003 | 0.027 | 0.012 | 0.001 | 0.000 | 0.000 |
| **ULC165-bin4** | 0.001 | 0.000 | 0.037 | 0.002 | | | 0.002 | |
| **ULC165-nobin** | 0.000 | 0.013 | 0.014 | 0.005 | 0.029 | 0.005 | 0.001 | 0.002 |
| **ULC179-bin1** | | 0.020 | 0.003 | 0.004 | 0.025 | 0.005 | 0.000 | 0.001 |
| **ULC179-bin2** | | 0.019 | 0.003 | 0.003 | 0.016 | 0.003 | 0.000 | 0.000 |
| **ULC179-bin3** | 0.000 | 0.017 | 0.003 | 0.003 | 0.051 | 0.008 | | 0.000 |
| **ULC179-bin4** | | 0.002 | 0.001 | 0.001 | 0.001 | 0.000 | 0.000 | |
| **ULC179-bin5** | | 0.021 | 0.003 | 0.003 | 0.026 | 0.004 | | 0.001 |
| **ULC179-bin6** | | 0.022 | 0.003 | 0.004 | 0.022 | 0.005 | | 0.000 |
| **ULC179-bin7** | | 0.002 | | 0.000 | 0.006 | 0.004 | | 0.002 |
| **ULC179-nobin** | 0.000 | 0.010 | 0.003 | 0.002 | 0.008 | 0.001 | 0.000 | 0.001 |
| **ULC186-bin1** | 0.000 | 0.002 | 0.214 | 0.001 | 0.002 | 0.000 | 0.004 | 0.000 |
| **ULC186-nobin** | 0.000 | 0.009 | 0.038 | 0.007 | 0.003 | 0.000 | 0.002 | 0.001 |
| **ULC187-bin1** | 0.006 | 0.005 | 0.019 | 0.001 | 0.000 | 0.000 | 0.000 | |
| **ULC187-nobin** | 0.015 | 0.036 | 0.003 | 0.031 | 0.126 | 0.012 | 0.000 | 0.019 |

| | ULC073-nobin | ULC077-bin1 | ULC077-nobin | ULC082-bin1 | ULC082-bin2 | ULC082-bin3 | ULC082-bin4 | ULC082-bin5 |
|---|---|---|---|---|---|---|---|---|
| **ULC335-bin1** | 0.001 | 0.042 | 0.002 | 0.006 | 0.000 | 0.000 | | |
| **ULC335-bin2** | 0.000 | 0.001 | 0.000 | | 0.000 | 0.000 | 0.000 | |
| **ULC335-bin3** | 0.000 | 0.000 | 0.001 | 0.001 | 0.001 | 0.001 | 0.000 | |
| **ULC335-bin4** | 0.042 | 0.001 | 0.036 | 0.006 | 0.060 | 0.007 | 0.046 | 0.000 |
| **ULC335-nobin** | 0.035 | 0.001 | 0.021 | 0.003 | 0.025 | 0.008 | 0.010 | 0.000 |
| **ULC007-bin1** | 0.003 | 0.081 | 0.002 | 0.010 | 0.000 | 0.001 | 0.000 | |
| **ULC007-bin2** | 0.001 | 0.028 | 0.027 | 0.002 | | | | |
| **ULC007-nobin** | 0.027 | 0.018 | 0.041 | 0.006 | 0.000 | | | |
| **ULC027-bin1** | 0.004 | 0.037 | 0.003 | 0.011 | 0.001 | 0.001 | 0.000 | |
| **ULC027-bin2** | 0.026 | 0.001 | 0.012 | 0.003 | 0.013 | 0.002 | 0.005 | 0.000 |
| **ULC027-bin3** | 0.141 | 0.000 | 0.036 | 0.002 | 0.014 | 0.004 | 0.004 | 0.000 |
| **ULC027-bin4** | 0.182 | 0.001 | 0.024 | 0.001 | 0.010 | 0.002 | 0.003 | |
| **ULC027-nobin** | 0.175 | 0.002 | 0.044 | 0.003 | 0.011 | 0.004 | 0.003 | 0.000 |
| **ULC041-bin1** | 0.033 | 0.042 | 0.005 | 0.022 | 0.002 | 0.002 | 0.001 | 0.000 |
| **ULC041-bin2** | 0.018 | 0.017 | 0.004 | 0.005 | 0.002 | | 0.001 | |
| **ULC041-nobin** | 0.136 | 0.013 | 0.017 | 0.006 | | 0.001 | | |
| **ULC065-bin1** | 0.006 | 0.016 | 0.006 | 0.211 | 0.003 | 0.002 | 0.001 | |
| **ULC065-bin2** | 0.018 | 0.001 | 0.243 | 0.005 | 0.009 | 0.012 | 0.004 | 0.000 |
| **ULC065-nobin** | 0.081 | 0.001 | 0.035 | 0.003 | 0.049 | 0.005 | 0.004 | 0.000 |
| **ULC066-bin1** | 0.001 | 0.033 | 0.002 | 0.005 | 0.000 | 0.000 | 0.000 | |
| **ULC066-bin2** | 0.000 | 0.000 | 0.000 | 0.000 | 0.000 | 0.001 | | |
| **ULC066-bin3** | 0.011 | 0.000 | 0.268 | 0.001 | 0.003 | 0.007 | 0.001 | |
| **ULC066-nobin** | 0.047 | 0.001 | 0.086 | 0.003 | 0.017 | 0.005 | 0.003 | 0.000 |
| **ULC068-bin1** | 0.001 | 0.036 | 0.002 | 0.004 | 0.000 | 0.001 | 0.000 | |
| **ULC068-bin2** | 0.002 | 0.003 | 0.002 | | 0.002 | | | |
| **ULC068-nobin** | 0.065 | 0.002 | 0.093 | 0.004 | 0.019 | 0.007 | 0.005 | 0.000 |
| **ULC073-bin1** | 0.006 | 0.044 | 0.005 | 0.021 | 0.002 | 0.002 | 0.000 | |
| **ULC073-bin2** | 0.008 | 0.002 | 0.275 | 0.002 | 0.003 | 0.007 | 0.001 | 0.000 |
| **ULC073-bin3** | 0.078 | 0.000 | 0.028 | 0.003 | 0.013 | 0.003 | 0.004 | 0.000 |
| **ULC073-bin4** | 0.078 | 0.002 | 0.019 | 0.003 | 0.011 | 0.003 | 0.006 | |
| **ULC073-bin5** | 0.005 | 0.027 | 0.023 | 0.009 | | | | |
| **ULC073-bin6** | 0.024 | | 0.043 | | 0.002 | 0.001 | 0.001 | |
| **ULC073-nobin** | | 0.002 | 0.025 | 0.003 | 0.011 | 0.004 | 0.003 | 0.000 |
| **ULC077-bin1** | 0.002 | | 0.011 | 0.008 | 0.001 | 0.001 | 0.000 | |
| **ULC077-nobin** | 0.032 | 0.018 | | 0.003 | 0.008 | 0.007 | 0.004 | 0.000 |
| **ULC082-bin1** | 0.006 | 0.016 | 0.007 | | 0.003 | 0.003 | 0.001 | 0.000 |
| **ULC082-bin2** | 0.028 | 0.001 | 0.015 | 0.003 | | 0.003 | 0.004 | |
| **ULC082-bin3** | 0.011 | 0.002 | 0.017 | 0.004 | 0.004 | | 0.001 | 0.001 |
| **ULC082-bin4** | 0.043 | 0.002 | 0.041 | 0.007 | 0.022 | 0.005 | | |
| **ULC082-bin5** | 0.001 | | 0.003 | | | 0.004 | | |
| **ULC082-nobin** | 0.025 | 0.001 | 0.021 | 0.003 | 0.011 | 0.005 | 0.006 | 0.001 |
| **ULC084-bin1** | 0.017 | 0.001 | 0.279 | 0.002 | 0.003 | 0.008 | 0.001 | 0.000 |
| **ULC084-bin2** | 0.030 | 0.001 | 0.065 | 0.004 | 0.039 | 0.005 | 0.032 | 0.000 |
| **ULC084-bin3** | 0.006 | 0.014 | 0.006 | 0.383 | 0.003 | 0.003 | 0.001 | |
| **ULC084-nobin** | 0.087 | 0.001 | 0.052 | 0.014 | 0.033 | 0.004 | 0.004 | 0.000 |
| **ULC129-bin1** | 0.004 | 0.036 | 0.002 | 0.010 | 0.001 | 0.001 | 0.000 | |
| **ULC129-nobin** | 0.041 | 0.001 | 0.066 | 0.002 | 0.010 | 0.004 | 0.002 | 0.000 |
| **ULC146-bin1** | 0.062 | 0.001 | 0.028 | 0.005 | 0.006 | 0.013 | 0.002 | 0.000 |
| **ULC146-bin2** | | 0.000 | 0.000 | | 0.000 | 0.000 | 0.000 | |
| **ULC146-bin3** | 0.168 | 0.001 | 0.033 | 0.003 | 0.013 | 0.003 | 0.005 | 0.000 |
| **ULC146-bin4** | 0.020 | 0.001 | 0.278 | 0.001 | 0.002 | 0.007 | 0.001 | 0.001 |
| **ULC146-bin5** | 0.035 | 0.000 | 0.034 | 0.005 | 0.053 | 0.005 | 0.038 | 0.000 |
| **ULC146-bin6** | 0.305 | 0.001 | 0.014 | 0.001 | 0.010 | 0.004 | 0.006 | 0.000 |
| **ULC146-bin7** | 0.215 | 0.001 | 0.026 | 0.004 | 0.007 | 0.001 | 0.005 | |
| **ULC146-nobin** | 0.126 | 0.007 | 0.020 | 0.003 | 0.010 | 0.003 | 0.003 | 0.000 |
| **ULC165-bin1** | 0.017 | 0.001 | 0.235 | 0.005 | 0.006 | 0.013 | 0.003 | 0.000 |
| **ULC165-bin2** | 0.030 | 0.001 | 0.065 | 0.004 | 0.040 | 0.004 | 0.024 | |
| **ULC165-bin3** | 0.068 | 0.000 | 0.031 | 0.005 | 0.007 | 0.022 | 0.001 | 0.001 |
| **ULC165-bin4** | 0.003 | 0.068 | 0.003 | 0.008 | | 0.001 | 0.000 | |
| **ULC165-nobin** | 0.051 | 0.020 | 0.016 | 0.005 | 0.007 | 0.004 | 0.002 | 0.000 |
| **ULC179-bin1** | 0.033 | 0.001 | 0.027 | 0.004 | 0.029 | 0.005 | 0.029 | 0.000 |
| **ULC179-bin2** | 0.028 | 0.000 | 0.012 | 0.003 | 0.014 | 0.004 | 0.004 | 0.000 |
| **ULC179-bin3** | 0.060 | 0.001 | 0.014 | 0.004 | 0.015 | 0.005 | 0.004 | |
| **ULC179-bin4** | 0.002 | 0.001 | 0.001 | 0.001 | 0.001 | 0.001 | 0.000 | |
| **ULC179-bin5** | 0.030 | 0.001 | 0.028 | 0.004 | 0.040 | 0.003 | 0.031 | |
| **ULC179-bin6** | 0.031 | 0.001 | 0.012 | 0.003 | 0.024 | 0.008 | 0.002 | |
| **ULC179-bin7** | 0.014 | 0.001 | 0.004 | 0.002 | 0.003 | | 0.000 | |
| **ULC179-nobin** | 0.015 | 0.004 | 0.009 | 0.002 | 0.007 | 0.002 | 0.002 | 0.000 |
| **ULC186-bin1** | 0.017 | 0.042 | 0.005 | 0.022 | 0.002 | 0.002 | 0.000 | 0.000 |
| **ULC186-nobin** | 0.024 | 0.007 | 0.015 | 0.004 | 0.004 | 0.005 | 0.001 | |
| **ULC187-bin1** | 0.002 | 0.033 | 0.002 | 0.007 | 0.001 | 0.001 | 0.000 | |
| **ULC187-nobin** | 0.122 | 0.001 | 0.051 | 0.003 | 0.017 | 0.003 | 0.007 | |

| | ULC082-nobin | ULC084-bin1 | ULC084-bin2 | ULC084-bin3 | ULC084-nobin | ULC129-bin1 | ULC129-nobin | ULC146-bin1 |
|---|---|---|---|---|---|---|---|---|
| **ULC335-bin1** | 0.001 | 0.001 | 0.000 | 0.006 | 0.000 | 0.023 | 0.001 | 0.000 |
| **ULC335-bin2** | 0.000 | 0.000 | 0.000 | 0.000 | 0.000 | 0.000 | 0.004 | 0.000 |
| **ULC335-bin3** | 0.002 | 0.002 | 0.001 | 0.000 | 0.001 | 0.001 | 0.004 | 0.001 |
| **ULC335-bin4** | 0.237 | 0.007 | 0.165 | 0.006 | 0.023 | 0.001 | 0.027 | 0.019 |
| **ULC335-nobin** | 0.114 | 0.010 | 0.037 | 0.003 | 0.023 | 0.002 | 0.023 | 0.085 |
| **ULC007-bin1** | 0.003 | 0.001 | 0.001 | 0.008 | 0.001 | 0.032 | 0.002 | 0.001 |
| **ULC007-bin2** | 0.001 | 0.001 | | 0.005 | | 0.009 | 0.002 | |
| **ULC007-nobin** | 0.011 | | 0.002 | 0.006 | 0.013 | 0.015 | 0.017 | 0.009 |
| **ULC027-bin1** | 0.003 | 0.001 | 0.001 | 0.010 | 0.001 | 0.127 | 0.009 | 0.002 |
| **ULC027-bin2** | 0.043 | 0.003 | 0.029 | 0.002 | 0.013 | 0.001 | 0.165 | 0.010 |
| **ULC027-bin3** | 0.034 | 0.003 | 0.027 | 0.004 | 0.175 | 0.001 | 0.044 | 0.011 |
| **ULC027-bin4** | 0.028 | 0.002 | 0.021 | 0.002 | 0.140 | 0.001 | 0.028 | 0.011 |
| **ULC027-nobin** | 0.039 | 0.039 | 0.015 | 0.003 | 0.034 | 0.007 | 0.054 | 0.020 |
| **ULC041-bin1** | 0.007 | 0.002 | 0.002 | 0.021 | 0.002 | 0.052 | 0.006 | 0.004 |
| **ULC041-bin2** | 0.006 | | 0.003 | 0.005 | 0.001 | 0.020 | 0.003 | 0.000 |
| **ULC041-nobin** | 0.007 | 0.001 | | 0.006 | 0.010 | 0.009 | 0.009 | 0.007 |
| **ULC065-bin1** | 0.016 | 0.002 | 0.003 | 0.205 | 0.009 | 0.019 | 0.003 | 0.008 |
| **ULC065-bin2** | 0.048 | 0.012 | 0.014 | 0.007 | 0.009 | 0.002 | 0.013 | 0.030 |
| **ULC065-nobin** | 0.081 | 0.005 | 0.035 | 0.004 | 0.058 | 0.001 | 0.042 | 0.055 |
| **ULC066-bin1** | 0.001 | 0.001 | 0.000 | 0.004 | 0.000 | 0.016 | 0.001 | 0.000 |
| **ULC066-bin2** | 0.001 | 0.001 | 0.000 | 0.000 | 0.000 | 0.000 | 0.003 | 0.000 |
| **ULC066-bin3** | 0.035 | 0.478 | 0.002 | 0.001 | 0.052 | 0.002 | 0.189 | 0.046 |
| **ULC066-nobin** | 0.041 | 0.110 | 0.016 | 0.003 | 0.046 | 0.001 | 0.103 | 0.019 |
| **ULC068-bin1** | 0.002 | 0.000 | 0.001 | 0.006 | 0.001 | 0.017 | 0.001 | 0.000 |
| **ULC068-bin2** | 0.003 | 0.002 | | 0.002 | 0.001 | 0.003 | 0.002 | |
| **ULC068-nobin** | 0.059 | 0.007 | 0.037 | 0.005 | 0.022 | 0.002 | 0.046 | 0.029 |
| **ULC073-bin1** | 0.007 | 0.002 | 0.002 | 0.024 | 0.002 | 0.049 | 0.006 | 0.004 |
| **ULC073-bin2** | 0.029 | 0.452 | 0.003 | 0.002 | 0.058 | 0.001 | 0.179 | 0.033 |
| **ULC073-bin3** | 0.036 | 0.004 | 0.022 | 0.003 | 0.145 | 0.001 | 0.039 | 0.012 |
| **ULC073-bin4** | 0.026 | 0.001 | 0.016 | 0.002 | 0.028 | 0.001 | 0.028 | 0.010 |
| **ULC073-bin5** | 0.008 | 0.001 | | 0.016 | 0.032 | 0.011 | 0.018 | 0.002 |
| **ULC073-bin6** | 0.016 | 0.001 | 0.002 | | 0.221 | 0.003 | 0.068 | 0.005 |
| **ULC073-nobin** | 0.041 | 0.007 | 0.015 | 0.003 | 0.048 | 0.003 | 0.039 | 0.046 |
| **ULC077-bin1** | 0.002 | 0.001 | 0.001 | 0.007 | 0.001 | 0.031 | 0.002 | 0.001 |
| **ULC077-nobin** | 0.048 | 0.166 | 0.039 | 0.003 | 0.039 | 0.002 | 0.081 | 0.028 |
| **ULC082-bin1** | 0.011 | 0.002 | 0.004 | 0.387 | 0.020 | 0.017 | 0.003 | 0.008 |
| **ULC082-bin2** | 0.052 | 0.004 | 0.047 | 0.004 | 0.049 | 0.002 | 0.025 | 0.013 |
| **ULC082-bin3** | 0.031 | 0.013 | 0.007 | 0.004 | 0.008 | 0.003 | 0.010 | 0.031 |
| **ULC082-bin4** | 0.161 | 0.005 | 0.225 | 0.006 | 0.030 | 0.000 | 0.030 | 0.021 |
| **ULC082-bin5** | 0.020 | 0.003 | 0.001 | | 0.001 | 0.000 | 0.001 | 0.003 |
| **ULC082-nobin** | | 0.009 | 0.057 | 0.005 | 0.020 | 0.001 | 0.019 | 0.074 |
| **ULC084-bin1** | 0.035 | | 0.003 | 0.002 | 0.008 | 0.001 | 0.181 | 0.033 |
| **ULC084-bin2** | 0.215 | 0.003 | | 0.006 | 0.016 | 0.001 | 0.019 | 0.012 |
| **ULC084-bin3** | 0.023 | 0.002 | 0.007 | | 0.006 | 0.019 | 0.004 | 0.007 |
| **ULC084-nobin** | 0.067 | 0.007 | 0.014 | 0.004 | | 0.001 | 0.050 | 0.021 |
| **ULC129-bin1** | 0.003 | 0.001 | 0.001 | 0.012 | 0.001 | | 0.007 | 0.001 |
| **ULC129-nobin** | 0.035 | 0.088 | 0.010 | 0.002 | 0.030 | 0.006 | | 0.043 |
| **ULC146-bin1** | 0.174 | 0.020 | 0.007 | 0.004 | 0.015 | 0.001 | 0.054 | |
| **ULC146-bin2** | 0.000 | 0.000 | 0.000 | 0.000 | 0.000 | 0.000 | 0.004 | 0.000 |
| **ULC146-bin3** | 0.039 | 0.003 | 0.025 | 0.004 | 0.172 | 0.001 | 0.038 | 0.012 |
| **ULC146-bin4** | 0.030 | 0.439 | 0.003 | 0.002 | 0.048 | 0.001 | 0.180 | 0.029 |
| **ULC146-bin5** | 0.248 | 0.005 | 0.185 | 0.005 | 0.024 | 0.001 | 0.024 | 0.013 |
| **ULC146-bin6** | 0.024 | 0.001 | 0.011 | 0.002 | 0.024 | 0.001 | 0.020 | 0.008 |
| **ULC146-bin7** | 0.026 | 0.002 | 0.022 | 0.003 | 0.148 | | 0.031 | 0.008 |
| **ULC146-nobin** | 0.047 | 0.004 | 0.020 | 0.003 | 0.028 | 0.004 | 0.030 | 0.009 |
| **ULC165-bin1** | 0.040 | 0.010 | 0.013 | 0.007 | 0.009 | 0.002 | 0.012 | 0.029 |
| **ULC165-bin2** | 0.210 | 0.004 | 0.483 | 0.004 | 0.038 | 0.001 | 0.020 | 0.013 |
| **ULC165-bin3** | 0.169 | 0.025 | 0.008 | 0.004 | 0.016 | 0.002 | 0.058 | 0.477 |
| **ULC165-bin4** | 0.001 | 0.001 | 0.000 | 0.006 | 0.001 | 0.034 | 0.004 | 0.001 |
| **ULC165-nobin** | 0.058 | 0.005 | 0.013 | 0.005 | 0.020 | 0.011 | 0.022 | 0.123 |
| **ULC179-bin1** | 0.185 | 0.003 | 0.149 | 0.003 | 0.022 | 0.001 | 0.021 | 0.014 |
| **ULC179-bin2** | 0.064 | 0.003 | 0.018 | 0.003 | 0.016 | 0.001 | 0.016 | 0.015 |
| **ULC179-bin3** | 0.040 | 0.003 | 0.027 | 0.005 | 0.026 | 0.002 | 0.017 | 0.014 |
| **ULC179-bin4** | 0.002 | 0.001 | 0.002 | 0.001 | 0.001 | 0.001 | 0.005 | 0.002 |
| **ULC179-bin5** | 0.178 | 0.004 | 0.157 | 0.003 | 0.022 | 0.001 | 0.019 | 0.010 |
| **ULC179-bin6** | 0.044 | 0.003 | 0.020 | 0.003 | 0.017 | 0.002 | 0.024 | 0.013 |
| **ULC179-bin7** | 0.010 | 0.000 | 0.003 | | 0.006 | 0.000 | 0.002 | 0.003 |
| **ULC179-nobin** | 0.028 | 0.002 | 0.011 | 0.002 | 0.011 | 0.002 | 0.011 | 0.010 |
| **ULC186-bin1** | 0.006 | 0.001 | 0.002 | 0.020 | 0.002 | 0.047 | 0.003 | 0.004 |
| **ULC186-nobin** | 0.037 | 0.006 | 0.005 | 0.005 | 0.011 | 0.007 | 0.018 | 0.035 |
| **ULC187-bin1** | 0.002 | 0.001 | 0.000 | 0.007 | 0.001 | 0.019 | 0.001 | 0.001 |
| **ULC187-nobin** | 0.060 | 0.029 | 0.075 | 0.004 | 0.085 | 0.001 | 0.053 | 0.015 |

| | ULC146-bin2 | ULC146-bin3 | ULC146-bin4 | ULC146-bin5 | ULC146-bin6 | ULC146-bin7 | ULC146-nobin | ULC165-bin1 |
|---|---|---|---|---|---|---|---|---|
| **ULC335-bin1** | 0.000 | 0.000 | 0.001 | 0.000 | | | 0.011 | 0.000 |
| **ULC335-bin2** | 0.124 | | 0.000 | 0.000 | | | 0.006 | 0.000 |
| **ULC335-bin3** | 0.005 | 0.000 | 0.002 | 0.001 | | 0.000 | 0.002 | 0.001 |
| **ULC335-bin4** | 0.000 | 0.036 | 0.004 | 0.212 | 0.007 | 0.006 | 0.064 | 0.021 |
| **ULC335-nobin** | 0.001 | 0.025 | 0.010 | 0.042 | 0.004 | 0.004 | 0.037 | 0.013 |
| **ULC007-bin1** | 0.000 | 0.001 | 0.001 | 0.000 | 0.000 | 0.000 | 0.015 | 0.001 |
| **ULC007-bin2** | | | | | | | 0.010 | 0.001 |
| **ULC007-nobin** | | 0.008 | 0.000 | 0.001 | | | 0.021 | |
| **ULC027-bin1** | 0.000 | 0.001 | 0.001 | 0.001 | 0.000 | | 0.008 | 0.001 |
| **ULC027-bin2** | 0.000 | 0.021 | 0.003 | 0.027 | 0.004 | 0.002 | 0.027 | 0.010 |
| **ULC027-bin3** | 0.000 | 0.460 | 0.005 | 0.020 | 0.019 | 0.040 | 0.073 | 0.009 |
| **ULC027-bin4** | | 0.354 | 0.001 | 0.012 | 0.014 | 0.135 | 0.075 | 0.010 |
| **ULC027-nobin** | 0.000 | 0.054 | 0.044 | 0.014 | 0.039 | 0.010 | 0.160 | 0.008 |
| **ULC041-bin1** | 0.000 | 0.003 | 0.002 | 0.002 | 0.000 | | 0.010 | 0.003 |
| **ULC041-bin2** | | 0.001 | 0.001 | 0.001 | | 0.001 | 0.003 | 0.005 |
| **ULC041-nobin** | | 0.001 | | 0.001 | | 0.001 | 0.012 | 0.002 |
| **ULC065-bin1** | 0.000 | 0.004 | 0.002 | 0.003 | 0.001 | 0.000 | 0.008 | 0.007 |
| **ULC065-bin2** | 0.000 | 0.015 | 0.011 | 0.014 | 0.002 | 0.002 | 0.017 | 0.488 |
| **ULC065-nobin** | 0.000 | 0.081 | 0.005 | 0.020 | 0.005 | 0.012 | 0.070 | 0.036 |
| **ULC066-bin1** | 0.000 | 0.000 | 0.001 | 0.000 | 0.000 | | 0.007 | 0.000 |
| **ULC066-bin2** | 0.010 | 0.001 | 0.001 | 0.000 | 0.000 | | 0.002 | 0.000 |
| **ULC066-bin3** | | 0.003 | 0.521 | 0.001 | 0.000 | | 0.015 | 0.007 |
| **ULC066-nobin** | 0.001 | 0.055 | 0.118 | 0.016 | 0.004 | 0.008 | 0.048 | 0.010 |
| **ULC068-bin1** | 0.001 | 0.000 | 0.001 | 0.000 | 0.000 | 0.000 | 0.009 | 0.000 |
| **ULC068-bin2** | | 0.001 | 0.001 | | 0.003 | | 0.002 | |
| **ULC068-nobin** | 0.008 | 0.022 | 0.006 | 0.023 | 0.004 | 0.002 | 0.067 | 0.126 |
| **ULC073-bin1** | 0.000 | 0.002 | 0.002 | 0.002 | 0.000 | 0.000 | 0.009 | 0.003 |
| **ULC073-bin2** | 0.000 | 0.004 | 0.453 | 0.002 | 0.000 | 0.001 | 0.013 | 0.009 |
| **ULC073-bin3** | 0.000 | 0.383 | 0.003 | 0.020 | 0.024 | 0.055 | 0.093 | 0.011 |
| **ULC073-bin4** | | 0.059 | 0.002 | 0.013 | 0.033 | 0.010 | 0.211 | 0.011 |
| **ULC073-bin5** | | | 0.001 | | | 0.001 | 0.030 | |
| **ULC073-bin6** | | 0.244 | | 0.005 | 0.001 | 0.001 | 0.107 | 0.003 |
| **ULC073-nobin** | 0.000 | 0.093 | 0.009 | 0.015 | 0.047 | 0.019 | 0.178 | 0.010 |
| **ULC077-bin1** | 0.000 | 0.001 | 0.001 | 0.000 | 0.000 | 0.000 | 0.012 | 0.001 |
| **ULC077-nobin** | 0.000 | 0.023 | 0.184 | 0.017 | 0.003 | 0.003 | 0.034 | 0.146 |
| **ULC082-bin1** | | 0.004 | 0.002 | 0.004 | 0.000 | 0.001 | 0.009 | 0.006 |
| **ULC082-bin2** | 0.000 | 0.020 | 0.003 | 0.053 | 0.004 | 0.002 | 0.035 | 0.008 |
| **ULC082-bin3** | 0.001 | 0.006 | 0.011 | 0.006 | 0.002 | 0.000 | 0.010 | 0.020 |
| **ULC082-bin4** | 0.001 | 0.037 | 0.006 | 0.211 | 0.013 | 0.006 | 0.061 | 0.023 |
| **ULC082-bin5** | | 0.000 | 0.005 | 0.002 | 0.001 | | 0.001 | 0.002 |
| **ULC082-nobin** | 0.000 | 0.014 | 0.009 | 0.054 | 0.002 | 0.002 | 0.035 | 0.013 |
| **ULC084-bin1** | 0.000 | 0.004 | 0.487 | 0.004 | 0.000 | 0.000 | 0.010 | 0.010 |
| **ULC084-bin2** | 0.000 | 0.030 | 0.003 | 0.146 | 0.004 | 0.004 | 0.055 | 0.014 |
| **ULC084-bin3** | 0.000 | 0.005 | 0.002 | 0.004 | 0.001 | 0.001 | 0.009 | 0.008 |
| **ULC084-nobin** | 0.000 | 0.174 | 0.040 | 0.017 | 0.006 | 0.025 | 0.067 | 0.008 |
| **ULC129-bin1** | 0.000 | 0.001 | 0.001 | 0.001 | 0.000 | 0.000 | 0.008 | 0.001 |
| **ULC129-nobin** | 0.002 | 0.023 | 0.097 | 0.010 | 0.003 | 0.003 | 0.040 | 0.007 |
| **ULC146-bin1** | 0.000 | 0.009 | 0.020 | 0.007 | 0.002 | 0.001 | 0.013 | 0.019 |
| **ULC146-bin2** | | 0.000 | 0.000 | 0.000 | | | 0.004 | 0.000 |
| **ULC146-bin3** | 0.000 | | 0.003 | 0.020 | 0.017 | 0.001 | 0.066 | 0.011 |
| **ULC146-bin4** | 0.001 | 0.003 | | 0.004 | 0.001 | 0.000 | 0.007 | 0.009 |
| **ULC146-bin5** | 0.000 | 0.030 | 0.006 | | 0.007 | 0.003 | 0.039 | 0.020 |
| **ULC146-bin6** | | 0.063 | 0.002 | 0.017 | | 0.008 | 0.044 | 0.007 |
| **ULC146-bin7** | | 0.003 | 0.002 | 0.010 | 0.013 | | 0.066 | 0.013 |
| **ULC146-nobin** | 0.002 | 0.029 | 0.003 | 0.012 | 0.005 | 0.004 | | 0.006 |
| **ULC165-bin1** | 0.000 | 0.012 | 0.009 | 0.015 | 0.002 | 0.003 | 0.014 | |
| **ULC165-bin2** | 0.000 | 0.030 | 0.003 | 0.147 | 0.005 | 0.003 | 0.057 | 0.012 |
| **ULC165-bin3** | 0.000 | 0.010 | 0.023 | 0.007 | 0.002 | 0.001 | 0.016 | 0.018 |
| **ULC165-bin4** | 0.000 | 0.000 | 0.001 | | 0.000 | | 0.015 | 0.000 |
| **ULC165-nobin** | 0.000 | 0.032 | 0.005 | 0.011 | 0.008 | 0.005 | 0.046 | 0.009 |
| **ULC179-bin1** | 0.000 | 0.026 | 0.004 | 0.148 | 0.005 | 0.004 | 0.062 | 0.015 |
| **ULC179-bin2** | 0.000 | 0.020 | 0.004 | 0.020 | 0.003 | 0.002 | 0.128 | 0.009 |
| **ULC179-bin3** | 0.000 | 0.060 | 0.003 | 0.025 | 0.012 | 0.014 | 0.053 | 0.015 |
| **ULC179-bin4** | 0.004 | 0.002 | 0.001 | 0.001 | 0.000 | 0.000 | 0.006 | 0.002 |
| **ULC179-bin5** | 0.000 | 0.029 | 0.003 | 0.152 | 0.006 | 0.002 | 0.056 | 0.014 |
| **ULC179-bin6** | | 0.020 | 0.003 | 0.020 | 0.004 | 0.002 | 0.055 | 0.015 |
| **ULC179-bin7** | | 0.009 | | 0.008 | 0.000 | 0.000 | 0.009 | 0.000 |
| **ULC179-nobin** | 0.001 | 0.010 | 0.002 | 0.009 | 0.002 | 0.001 | 0.037 | 0.005 |
| **ULC186-bin1** | 0.000 | 0.002 | 0.002 | 0.002 | 0.000 | 0.000 | 0.009 | 0.003 |
| **ULC186-nobin** | 0.004 | 0.007 | 0.008 | 0.003 | 0.001 | 0.001 | 0.018 | 0.006 |
| **ULC187-bin1** | 0.001 | 0.001 | 0.001 | 0.000 | 0.000 | 0.000 | 0.008 | 0.000 |
| **ULC187-nobin** | 0.001 | 0.162 | 0.029 | 0.032 | 0.012 | 0.021 | 0.098 | 0.010 |

| | ULC165-bin2 | ULC165-bin3 | ULC165-bin4 | ULC165-nobin | ULC179-bin1 | ULC179-bin2 | ULC179-bin3 | ULC179-bin4 |
|---|---|---|---|---|---|---|---|---|
| **ULC335-bin1** | 0.000 | | 0.010 | 0.022 | 0.000 | 0.000 | 0.000 | 0.000 |
| **ULC335-bin2** | 0.000 | | 0.000 | 0.000 | 0.000 | | 0.000 | 0.004 |
| **ULC335-bin3** | 0.000 | 0.000 | 0.000 | 0.001 | 0.001 | 0.000 | 0.001 | 0.005 |
| **ULC335-bin4** | 0.166 | 0.009 | 0.000 | 0.032 | 0.232 | 0.041 | 0.038 | 0.002 |
| **ULC335-nobin** | 0.038 | 0.036 | 0.000 | 0.055 | 0.057 | 0.021 | 0.021 | 0.002 |
| **ULC007-bin1** | 0.000 | 0.000 | 0.018 | 0.039 | 0.001 | 0.001 | 0.001 | 0.001 |
| **ULC007-bin2** | | | 0.007 | 0.017 | | | | |
| **ULC007-nobin** | 0.001 | | 0.007 | 0.043 | | 0.002 | 0.006 | |
| **ULC027-bin1** | 0.001 | 0.001 | 0.009 | 0.024 | 0.001 | 0.001 | 0.001 | 0.001 |
| **ULC027-bin2** | 0.029 | 0.004 | 0.000 | 0.021 | 0.043 | 0.027 | 0.028 | 0.001 |
| **ULC027-bin3** | 0.020 | 0.007 | 0.000 | 0.072 | 0.033 | 0.021 | 0.064 | 0.001 |
| **ULC027-bin4** | 0.019 | 0.006 | | 0.059 | 0.028 | 0.018 | 0.076 | 0.000 |
| **ULC027-nobin** | 0.015 | 0.010 | 0.000 | 0.051 | 0.023 | 0.036 | 0.032 | 0.001 |
| **ULC041-bin1** | 0.002 | 0.002 | 0.012 | 0.030 | 0.004 | 0.004 | 0.003 | 0.001 |
| **ULC041-bin2** | 0.004 | 0.000 | 0.006 | 0.014 | 0.000 | 0.000 | 0.001 | 0.000 |
| **ULC041-nobin** | | 0.002 | | 0.016 | 0.001 | 0.002 | 0.004 | |
| **ULC065-bin1** | 0.004 | 0.003 | 0.003 | 0.012 | 0.006 | 0.005 | 0.005 | 0.001 |
| **ULC065-bin2** | 0.015 | 0.014 | 0.000 | 0.029 | 0.022 | 0.015 | 0.021 | 0.003 |
| **ULC065-nobin** | 0.037 | 0.027 | 0.000 | 0.057 | 0.031 | 0.030 | 0.026 | 0.001 |
| **ULC066-bin1** | 0.000 | 0.000 | 0.008 | 0.017 | 0.000 | 0.001 | 0.001 | 0.000 |
| **ULC066-bin2** | 0.000 | 0.000 | 0.000 | 0.001 | 0.000 | 0.000 | 0.000 | 0.005 |
| **ULC066-bin3** | 0.004 | 0.015 | 0.000 | 0.022 | 0.007 | 0.003 | 0.002 | 0.001 |
| **ULC066-nobin** | 0.017 | 0.009 | 0.000 | 0.034 | 0.025 | 0.025 | 0.023 | 0.001 |
| **ULC068-bin1** | 0.000 | 0.000 | 0.010 | 0.019 | 0.001 | 0.001 | 0.001 | 0.001 |
| **ULC068-bin2** | | | 0.003 | 0.007 | | | 0.001 | |
| **ULC068-nobin** | 0.037 | 0.013 | 0.000 | 0.040 | 0.035 | 0.034 | 0.027 | 0.002 |
| **ULC073-bin1** | 0.003 | 0.001 | 0.010 | 0.028 | 0.003 | 0.003 | 0.003 | 0.001 |
| **ULC073-bin2** | 0.004 | 0.016 | 0.001 | 0.015 | 0.005 | 0.004 | 0.003 | 0.001 |
| **ULC073-bin3** | 0.022 | 0.008 | | 0.075 | 0.037 | 0.025 | 0.064 | 0.001 |
| **ULC073-bin4** | 0.019 | 0.004 | | 0.077 | 0.036 | 0.022 | 0.055 | 0.000 |
| **ULC073-bin5** | | 0.001 | 0.010 | 0.018 | 0.001 | 0.001 | | 0.001 |
| **ULC073-bin6** | 0.006 | 0.003 | | 0.066 | 0.015 | 0.006 | 0.007 | |
| **ULC073-nobin** | 0.015 | 0.021 | 0.000 | 0.069 | 0.024 | 0.022 | 0.039 | 0.001 |
| **ULC077-bin1** | 0.001 | 0.000 | 0.016 | 0.036 | 0.001 | 0.000 | 0.000 | 0.000 |
| **ULC077-nobin** | 0.040 | 0.014 | 0.001 | 0.030 | 0.025 | 0.012 | 0.012 | 0.001 |
| **ULC082-bin1** | 0.004 | 0.004 | 0.004 | 0.014 | 0.007 | 0.006 | 0.005 | 0.001 |
| **ULC082-bin2** | 0.051 | 0.007 | | 0.024 | 0.059 | 0.029 | 0.025 | 0.001 |
| **ULC082-bin3** | 0.006 | 0.024 | 0.000 | 0.015 | 0.011 | 0.009 | 0.009 | 0.001 |
| **ULC082-bin4** | 0.169 | 0.004 | 0.000 | 0.040 | 0.280 | 0.042 | 0.038 | 0.001 |
| **ULC082-bin5** | | 0.008 | | 0.002 | 0.002 | 0.001 | | 0.000 |
| **ULC082-nobin** | 0.057 | 0.030 | 0.000 | 0.048 | 0.075 | 0.028 | 0.016 | 0.001 |
| **ULC084-bin1** | 0.004 | 0.017 | 0.001 | 0.015 | 0.005 | 0.005 | 0.004 | 0.001 |
| **ULC084-bin2** | 0.491 | 0.005 | 0.000 | 0.035 | 0.218 | 0.030 | 0.034 | 0.002 |
| **ULC084-bin3** | 0.004 | 0.003 | 0.003 | 0.013 | 0.006 | 0.005 | 0.006 | 0.001 |
| **ULC084-nobin** | 0.031 | 0.010 | 0.000 | 0.050 | 0.028 | 0.023 | 0.031 | 0.001 |
| **ULC129-bin1** | 0.001 | 0.001 | 0.009 | 0.023 | 0.001 | 0.001 | 0.002 | 0.001 |
| **ULC129-nobin** | 0.011 | 0.019 | 0.001 | 0.033 | 0.016 | 0.013 | 0.012 | 0.003 |
| **ULC146-bin1** | 0.008 | 0.196 | 0.000 | 0.243 | 0.013 | 0.014 | 0.012 | 0.001 |
| **ULC146-bin2** | 0.000 | 0.000 | 0.000 | 0.000 | 0.000 | 0.000 | 0.000 | 0.004 |
| **ULC146-bin3** | 0.025 | 0.006 | 0.000 | 0.073 | 0.032 | 0.027 | 0.066 | 0.002 |
| **ULC146-bin4** | 0.002 | 0.014 | 0.000 | 0.015 | 0.005 | 0.005 | 0.004 | 0.001 |
| **ULC146-bin5** | 0.189 | 0.006 | | 0.039 | 0.272 | 0.041 | 0.041 | 0.002 |
| **ULC146-bin6** | 0.017 | 0.003 | | 0.067 | 0.024 | 0.016 | 0.048 | 0.000 |
| **ULC146-bin7** | 0.015 | 0.005 | | 0.064 | 0.026 | 0.014 | 0.090 | 0.001 |
| **ULC146-nobin** | 0.021 | 0.004 | 0.002 | 0.046 | 0.032 | 0.076 | 0.027 | 0.003 |
| **ULC165-bin1** | 0.012 | 0.012 | 0.000 | 0.021 | 0.021 | 0.013 | 0.018 | 0.002 |
| **ULC165-bin2** | | 0.006 | 0.000 | 0.027 | 0.223 | 0.035 | 0.034 | 0.002 |
| **ULC165-bin3** | 0.009 | | 0.000 | 0.017 | 0.014 | 0.011 | 0.010 | 0.002 |
| **ULC165-bin4** | 0.001 | 0.000 | | 0.013 | 0.000 | 0.001 | 0.000 | 0.000 |
| **ULC165-nobin** | 0.010 | 0.004 | 0.002 | | 0.017 | 0.021 | 0.029 | 0.001 |
| **ULC179-bin1** | 0.156 | 0.006 | 0.000 | 0.031 | | 0.037 | 0.032 | 0.002 |
| **ULC179-bin2** | 0.022 | 0.005 | 0.000 | 0.036 | 0.033 | | 0.024 | 0.001 |
| **ULC179-bin3** | 0.026 | 0.006 | 0.000 | 0.059 | 0.038 | 0.029 | | 0.002 |
| **ULC179-bin4** | 0.002 | 0.001 | 0.000 | 0.002 | 0.002 | 0.002 | 0.002 | |
| **ULC179-bin5** | 0.156 | 0.004 | | 0.027 | 0.196 | 0.026 | 0.029 | 0.001 |
| **ULC179-bin6** | 0.021 | 0.009 | | 0.025 | 0.035 | 0.048 | 0.020 | 0.002 |
| **ULC179-bin7** | 0.007 | 0.001 | | 0.012 | 0.008 | 0.002 | 0.021 | 0.000 |
| **ULC179-nobin** | 0.012 | 0.004 | 0.001 | 0.019 | 0.015 | 0.023 | 0.012 | 0.003 |
| **ULC186-bin1** | 0.002 | 0.002 | 0.011 | 0.028 | 0.003 | 0.003 | 0.003 | 0.001 |
| **ULC186-nobin** | 0.004 | 0.017 | 0.002 | 0.029 | 0.007 | 0.009 | 0.008 | 0.002 |
| **ULC187-bin1** | 0.001 | 0.000 | 0.009 | 0.017 | 0.000 | 0.000 | 0.001 | 0.000 |
| **ULC187-nobin** | 0.076 | 0.006 | 0.000 | 0.051 | 0.048 | 0.029 | 0.038 | 0.002 |

| | ULC179-bin5 | ULC179-bin6 | ULC179-bin7 | ULC179-nobin | ULC186-bin1 | ULC186-nobin | ULC187-bin1 | ULC187-nobin |
|---|---|---|---|---|---|---|---|---|
| **ULC335-bin1** | 0.000 | | | 0.005 | 0.023 | 0.003 | 0.030 | 0.000 |
| **ULC335-bin2** | | | | 0.002 | 0.000 | 0.004 | 0.001 | 0.000 |
| **ULC335-bin3** | 0.000 | 0.000 | | 0.003 | 0.001 | 0.002 | 0.001 | 0.001 |
| **ULC335-bin4** | 0.064 | 0.007 | 0.001 | 0.027 | 0.004 | 0.004 | 0.001 | 0.029 |
| **ULC335-nobin** | 0.018 | 0.003 | 0.000 | 0.017 | 0.003 | 0.013 | 0.001 | 0.014 |
| **ULC007-bin1** | 0.000 | 0.000 | | 0.006 | 0.040 | 0.003 | 0.027 | 0.001 |
| **ULC007-bin2** | | | | 0.004 | 0.013 | 0.005 | 0.001 | |
| **ULC007-nobin** | | | | 0.019 | 0.016 | 0.013 | 0.022 | 0.011 |
| **ULC027-bin1** | 0.000 | 0.000 | 0.000 | 0.004 | 0.050 | 0.005 | 0.021 | 0.001 |
| **ULC027-bin2** | 0.014 | 0.003 | 0.000 | 0.018 | 0.002 | 0.003 | 0.001 | 0.012 |
| **ULC027-bin3** | 0.010 | 0.003 | 0.000 | 0.019 | 0.002 | 0.003 | 0.001 | 0.104 |
| **ULC027-bin4** | 0.007 | 0.004 | | 0.009 | 0.002 | 0.003 | | 0.079 |
| **ULC027-nobin** | 0.008 | 0.004 | 0.001 | 0.019 | 0.004 | 0.005 | 0.001 | 0.039 |
| **ULC041-bin1** | 0.001 | 0.001 | | 0.005 | 0.249 | 0.020 | 0.018 | 0.001 |
| **ULC041-bin2** | | | | 0.001 | 0.063 | 0.014 | 0.012 | 0.001 |
| **ULC041-nobin** | | 0.001 | | 0.009 | 0.097 | 0.028 | 0.007 | 0.007 |
| **ULC065-bin1** | 0.002 | 0.001 | | 0.006 | 0.034 | 0.003 | 0.011 | 0.002 |
| **ULC065-bin2** | 0.008 | 0.002 | | 0.011 | 0.005 | 0.005 | 0.001 | 0.009 |
| **ULC065-nobin** | 0.010 | 0.005 | 0.001 | 0.024 | 0.003 | 0.007 | 0.001 | 0.042 |
| **ULC066-bin1** | 0.000 | 0.000 | | 0.003 | 0.017 | 0.002 | 0.170 | 0.001 |
| **ULC066-bin2** | | | | 0.003 | 0.000 | 0.003 | 0.001 | 0.000 |
| **ULC066-bin3** | 0.003 | 0.000 | | 0.003 | 0.004 | 0.005 | 0.002 | 0.020 |
| **ULC066-nobin** | 0.008 | 0.003 | 0.000 | 0.017 | 0.003 | 0.005 | 0.001 | 0.028 |
| **ULC068-bin1** | 0.000 | 0.000 | | 0.004 | 0.019 | 0.002 | 0.185 | 0.001 |
| **ULC068-bin2** | | | | 0.001 | 0.001 | 0.001 | 0.131 | 0.149 |
| **ULC068-nobin** | 0.011 | 0.005 | 0.000 | 0.024 | 0.003 | 0.008 | 0.009 | 0.032 |
| **ULC073-bin1** | 0.001 | 0.000 | | 0.005 | 0.236 | 0.020 | 0.018 | 0.001 |
| **ULC073-bin2** | 0.001 | 0.001 | 0.000 | 0.004 | 0.002 | 0.004 | 0.001 | 0.021 |
| **ULC073-bin3** | 0.012 | 0.004 | 0.001 | 0.016 | 0.003 | 0.002 | 0.000 | 0.087 |
| **ULC073-bin4** | 0.010 | 0.005 | 0.003 | 0.017 | 0.002 | 0.001 | 0.000 | 0.043 |
| **ULC073-bin5** | | | | 0.002 | 0.075 | 0.014 | 0.008 | |
| **ULC073-bin6** | 0.003 | 0.001 | 0.003 | 0.014 | 0.002 | 0.005 | | 0.142 |
| **ULC073-nobin** | 0.007 | 0.004 | 0.001 | 0.017 | 0.012 | 0.007 | 0.001 | 0.044 |
| **ULC077-bin1** | 0.000 | 0.000 | 0.000 | 0.005 | 0.039 | 0.003 | 0.028 | 0.001 |
| **ULC077-nobin** | 0.008 | 0.002 | 0.000 | 0.011 | 0.004 | 0.007 | 0.001 | 0.023 |
| **ULC082-bin1** | 0.002 | 0.001 | 0.000 | 0.006 | 0.038 | 0.004 | 0.010 | 0.002 |
| **ULC082-bin2** | 0.023 | 0.007 | 0.001 | 0.019 | 0.005 | 0.003 | 0.001 | 0.017 |
| **ULC082-bin3** | 0.002 | 0.003 | | 0.008 | 0.004 | 0.005 | 0.001 | 0.004 |
| **ULC082-bin4** | 0.096 | 0.004 | 0.001 | 0.022 | 0.004 | 0.002 | 0.001 | 0.032 |
| **ULC082-bin5** | | | | 0.001 | 0.001 | | | |
| **ULC082-nobin** | 0.023 | 0.003 | 0.000 | 0.017 | 0.003 | 0.007 | 0.001 | 0.013 |
| **ULC084-bin1** | 0.002 | 0.001 | 0.000 | 0.003 | 0.002 | 0.004 | 0.001 | 0.023 |
| **ULC084-bin2** | 0.072 | 0.004 | 0.000 | 0.024 | 0.003 | 0.003 | 0.001 | 0.057 |
| **ULC084-bin3** | 0.002 | 0.001 | | 0.005 | 0.035 | 0.004 | 0.011 | 0.003 |
| **ULC084-nobin** | 0.008 | 0.003 | 0.001 | 0.020 | 0.003 | 0.007 | 0.001 | 0.053 |
| **ULC129-bin1** | 0.000 | 0.000 | 0.000 | 0.003 | 0.050 | 0.004 | 0.018 | 0.001 |
| **ULC129-nobin** | 0.004 | 0.003 | 0.000 | 0.012 | 0.003 | 0.006 | 0.001 | 0.020 |
| **ULC146-bin1** | 0.003 | 0.002 | 0.000 | 0.012 | 0.004 | 0.015 | 0.001 | 0.007 |
| **ULC146-bin2** | 0.000 | | | 0.001 | 0.000 | 0.002 | 0.001 | 0.000 |
| **ULC146-bin3** | 0.011 | 0.004 | 0.002 | 0.017 | 0.003 | 0.004 | 0.001 | 0.102 |
| **ULC146-bin4** | 0.001 | 0.001 | | 0.003 | 0.003 | 0.005 | 0.001 | 0.020 |
| **ULC146-bin5** | 0.088 | 0.005 | 0.002 | 0.025 | 0.003 | 0.003 | 0.000 | 0.031 |
| **ULC146-bin6** | 0.008 | 0.002 | 0.000 | 0.012 | 0.002 | 0.001 | 0.000 | 0.029 |
| **ULC146-bin7** | 0.004 | 0.003 | 0.000 | 0.011 | 0.002 | 0.003 | 0.001 | 0.088 |
| **ULC146-nobin** | 0.009 | 0.005 | 0.001 | 0.030 | 0.005 | 0.005 | 0.004 | 0.026 |
| **ULC165-bin1** | 0.006 | 0.003 | 0.000 | 0.009 | 0.004 | 0.004 | 0.000 | 0.007 |
| **ULC165-bin2** | 0.070 | 0.004 | 0.001 | 0.025 | 0.004 | 0.003 | 0.001 | 0.056 |
| **ULC165-bin3** | 0.003 | 0.003 | 0.000 | 0.013 | 0.005 | 0.017 | 0.001 | 0.007 |
| **ULC165-bin4** | | | | 0.005 | 0.041 | 0.003 | 0.032 | 0.000 |
| **ULC165-nobin** | 0.005 | 0.002 | 0.001 | 0.014 | 0.015 | 0.007 | 0.008 | 0.014 |
| **ULC179-bin1** | 0.049 | 0.005 | 0.001 | 0.022 | 0.003 | 0.003 | 0.000 | 0.024 |
| **ULC179-bin2** | 0.008 | 0.007 | 0.000 | 0.032 | 0.003 | 0.004 | 0.000 | 0.013 |
| **ULC179-bin3** | 0.010 | 0.003 | 0.002 | 0.019 | 0.003 | 0.004 | 0.001 | 0.021 |
| **ULC179-bin4** | 0.000 | 0.000 | 0.000 | 0.005 | 0.002 | 0.001 | 0.000 | 0.001 |
| **ULC179-bin5** | | 0.006 | 0.001 | 0.023 | 0.003 | 0.002 | 0.001 | 0.024 |
| **ULC179-bin6** | 0.013 | | 0.001 | 0.022 | 0.003 | 0.008 | 0.001 | 0.013 |
| **ULC179-bin7** | 0.003 | 0.001 | | 0.020 | | | 0.001 | 0.003 |
| **ULC179-nobin** | 0.005 | 0.003 | 0.001 | | 0.004 | 0.004 | 0.002 | 0.008 |
| **ULC186-bin1** | 0.001 | 0.000 | | 0.006 | | 0.007 | 0.019 | 0.002 |
| **ULC186-nobin** | 0.002 | 0.002 | | 0.012 | 0.020 | | 0.003 | 0.010 |
| **ULC187-bin1** | 0.000 | 0.000 | 0.000 | 0.003 | 0.021 | 0.002 | | 0.005 |
| **ULC187-nobin** | 0.015 | 0.004 | 0.001 | 0.021 | 0.004 | 0.010 | 0.010 | |

| | ULC335-bin1 | ULC335-bin2 | ULC335-bin3 | ULC335-bin4 | ULC335-nobin | ULC007-bin1 | ULC007-bin2 | ULC007-nobin |
|---|---|---|---|---|---|---|---|---|
| **ULC335-bin1** | | | | | | | | |
| **ULC335-bin2** | 0.562 | | | | | | | |
| **ULC335-bin3** | 0.000 | 0.000 | | | | | | |
| **ULC335-bin4** | 0.000 | 0.000 | 0.000 | | | | | |
| **ULC335-nobin** | 0.372 | 0.157 | 0.000 | 0.000 | | | | |
| **ULC007-bin1** | 0.000 | 0.000 | 0.000 | 0.000 | 0.000 | | | |
| **ULC007-bin2** | 0.000 | 0.000 | 0.000 | 0.000 | 0.000 | 0.000 | | |
| **ULC007-nobin** | 0.000 | 0.000 | 0.000 | 0.000 | 0.213 | 1.063 | 0.106 | |
| **ULC027-bin1** | 0.000 | 0.000 | 0.000 | 0.000 | 0.000 | 0.000 | 0.000 | 0.106 |
| **ULC027-bin2** | 0.000 | 0.000 | 0.000 | 0.000 | 0.000 | 0.000 | 0.000 | 0.106 |
| **ULC027-bin3** | 0.000 | 0.000 | 0.000 | 0.000 | 0.000 | 0.000 | 0.000 | 0.106 |
| **ULC027-bin4** | 0.000 | 0.000 | 0.000 | 0.000 | 0.000 | 0.000 | 0.000 | 0.000 |
| **ULC027-nobin** | 0.000 | 0.000 | 0.000 | 0.000 | 0.008 | 0.005 | 0.000 | 0.106 |
| **ULC041-bin1** | 0.000 | 0.000 | 0.000 | 0.000 | 0.000 | 0.000 | 0.000 | 0.850 |
| **ULC041-bin2** | 0.000 | 0.000 | 0.000 | 0.000 | 0.000 | 0.000 | 0.000 | 0.106 |
| **ULC041-nobin** | 0.000 | 0.000 | 0.000 | 0.000 | 0.071 | 0.142 | 0.000 | 0.106 |
| **ULC065-bin1** | 0.000 | 0.000 | 0.000 | 0.000 | 0.000 | 0.000 | 0.000 | 0.531 |
| **ULC065-bin2** | 0.000 | 0.000 | 0.000 | 0.000 | 0.010 | 0.000 | 0.000 | 0.000 |
| **ULC065-nobin** | 0.000 | 0.000 | 0.000 | 0.000 | 0.011 | 0.010 | 0.000 | 0.319 |
| **ULC066-bin1** | 0.000 | 0.000 | 0.000 | 0.000 | 0.000 | 0.000 | 0.000 | 2.019 |
| **ULC066-bin2** | 0.000 | 0.000 | 0.000 | 0.000 | 0.000 | 0.000 | 0.000 | 0.213 |
| **ULC066-bin3** | 0.000 | 0.000 | 0.000 | 0.000 | 0.000 | 0.000 | 0.000 | 0.000 |
| **ULC066-nobin** | 0.000 | 0.000 | 0.000 | 0.000 | 0.006 | 0.010 | 0.000 | 0.425 |
| **ULC068-bin1** | 0.000 | 0.000 | 0.000 | 0.000 | 0.000 | 0.000 | 0.000 | 0.850 |
| **ULC068-bin2** | 0.000 | 0.000 | 0.000 | 0.000 | 0.000 | 0.000 | 0.000 | 0.000 |
| **ULC068-nobin** | 0.000 | 0.000 | 0.000 | 0.000 | 0.008 | 0.025 | 0.000 | 0.106 |
| **ULC073-bin1** | 0.000 | 0.000 | 0.000 | 0.000 | 0.000 | 0.000 | 0.000 | 0.319 |
| **ULC073-bin2** | 0.000 | 0.000 | 0.000 | 0.000 | 0.000 | 0.000 | 0.000 | 0.106 |
| **ULC073-bin3** | 0.000 | 0.000 | 0.000 | 0.000 | 0.000 | 0.000 | 0.000 | 0.106 |
| **ULC073-bin4** | 0.000 | 0.000 | 0.000 | 0.000 | 0.000 | 0.000 | 0.000 | 0.000 |
| **ULC073-bin5** | 0.000 | 0.000 | 0.000 | 0.000 | 0.000 | 0.000 | 0.000 | 0.000 |
| **ULC073-bin6** | 0.000 | 0.000 | 0.000 | 0.000 | 0.000 | 0.000 | 0.000 | 0.000 |
| **ULC073-nobin** | 0.000 | 0.000 | 0.000 | 0.000 | 0.007 | 0.010 | 0.000 | 0.213 |
| **ULC077-bin1** | 0.000 | 0.000 | 0.000 | 0.000 | 0.000 | 0.000 | 0.000 | 0.531 |
| **ULC077-nobin** | 0.000 | 0.000 | 0.000 | 0.000 | 0.009 | 0.010 | 0.000 | 3.613 |
| **ULC082-bin1** | 0.000 | 0.000 | 0.000 | 0.000 | 0.000 | 0.000 | 0.000 | 0.000 |
| **ULC082-bin2** | 0.000 | 0.000 | 0.000 | 0.025 | 0.011 | 0.000 | 0.000 | 0.000 |
| **ULC082-bin3** | 0.000 | 0.000 | 0.000 | 0.000 | 0.000 | 0.000 | 0.000 | 0.000 |
| **ULC082-bin4** | 0.000 | 0.000 | 0.000 | 0.000 | 0.000 | 0.000 | 0.000 | 0.000 |
| **ULC082-bin5** | 0.000 | 0.000 | 0.000 | 0.000 | 0.000 | 0.000 | 0.000 | 0.000 |
| **ULC082-nobin** | 0.000 | 0.000 | 0.000 | 0.025 | 0.048 | 0.000 | 0.000 | 0.213 |
| **ULC084-bin1** | 0.000 | 0.000 | 0.000 | 0.000 | 0.000 | 0.000 | 0.000 | 0.000 |
| **ULC084-bin2** | 0.000 | 0.000 | 0.000 | 0.000 | 0.000 | 0.000 | 0.000 | 0.000 |
| **ULC084-bin3** | 0.000 | 0.000 | 0.000 | 0.000 | 0.000 | 0.000 | 0.000 | 0.000 |
| **ULC084-nobin** | 0.000 | 0.000 | 0.009 | 0.000 | 0.092 | 0.000 | 0.000 | 0.213 |
| **ULC129-bin1** | 0.000 | 0.000 | 0.000 | 0.000 | 0.000 | 0.000 | 0.000 | 0.000 |
| **ULC129-nobin** | 0.005 | 0.000 | 0.000 | 0.000 | 0.048 | 0.000 | 0.000 | 0.213 |
| **ULC146-bin1** | 0.000 | 0.000 | 0.000 | 0.000 | 0.046 | 0.000 | 0.000 | 0.000 |
| **ULC146-bin2** | 0.000 | 0.000 | 0.000 | 0.000 | 0.000 | 0.000 | 0.000 | 0.000 |
| **ULC146-bin3** | 0.000 | 0.000 | 0.000 | 0.000 | 0.000 | 0.000 | 0.000 | 0.106 |
| **ULC146-bin4** | 0.000 | 0.000 | 0.000 | 0.000 | 0.000 | 0.000 | 0.000 | 0.106 |
| **ULC146-bin5** | 0.000 | 0.000 | 0.000 | 0.000 | 0.011 | 0.000 | 0.000 | 0.000 |
| **ULC146-bin6** | 0.000 | 0.000 | 0.000 | 0.000 | 0.000 | 0.000 | 0.000 | 0.000 |
| **ULC146-bin7** | 0.000 | 0.000 | 0.000 | 0.000 | 0.000 | 0.000 | 0.000 | 0.000 |
| **ULC146-nobin** | 0.000 | 0.000 | 0.000 | 0.000 | 0.065 | 0.000 | 0.000 | 0.213 |
| **ULC165-bin1** | 0.000 | 0.000 | 0.000 | 0.000 | 0.008 | 0.000 | 0.000 | 0.000 |
| **ULC165-bin2** | 0.000 | 0.000 | 0.000 | 0.000 | 0.000 | 0.000 | 0.000 | 0.000 |
| **ULC165-bin3** | 0.000 | 0.000 | 0.000 | 0.000 | 0.064 | 0.000 | 0.000 | 0.000 |
| **ULC165-bin4** | 0.000 | 0.000 | 0.000 | 0.000 | 0.000 | 0.000 | 0.000 | 0.000 |
| **ULC165-nobin** | 0.005 | 0.017 | 0.000 | 0.000 | 0.043 | 0.000 | 0.000 | 0.319 |
| **ULC179-bin1** | 0.000 | 0.000 | 0.000 | 0.000 | 0.000 | 0.000 | 0.000 | 0.000 |
| **ULC179-bin2** | 0.000 | 0.000 | 0.000 | 0.000 | 0.000 | 0.000 | 0.000 | 0.000 |
| **ULC179-bin3** | 0.000 | 0.000 | 0.000 | 0.000 | 0.000 | 0.000 | 0.000 | 0.000 |
| **ULC179-bin4** | 0.000 | 0.000 | 0.000 | 0.000 | 0.000 | 0.000 | 0.000 | 0.000 |
| **ULC179-bin5** | 0.000 | 0.000 | 0.000 | 0.000 | 0.000 | 0.000 | 0.000 | 0.000 |
| **ULC179-bin6** | 0.000 | 0.000 | 0.000 | 0.000 | 0.000 | 0.000 | 0.000 | 0.000 |
| **ULC179-bin7** | 0.000 | 0.000 | 0.000 | 0.000 | 0.000 | 0.000 | 0.000 | 0.000 |
| **ULC179-nobin** | 0.000 | 0.000 | 0.000 | 0.000 | 0.014 | 0.000 | 0.000 | 0.213 |
| **ULC186-bin1** | 0.000 | 0.000 | 0.000 | 0.000 | 0.005 | 0.000 | 0.000 | 0.000 |
| **ULC186-nobin** | 0.022 | 0.022 | 0.000 | 0.000 | 0.345 | 0.000 | 0.000 | 0.213 |
| **ULC187-bin1** | 0.000 | 0.000 | 0.000 | 0.000 | 0.006 | 0.000 | 0.000 | 0.000 |
| **ULC187-nobin** | 0.000 | 0.000 | 0.000 | 0.000 | 0.112 | 0.000 | 0.000 | 0.213 |

| | ULC027-bin1 | ULC027-bin2 | ULC027-bin3 | ULC027-bin4 | ULC027-nobin | ULC041-bin1 | ULC041-bin2 | ULC041-nobin |
|---|---|---|---|---|---|---|---|---|
| **ULC335-bin1** | | | | | | | | |
| **ULC335-bin2** | | | | | | | | |
| **ULC335-bin3** | | | | | | | | |
| **ULC335-bin4** | | | | | | | | |
| **ULC335-nobin** | | | | | | | | |
| **ULC007-bin1** | | | | | | | | |
| **ULC007-bin2** | | | | | | | | |
| **ULC007-nobin** | | | | | | | | |
| **ULC027-bin1** | | | | | | | | |
| **ULC027-bin2** | 0.000 | | | | | | | |
| **ULC027-bin3** | 0.000 | 0.000 | | | | | | |
| **ULC027-bin4** | 0.000 | 0.000 | 0.000 | | | | | |
| **ULC027-nobin** | 0.059 | 0.007 | 0.000 | 0.000 | | | | |
| **ULC041-bin1** | 0.000 | 0.000 | 0.000 | 0.000 | 0.005 | | | |
| **ULC041-bin2** | 0.000 | 0.000 | 0.000 | 0.000 | 0.000 | 0.145 | | |
| **ULC041-nobin** | 0.000 | 0.071 | 0.071 | 0.000 | 0.212 | 0.921 | 0.496 | |
| **ULC065-bin1** | 0.000 | 0.000 | 0.000 | 0.000 | 0.024 | 0.000 | 0.000 | 0.071 |
| **ULC065-bin2** | 0.000 | 0.000 | 0.000 | 0.000 | 0.000 | 0.000 | 0.000 | 0.000 |
| **ULC065-nobin** | 0.005 | 0.007 | 19.065 | 14.226 | 2.087 | 0.005 | 0.097 | 0.212 |
| **ULC066-bin1** | 0.000 | 0.000 | 0.000 | 0.000 | 0.019 | 0.000 | 0.000 | 0.850 |
| **ULC066-bin2** | 0.000 | 0.000 | 0.000 | 0.000 | 0.000 | 0.000 | 0.000 | 0.000 |
| **ULC066-bin3** | 0.000 | 0.000 | 0.000 | 0.000 | 14.251 | 0.000 | 0.000 | 0.000 |
| **ULC066-nobin** | 0.005 | 36.297 | 15.940 | 11.758 | 3.070 | 0.005 | 0.000 | 0.142 |
| **ULC068-bin1** | 0.000 | 0.000 | 0.000 | 0.000 | 0.011 | 0.000 | 0.000 | 0.354 |
| **ULC068-bin2** | 0.000 | 0.000 | 0.000 | 0.000 | 0.000 | 0.000 | 0.000 | 0.000 |
| **ULC068-nobin** | 0.000 | 3.580 | 0.114 | 0.028 | 1.047 | 0.000 | 0.000 | 0.071 |
| **ULC073-bin1** | 0.006 | 0.000 | 0.000 | 0.000 | 0.006 | 19.619 | 7.081 | 5.312 |
| **ULC073-bin2** | 0.000 | 0.000 | 0.000 | 0.000 | 16.106 | 0.000 | 0.000 | 0.071 |
| **ULC073-bin3** | 0.000 | 0.000 | 55.001 | 43.844 | 7.815 | 0.000 | 0.000 | 0.000 |
| **ULC073-bin4** | 0.000 | 0.000 | 0.000 | 0.967 | 21.787 | 0.000 | 0.000 | 0.071 |
| **ULC073-bin5** | 0.000 | 0.000 | 0.000 | 0.000 | 0.890 | 4.982 | 6.495 | 1.601 |
| **ULC073-bin6** | 0.000 | 0.091 | 82.743 | 0.000 | 3.633 | 0.000 | 0.000 | 0.000 |
| **ULC073-nobin** | 0.000 | 0.007 | 10.848 | 19.440 | 8.979 | 1.057 | 1.552 | 4.462 |
| **ULC077-bin1** | 0.000 | 0.000 | 0.000 | 0.000 | 0.000 | 0.000 | 0.000 | 0.142 |
| **ULC077-nobin** | 0.000 | 0.140 | 3.296 | 2.413 | 2.361 | 0.000 | 0.000 | 0.142 |
| **ULC082-bin1** | 0.000 | 0.000 | 0.000 | 0.000 | 0.000 | 0.000 | 0.000 | 0.000 |
| **ULC082-bin2** | 0.000 | 0.000 | 0.000 | 0.000 | 0.000 | 0.000 | 0.000 | 0.000 |
| **ULC082-bin3** | 0.000 | 0.000 | 0.000 | 0.000 | 0.000 | 0.000 | 0.000 | 0.000 |
| **ULC082-bin4** | 0.000 | 0.000 | 0.000 | 0.000 | 0.000 | 0.000 | 0.000 | 0.000 |
| **ULC082-bin5** | 0.000 | 0.000 | 0.000 | 0.000 | 0.000 | 0.000 | 0.000 | 0.000 |
| **ULC082-nobin** | 0.000 | 0.000 | 0.000 | 0.000 | 0.077 | 0.000 | 0.000 | 0.071 |
| **ULC084-bin1** | 0.000 | 0.000 | 0.000 | 0.000 | 15.489 | 0.000 | 0.000 | 0.071 |
| **ULC084-bin2** | 0.000 | 0.000 | 0.000 | 0.000 | 0.000 | 0.000 | 0.000 | 0.000 |
| **ULC084-bin3** | 0.000 | 0.000 | 0.000 | 0.000 | 0.000 | 0.000 | 0.000 | 0.000 |
| **ULC084-nobin** | 0.000 | 0.000 | 23.710 | 19.052 | 2.571 | 0.000 | 0.000 | 0.071 |
| **ULC129-bin1** | 0.016 | 0.000 | 0.000 | 0.000 | 0.053 | 0.000 | 0.000 | 0.000 |
| **ULC129-nobin** | 0.011 | 24.048 | 4.455 | 2.551 | 1.769 | 0.005 | 0.000 | 0.071 |
| **ULC146-bin1** | 0.000 | 0.000 | 0.000 | 0.000 | 2.796 | 0.000 | 0.000 | 0.071 |
| **ULC146-bin2** | 0.000 | 0.000 | 0.000 | 0.000 | 0.000 | 0.000 | 0.000 | 0.000 |
| **ULC146-bin3** | 0.000 | 0.014 | 81.733 | 60.094 | 6.862 | 0.000 | 0.000 | 0.071 |
| **ULC146-bin4** | 0.000 | 0.000 | 0.000 | 0.000 | 16.060 | 0.000 | 0.000 | 0.071 |
| **ULC146-bin5** | 0.000 | 0.000 | 0.000 | 0.000 | 0.190 | 0.000 | 0.000 | 0.000 |
| **ULC146-bin6** | 0.000 | 0.000 | 1.798 | 0.305 | 36.421 | 0.000 | 0.000 | 0.000 |
| **ULC146-bin7** | 0.000 | 0.000 | 32.378 | 38.737 | 9.987 | 0.000 | 0.000 | 0.000 |
| **ULC146-nobin** | 0.000 | 0.014 | 1.415 | 1.248 | 6.304 | 0.000 | 0.000 | 0.071 |
| **ULC165-bin1** | 0.000 | 0.000 | 0.000 | 0.000 | 0.000 | 0.000 | 0.000 | 0.000 |
| **ULC165-bin2** | 0.000 | 0.000 | 0.000 | 0.000 | 0.000 | 0.000 | 0.000 | 0.000 |
| **ULC165-bin3** | 0.000 | 0.000 | 0.000 | 0.000 | 2.723 | 0.000 | 0.000 | 0.071 |
| **ULC165-bin4** | 0.000 | 0.000 | 0.000 | 0.000 | 0.000 | 0.000 | 0.000 | 0.000 |
| **ULC165-nobin** | 0.000 | 0.007 | 4.370 | 2.856 | 1.153 | 0.000 | 0.000 | 0.071 |
| **ULC179-bin1** | 0.000 | 0.000 | 0.000 | 0.000 | 0.000 | 0.000 | 0.000 | 0.000 |
| **ULC179-bin2** | 0.000 | 0.000 | 0.000 | 0.000 | 0.000 | 0.000 | 0.000 | 0.000 |
| **ULC179-bin3** | 0.000 | 0.000 | 0.000 | 0.000 | 0.007 | 0.000 | 0.000 | 0.000 |
| **ULC179-bin4** | 0.000 | 0.000 | 0.000 | 0.000 | 0.000 | 0.000 | 0.000 | 0.000 |
| **ULC179-bin5** | 0.000 | 0.000 | 0.000 | 0.000 | 0.000 | 0.000 | 0.000 | 0.000 |
| **ULC179-bin6** | 0.000 | 0.000 | 0.000 | 0.000 | 0.000 | 0.000 | 0.000 | 0.000 |
| **ULC179-bin7** | 0.000 | 0.000 | 0.000 | 0.000 | 0.000 | 0.000 | 0.000 | 0.000 |
| **ULC179-nobin** | 0.000 | 0.014 | 0.019 | 0.000 | 0.007 | 0.000 | 0.000 | 0.071 |
| **ULC186-bin1** | 0.000 | 0.000 | 0.000 | 0.000 | 0.000 | 0.000 | 0.000 | 0.000 |
| **ULC186-nobin** | 0.000 | 0.000 | 0.000 | 0.000 | 0.022 | 0.000 | 0.000 | 0.212 |
| **ULC187-bin1** | 0.000 | 0.000 | 0.000 | 0.000 | 0.000 | 0.000 | 0.000 | 0.000 |
| **ULC187-nobin** | 0.000 | 0.000 | 15.438 | 10.261 | 3.327 | 0.010 | 0.000 | 0.071 |

| | ULC065-bin1 | ULC065-bin2 | ULC065-nobin | ULC066-bin1 | ULC066-bin2 | ULC066-bin3 | ULC066-nobin | ULC068-bin1 |
|---|---|---|---|---|---|---|---|---|
| **ULC335-bin1** | | | | | | | | |
| **ULC335-bin2** | | | | | | | | |
| **ULC335-bin3** | | | | | | | | |
| **ULC335-bin4** | | | | | | | | |
| **ULC335-nobin** | | | | | | | | |
| **ULC007-bin1** | | | | | | | | |
| **ULC007-bin2** | | | | | | | | |
| **ULC007-nobin** | | | | | | | | |
| **ULC027-bin1** | | | | | | | | |
| **ULC027-bin2** | | | | | | | | |
| **ULC027-bin3** | | | | | | | | |
| **ULC027-bin4** | | | | | | | | |
| **ULC027-nobin** | | | | | | | | |
| **ULC041-bin1** | | | | | | | | |
| **ULC041-bin2** | | | | | | | | |
| **ULC041-nobin** | | | | | | | | |
| **ULC065-bin1** | | | | | | | | |
| **ULC065-bin2** | 0.000 | | | | | | | |
| **ULC065-nobin** | 0.016 | 0.000 | | | | | | |
| **ULC066-bin1** | 0.000 | 0.000 | 0.061 | | | | | |
| **ULC066-bin2** | 0.000 | 0.000 | 0.000 | 0.000 | | | | |
| **ULC066-bin3** | 0.000 | 0.000 | 0.000 | 0.000 | 0.000 | | | |
| **ULC066-nobin** | 0.016 | 0.000 | 3.254 | 0.023 | 0.020 | 0.000 | | |
| **ULC068-bin1** | 0.000 | 0.000 | 0.017 | 0.000 | 0.000 | 0.000 | 0.000 | |
| **ULC068-bin2** | 0.000 | 0.000 | 0.000 | 0.000 | 0.000 | 0.000 | 0.000 | 0.369 |
| **ULC068-nobin** | 0.025 | 14.313 | 3.597 | 0.051 | 0.010 | 0.000 | 2.103 | 0.068 |
| **ULC073-bin1** | 0.000 | 0.000 | 0.000 | 0.000 | 0.000 | 0.000 | 0.006 | 0.000 |
| **ULC073-bin2** | 0.000 | 0.000 | 0.015 | 0.000 | 0.000 | 76.525 | 46.873 | 0.000 |
| **ULC073-bin3** | 0.000 | 0.000 | 14.180 | 0.000 | 0.000 | 0.000 | 11.449 | 0.000 |
| **ULC073-bin4** | 0.000 | 0.000 | 0.092 | 0.000 | 0.000 | 0.000 | 0.138 | 0.000 |
| **ULC073-bin5** | 0.000 | 0.000 | 0.000 | 0.000 | 0.000 | 0.801 | 2.046 | 0.000 |
| **ULC073-bin6** | 0.000 | 0.000 | 25.704 | 0.000 | 0.000 | 0.000 | 23.161 | 0.000 |
| **ULC073-nobin** | 0.024 | 0.000 | 2.161 | 0.037 | 0.000 | 1.367 | 2.653 | 0.000 |
| **ULC077-bin1** | 0.000 | 0.000 | 0.005 | 0.000 | 0.000 | 0.000 | 0.005 | 0.000 |
| **ULC077-nobin** | 0.000 | 30.895 | 2.068 | 0.037 | 0.000 | 29.283 | 9.116 | 0.017 |
| **ULC082-bin1** | 0.000 | 0.000 | 0.000 | 0.000 | 0.000 | 0.000 | 0.000 | 0.000 |
| **ULC082-bin2** | 0.000 | 0.000 | 0.022 | 0.000 | 0.000 | 0.000 | 0.000 | 0.000 |
| **ULC082-bin3** | 0.000 | 0.000 | 0.000 | 0.000 | 0.000 | 0.000 | 0.000 | 0.000 |
| **ULC082-bin4** | 0.000 | 0.000 | 0.000 | 0.000 | 0.000 | 0.000 | 0.000 | 0.000 |
| **ULC082-bin5** | 0.000 | 0.000 | 0.000 | 0.000 | 0.000 | 0.000 | 0.000 | 0.000 |
| **ULC082-nobin** | 0.000 | 0.000 | 0.025 | 0.000 | 0.000 | 0.000 | 0.006 | 0.000 |
| **ULC084-bin1** | 0.000 | 0.000 | 0.017 | 0.000 | 0.000 | 66.081 | 44.261 | 0.000 |
| **ULC084-bin2** | 0.000 | 0.000 | 7.492 | 0.000 | 0.000 | 0.000 | 0.000 | 0.000 |
| **ULC084-bin3** | 0.000 | 0.000 | 0.000 | 0.000 | 0.000 | 0.000 | 0.000 | 0.000 |
| **ULC084-nobin** | 0.000 | 0.000 | 2.723 | 0.000 | 0.000 | 5.661 | 6.817 | 0.000 |
| **ULC129-bin1** | 0.000 | 0.000 | 0.000 | 0.000 | 0.000 | 0.000 | 0.000 | 0.000 |
| **ULC129-nobin** | 0.000 | 0.010 | 0.667 | 0.000 | 0.000 | 20.400 | 8.685 | 0.000 |
| **ULC146-bin1** | 0.000 | 0.000 | 9.519 | 0.000 | 0.000 | 0.000 | 1.054 | 0.000 |
| **ULC146-bin2** | 0.000 | 0.000 | 0.000 | 0.000 | 0.000 | 0.000 | 0.000 | 0.000 |
| **ULC146-bin3** | 0.000 | 0.000 | 18.905 | 0.000 | 0.000 | 0.000 | 15.535 | 0.000 |
| **ULC146-bin4** | 0.000 | 0.000 | 0.015 | 0.000 | 0.000 | 75.549 | 46.542 | 0.000 |
| **ULC146-bin5** | 0.000 | 0.000 | 0.000 | 0.000 | 0.000 | 0.000 | 0.000 | 0.000 |
| **ULC146-bin6** | 0.000 | 0.000 | 0.083 | 0.000 | 0.000 | 0.000 | 0.028 | 0.000 |
| **ULC146-bin7** | 0.000 | 0.000 | 16.480 | 0.000 | 0.000 | 0.000 | 13.793 | 0.000 |
| **ULC146-nobin** | 0.000 | 0.000 | 1.057 | 0.000 | 0.000 | 2.001 | 1.503 | 0.000 |
| **ULC165-bin1** | 0.000 | 91.192 | 9.267 | 0.000 | 0.000 | 0.000 | 0.000 | 0.000 |
| **ULC165-bin2** | 0.000 | 0.000 | 7.601 | 0.000 | 0.000 | 0.000 | 0.000 | 0.000 |
| **ULC165-bin3** | 0.000 | 0.000 | 9.389 | 0.000 | 0.000 | 0.000 | 1.002 | 0.000 |
| **ULC165-bin4** | 0.000 | 0.000 | 0.041 | 0.000 | 0.000 | 0.000 | 0.000 | 0.000 |
| **ULC165-nobin** | 0.000 | 0.747 | 2.785 | 0.000 | 0.000 | 0.000 | 0.414 | 0.006 |
| **ULC179-bin1** | 0.000 | 0.000 | 0.000 | 0.000 | 0.000 | 0.000 | 0.000 | 0.000 |
| **ULC179-bin2** | 0.000 | 0.000 | 0.000 | 0.000 | 0.000 | 0.000 | 0.000 | 0.000 |
| **ULC179-bin3** | 0.000 | 0.000 | 0.007 | 0.000 | 0.000 | 0.000 | 0.000 | 0.000 |
| **ULC179-bin4** | 0.000 | 0.000 | 0.000 | 0.000 | 0.000 | 0.000 | 0.000 | 0.000 |
| **ULC179-bin5** | 0.000 | 0.000 | 0.000 | 0.000 | 0.000 | 0.000 | 0.000 | 0.000 |
| **ULC179-bin6** | 0.000 | 0.000 | 0.000 | 0.000 | 0.000 | 0.000 | 0.000 | 0.000 |
| **ULC179-bin7** | 0.000 | 0.000 | 0.000 | 0.000 | 0.000 | 0.000 | 0.000 | 0.000 |
| **ULC179-nobin** | 0.000 | 0.000 | 0.014 | 0.000 | 0.000 | 0.000 | 0.014 | 0.000 |
| **ULC186-bin1** | 0.000 | 0.000 | 0.000 | 0.000 | 0.000 | 0.000 | 0.000 | 0.000 |
| **ULC186-nobin** | 0.000 | 0.000 | 0.022 | 0.011 | 0.000 | 0.049 | 0.022 | 0.000 |
| **ULC187-bin1** | 0.000 | 0.000 | 0.000 | 0.000 | 0.000 | 0.000 | 0.000 | 0.000 |
| **ULC187-nobin** | 0.000 | 0.010 | 2.860 | 0.000 | 0.000 | 0.000 | 1.978 | 0.000 |

| | ULC068-bin2 | ULC068-nobin | ULC073-bin1 | ULC073-bin2 | ULC073-bin3 | ULC073-bin4 | ULC073-bin5 | ULC073-bin6 |
|---|---|---|---|---|---|---|---|---|
| **ULC335-bin1** | | | | | | | | |
| **ULC335-bin2** | | | | | | | | |
| **ULC335-bin3** | | | | | | | | |
| **ULC335-bin4** | | | | | | | | |
| **ULC335-nobin** | | | | | | | | |
| **ULC007-bin1** | | | | | | | | |
| **ULC007-bin2** | | | | | | | | |
| **ULC007-nobin** | | | | | | | | |
| **ULC027-bin1** | | | | | | | | |
| **ULC027-bin2** | | | | | | | | |
| **ULC027-bin3** | | | | | | | | |
| **ULC027-bin4** | | | | | | | | |
| **ULC027-nobin** | | | | | | | | |
| **ULC041-bin1** | | | | | | | | |
| **ULC041-bin2** | | | | | | | | |
| **ULC041-nobin** | | | | | | | | |
| **ULC065-bin1** | | | | | | | | |
| **ULC065-bin2** | | | | | | | | |
| **ULC065-nobin** | | | | | | | | |
| **ULC066-bin1** | | | | | | | | |
| **ULC066-bin2** | | | | | | | | |
| **ULC066-bin3** | | | | | | | | |
| **ULC066-nobin** | | | | | | | | |
| **ULC068-bin1** | | | | | | | | |
| **ULC068-bin2** | | | | | | | | |
| **ULC068-nobin** | 0.000 | | | | | | | |
| **ULC073-bin1** | 0.000 | 0.000 | | | | | | |
| **ULC073-bin2** | 0.000 | 0.084 | 0.000 | | | | | |
| **ULC073-bin3** | 0.000 | 0.034 | 0.000 | 0.000 | | | | |
| **ULC073-bin4** | 0.000 | 0.000 | 0.000 | 0.000 | 0.000 | | | |
| **ULC073-bin5** | 0.000 | 0.000 | 0.267 | 0.000 | 0.000 | 0.000 | | |
| **ULC073-bin6** | 0.000 | 1.181 | 0.000 | 0.000 | 0.000 | 0.000 | 0.000 | |
| **ULC073-nobin** | 0.000 | 1.385 | 0.083 | 0.038 | 0.000 | 0.000 | 0.089 | 0.000 |
| **ULC077-bin1** | 0.000 | 0.025 | 0.000 | 0.000 | 0.000 | 0.000 | 0.000 | 0.000 |
| **ULC077-nobin** | 0.000 | 4.205 | 0.000 | 37.470 | 2.328 | 0.276 | 2.046 | 5.904 |
| **ULC082-bin1** | 0.000 | 0.000 | 0.000 | 0.000 | 0.000 | 0.000 | 0.000 | 0.000 |
| **ULC082-bin2** | 0.000 | 0.011 | 0.000 | 0.000 | 0.000 | 0.000 | 0.000 | 0.000 |
| **ULC082-bin3** | 0.000 | 0.000 | 0.000 | 0.000 | 0.000 | 0.000 | 0.000 | 0.000 |
| **ULC082-bin4** | 0.000 | 0.000 | 0.000 | 0.000 | 0.000 | 0.000 | 0.000 | 0.000 |
| **ULC082-bin5** | 0.000 | 0.000 | 0.000 | 0.000 | 0.000 | 0.000 | 0.000 | 0.000 |
| **ULC082-nobin** | 0.000 | 0.017 | 0.000 | 0.000 | 0.000 | 0.000 | 0.000 | 0.000 |
| **ULC084-bin1** | 0.000 | 0.068 | 0.000 | 89.160 | 0.000 | 0.000 | 0.000 | 0.000 |
| **ULC084-bin2** | 0.000 | 2.656 | 0.000 | 0.000 | 0.000 | 0.000 | 0.000 | 0.000 |
| **ULC084-bin3** | 0.000 | 0.000 | 0.000 | 0.000 | 0.000 | 0.000 | 0.000 | 0.000 |
| **ULC084-nobin** | 0.000 | 0.135 | 0.000 | 8.430 | 17.515 | 0.000 | 3.826 | 32.243 |
| **ULC129-bin1** | 0.000 | 0.000 | 0.000 | 0.000 | 0.000 | 0.000 | 0.000 | 0.000 |
| **ULC129-nobin** | 0.000 | 1.191 | 0.000 | 25.784 | 3.609 | 1.612 | 1.246 | 8.174 |
| **ULC146-bin1** | 0.000 | 0.144 | 0.000 | 0.000 | 0.000 | 0.000 | 0.000 | 0.000 |
| **ULC146-bin2** | 0.000 | 0.000 | 0.000 | 0.000 | 0.000 | 0.000 | 0.000 | 0.000 |
| **ULC146-bin3** | 0.000 | 0.042 | 0.000 | 0.000 | 61.553 | 0.230 | 0.000 | 78.202 |
| **ULC146-bin4** | 0.000 | 0.084 | 0.000 | 92.403 | 0.000 | 0.000 | 0.000 | 0.000 |
| **ULC146-bin5** | 0.000 | 0.000 | 0.000 | 0.000 | 0.000 | 0.000 | 0.000 | 0.000 |
| **ULC146-bin6** | 0.000 | 0.028 | 0.000 | 0.000 | 0.581 | 0.092 | 0.000 | 0.000 |
| **ULC146-bin7** | 0.000 | 0.000 | 0.000 | 0.000 | 54.322 | 0.000 | 0.000 | 0.000 |
| **ULC146-nobin** | 0.000 | 1.816 | 0.000 | 1.978 | 2.667 | 11.792 | 4.270 | 11.444 |
| **ULC165-bin1** | 0.000 | 15.352 | 0.000 | 0.000 | 0.000 | 0.000 | 0.000 | 0.000 |
| **ULC165-bin2** | 0.000 | 2.706 | 0.000 | 0.000 | 0.000 | 0.000 | 0.000 | 0.000 |
| **ULC165-bin3** | 0.000 | 0.103 | 0.000 | 0.000 | 0.000 | 0.000 | 0.000 | 0.000 |
| **ULC165-bin4** | 0.000 | 0.000 | 0.000 | 0.000 | 0.000 | 0.000 | 0.000 | 0.000 |
| **ULC165-nobin** | 0.000 | 0.498 | 0.000 | 0.000 | 3.875 | 4.975 | 0.000 | 8.356 |
| **ULC179-bin1** | 0.000 | 0.000 | 0.000 | 0.000 | 0.000 | 0.000 | 0.000 | 0.000 |
| **ULC179-bin2** | 0.000 | 0.000 | 0.000 | 0.000 | 0.000 | 0.000 | 0.000 | 0.000 |
| **ULC179-bin3** | 0.000 | 0.000 | 0.000 | 0.000 | 0.000 | 0.000 | 0.000 | 0.000 |
| **ULC179-bin4** | 0.000 | 0.000 | 0.000 | 0.000 | 0.000 | 0.000 | 0.000 | 0.000 |
| **ULC179-bin5** | 0.000 | 0.000 | 0.000 | 0.000 | 0.000 | 0.000 | 0.000 | 0.000 |
| **ULC179-bin6** | 0.000 | 0.000 | 0.000 | 0.000 | 0.000 | 0.000 | 0.000 | 0.000 |
| **ULC179-bin7** | 0.000 | 0.000 | 0.000 | 0.000 | 0.000 | 0.000 | 0.000 | 0.000 |
| **ULC179-nobin** | 0.000 | 0.008 | 0.000 | 0.000 | 0.008 | 0.000 | 0.000 | 0.091 |
| **ULC186-bin1** | 0.000 | 0.000 | 0.000 | 0.000 | 0.000 | 0.000 | 0.000 | 0.000 |
| **ULC186-nobin** | 0.000 | 0.011 | 0.000 | 0.011 | 0.000 | 0.000 | 0.000 | 0.000 |
| **ULC187-bin1** | 0.123 | 0.000 | 0.000 | 0.000 | 0.000 | 0.000 | 0.000 | 0.000 |
| **ULC187-nobin** | 0.000 | 0.771 | 0.000 | 0.010 | 13.703 | 4.376 | 0.000 | 21.072 |

| | ULC073-nobin | ULC077-bin1 | ULC077-nobin | ULC082-bin1 | ULC082-bin2 | ULC082-bin3 | ULC082-bin4 | ULC082-bin5 |
|---|---|---|---|---|---|---|---|---|
| **ULC335-bin1** | | | | | | | | |
| **ULC335-bin2** | | | | | | | | |
| **ULC335-bin3** | | | | | | | | |
| **ULC335-bin4** | | | | | | | | |
| **ULC335-nobin** | | | | | | | | |
| **ULC007-bin1** | | | | | | | | |
| **ULC007-bin2** | | | | | | | | |
| **ULC007-nobin** | | | | | | | | |
| **ULC027-bin1** | | | | | | | | |
| **ULC027-bin2** | | | | | | | | |
| **ULC027-bin3** | | | | | | | | |
| **ULC027-bin4** | | | | | | | | |
| **ULC027-nobin** | | | | | | | | |
| **ULC041-bin1** | | | | | | | | |
| **ULC041-bin2** | | | | | | | | |
| **ULC041-nobin** | | | | | | | | |
| **ULC065-bin1** | | | | | | | | |
| **ULC065-bin2** | | | | | | | | |
| **ULC065-nobin** | | | | | | | | |
| **ULC066-bin1** | | | | | | | | |
| **ULC066-bin2** | | | | | | | | |
| **ULC066-bin3** | | | | | | | | |
| **ULC066-nobin** | | | | | | | | |
| **ULC068-bin1** | | | | | | | | |
| **ULC068-bin2** | | | | | | | | |
| **ULC068-nobin** | | | | | | | | |
| **ULC073-bin1** | | | | | | | | |
| **ULC073-bin2** | | | | | | | | |
| **ULC073-bin3** | | | | | | | | |
| **ULC073-bin4** | | | | | | | | |
| **ULC073-bin5** | | | | | | | | |
| **ULC073-bin6** | | | | | | | | |
| **ULC073-nobin** | | | | | | | | |
| **ULC077-bin1** | 0.000 | | | | | | | |
| **ULC077-nobin** | 1.473 | 0.970 | | | | | | |
| **ULC082-bin1** | 0.000 | 0.000 | 0.000 | | | | | |
| **ULC082-bin2** | 0.000 | 0.000 | 0.044 | 0.000 | | | | |
| **ULC082-bin3** | 0.000 | 0.000 | 0.000 | 0.000 | 0.000 | | | |
| **ULC082-bin4** | 0.000 | 0.000 | 0.187 | 0.000 | 0.000 | 0.000 | | |
| **ULC082-bin5** | 0.000 | 0.000 | 0.000 | 0.000 | 0.000 | 0.000 | 0.000 | |
| **ULC082-nobin** | 0.021 | 0.000 | 0.031 | 0.000 | 0.000 | 0.000 | 0.000 | 0.000 |
| **ULC084-bin1** | 1.286 | 0.000 | 35.431 | 0.000 | 0.000 | 0.000 | 0.000 | 0.000 |
| **ULC084-bin2** | 0.000 | 0.000 | 8.952 | 0.000 | 0.229 | 0.000 | 1.185 | 0.000 |
| **ULC084-bin3** | 0.000 | 0.000 | 0.000 | 21.352 | 0.000 | 0.000 | 0.000 | 0.000 |
| **ULC084-nobin** | 5.004 | 0.000 | 4.569 | 0.424 | 0.044 | 0.000 | 0.000 | 0.000 |
| **ULC129-bin1** | 0.000 | 0.000 | 0.000 | 0.000 | 0.000 | 0.000 | 0.000 | 0.000 |
| **ULC129-nobin** | 1.846 | 0.000 | 3.626 | 0.009 | 0.000 | 0.000 | 0.000 | 0.000 |
| **ULC146-bin1** | 8.131 | 0.000 | 0.247 | 0.000 | 0.000 | 0.000 | 0.000 | 0.000 |
| **ULC146-bin2** | 0.000 | 0.000 | 0.000 | 0.000 | 0.000 | 0.000 | 0.000 | 0.000 |
| **ULC146-bin3** | 16.232 | 0.000 | 3.169 | 0.000 | 0.000 | 0.000 | 0.000 | 0.000 |
| **ULC146-bin4** | 1.631 | 0.000 | 37.130 | 0.000 | 0.000 | 0.000 | 0.000 | 0.000 |
| **ULC146-bin5** | 0.000 | 0.000 | 0.022 | 0.000 | 0.000 | 0.000 | 0.000 | 0.000 |
| **ULC146-bin6** | 28.042 | 0.000 | 0.249 | 0.000 | 0.000 | 0.000 | 0.000 | 0.000 |
| **ULC146-bin7** | 18.451 | 0.000 | 2.732 | 0.000 | 0.000 | 0.000 | 0.000 | 0.000 |
| **ULC146-nobin** | 6.834 | 0.000 | 1.420 | 0.009 | 0.000 | 0.000 | 0.000 | 0.000 |
| **ULC165-bin1** | 0.000 | 0.000 | 31.584 | 0.000 | 0.000 | 0.000 | 0.000 | 0.000 |
| **ULC165-bin2** | 0.000 | 0.000 | 9.015 | 0.000 | 0.283 | 0.000 | 1.247 | 0.000 |
| **ULC165-bin3** | 8.490 | 0.000 | 0.257 | 0.000 | 0.000 | 0.000 | 0.000 | 0.000 |
| **ULC165-bin4** | 0.021 | 0.000 | 0.000 | 0.000 | 0.000 | 0.000 | 0.000 | 0.000 |
| **ULC165-nobin** | 2.159 | 0.000 | 0.555 | 0.009 | 0.000 | 0.000 | 0.000 | 0.000 |
| **ULC179-bin1** | 0.000 | 0.000 | 0.000 | 0.000 | 0.000 | 0.000 | 0.000 | 0.000 |
| **ULC179-bin2** | 0.000 | 0.000 | 0.000 | 0.000 | 0.000 | 0.000 | 0.000 | 0.000 |
| **ULC179-bin3** | 0.000 | 0.000 | 0.000 | 0.000 | 0.000 | 0.000 | 0.000 | 0.000 |
| **ULC179-bin4** | 0.000 | 0.000 | 0.000 | 0.000 | 0.000 | 0.000 | 0.000 | 0.000 |
| **ULC179-bin5** | 0.000 | 0.000 | 0.000 | 0.000 | 0.000 | 0.000 | 0.000 | 0.000 |
| **ULC179-bin6** | 0.000 | 0.000 | 0.000 | 0.000 | 0.000 | 0.000 | 0.000 | 0.000 |
| **ULC179-bin7** | 0.000 | 0.000 | 0.000 | 0.000 | 0.000 | 0.000 | 0.000 | 0.000 |
| **ULC179-nobin** | 0.007 | 0.000 | 0.009 | 0.000 | 0.000 | 0.000 | 0.000 | 0.000 |
| **ULC186-bin1** | 0.000 | 0.000 | 0.000 | 0.000 | 0.000 | 0.000 | 0.000 | 0.000 |
| **ULC186-nobin** | 0.032 | 0.000 | 0.022 | 0.000 | 0.000 | 0.026 | 0.000 | 0.000 |
| **ULC187-bin1** | 0.000 | 0.000 | 0.000 | 0.000 | 0.000 | 0.000 | 0.000 | 0.000 |
| **ULC187-nobin** | 6.441 | 0.000 | 0.882 | 0.010 | 0.022 | 0.000 | 0.125 | 0.000 |

| | ULC082-nobin | ULC084-bin1 | ULC084-bin2 | ULC084-bin3 | ULC084-nobin | ULC129-bin1 | ULC129-nobin | ULC146-bin1 |
|---|---|---|---|---|---|---|---|---|
| **ULC335-bin1** | | | | | | | | |
| **ULC335-bin2** | | | | | | | | |
| **ULC335-bin3** | | | | | | | | |
| **ULC335-bin4** | | | | | | | | |
| **ULC335-nobin** | | | | | | | | |
| **ULC007-bin1** | | | | | | | | |
| **ULC007-bin2** | | | | | | | | |
| **ULC007-nobin** | | | | | | | | |
| **ULC027-bin1** | | | | | | | | |
| **ULC027-bin2** | | | | | | | | |
| **ULC027-bin3** | | | | | | | | |
| **ULC027-bin4** | | | | | | | | |
| **ULC027-nobin** | | | | | | | | |
| **ULC041-bin1** | | | | | | | | |
| **ULC041-bin2** | | | | | | | | |
| **ULC041-nobin** | | | | | | | | |
| **ULC065-bin1** | | | | | | | | |
| **ULC065-bin2** | | | | | | | | |
| **ULC065-nobin** | | | | | | | | |
| **ULC066-bin1** | | | | | | | | |
| **ULC066-bin2** | | | | | | | | |
| **ULC066-bin3** | | | | | | | | |
| **ULC066-nobin** | | | | | | | | |
| **ULC068-bin1** | | | | | | | | |
| **ULC068-bin2** | | | | | | | | |
| **ULC068-nobin** | | | | | | | | |
| **ULC073-bin1** | | | | | | | | |
| **ULC073-bin2** | | | | | | | | |
| **ULC073-bin3** | | | | | | | | |
| **ULC073-bin4** | | | | | | | | |
| **ULC073-bin5** | | | | | | | | |
| **ULC073-bin6** | | | | | | | | |
| **ULC073-nobin** | | | | | | | | |
| **ULC077-bin1** | | | | | | | | |
| **ULC077-nobin** | | | | | | | | |
| **ULC082-bin1** | | | | | | | | |
| **ULC082-bin2** | | | | | | | | |
| **ULC082-bin3** | | | | | | | | |
| **ULC082-bin4** | | | | | | | | |
| **ULC082-bin5** | | | | | | | | |
| **ULC082-nobin** | | | | | | | | |
| **ULC084-bin1** | 0.000 | | | | | | | |
| **ULC084-bin2** | 0.044 | 0.000 | | | | | | |
| **ULC084-bin3** | 0.313 | 0.000 | 0.000 | | | | | |
| **ULC084-nobin** | 0.297 | 0.017 | 0.000 | 0.018 | | | | |
| **ULC129-bin1** | 0.000 | 0.000 | 0.000 | 0.000 | 0.000 | | | |
| **ULC129-nobin** | 0.063 | 24.625 | 0.009 | 0.009 | 3.356 | 0.053 | | |
| **ULC146-bin1** | 0.010 | 0.000 | 0.000 | 0.000 | 0.000 | 0.000 | 6.512 | |
| **ULC146-bin2** | 0.000 | 0.000 | 0.000 | 0.000 | 0.000 | 0.000 | 0.000 | 0.000 |
| **ULC146-bin3** | 0.000 | 0.000 | 0.000 | 0.000 | 24.100 | 0.000 | 4.124 | 0.000 |
| **ULC146-bin4** | 0.000 | 88.752 | 0.000 | 0.000 | 7.705 | 0.000 | 25.703 | 0.000 |
| **ULC146-bin5** | 0.045 | 0.000 | 0.000 | 0.000 | 0.022 | 0.000 | 0.000 | 0.000 |
| **ULC146-bin6** | 0.000 | 0.000 | 0.000 | 0.000 | 0.000 | 0.000 | 0.000 | 0.000 |
| **ULC146-bin7** | 0.000 | 0.000 | 0.000 | 0.000 | 21.093 | 0.000 | 2.956 | 0.000 |
| **ULC146-nobin** | 0.178 | 0.971 | 0.000 | 0.000 | 3.105 | 0.000 | 1.397 | 0.041 |
| **ULC165-bin1** | 0.000 | 0.000 | 0.000 | 0.000 | 0.000 | 0.000 | 0.008 | 0.000 |
| **ULC165-bin2** | 0.138 | 0.000 | 87.311 | 0.000 | 3.215 | 0.000 | 0.009 | 0.000 |
| **ULC165-bin3** | 0.000 | 0.000 | 0.000 | 0.000 | 0.000 | 0.000 | 6.358 | 84.601 |
| **ULC165-bin4** | 0.021 | 0.000 | 0.000 | 0.000 | 0.000 | 0.000 | 0.041 | 0.000 |
| **ULC165-nobin** | 0.109 | 0.000 | 0.871 | 0.009 | 1.661 | 0.005 | 1.662 | 35.053 |
| **ULC179-bin1** | 0.000 | 0.000 | 0.000 | 0.000 | 0.000 | 0.000 | 0.000 | 0.000 |
| **ULC179-bin2** | 0.000 | 0.000 | 0.000 | 0.000 | 0.000 | 0.000 | 0.000 | 0.000 |
| **ULC179-bin3** | 0.000 | 0.000 | 0.000 | 0.000 | 0.000 | 0.000 | 0.000 | 0.000 |
| **ULC179-bin4** | 0.000 | 0.000 | 0.000 | 0.000 | 0.000 | 0.000 | 0.000 | 0.000 |
| **ULC179-bin5** | 0.000 | 0.000 | 0.000 | 0.000 | 0.000 | 0.000 | 0.000 | 0.000 |
| **ULC179-bin6** | 0.000 | 0.000 | 0.000 | 0.000 | 0.000 | 0.000 | 0.000 | 0.000 |
| **ULC179-bin7** | 0.000 | 0.000 | 0.000 | 0.000 | 0.000 | 0.000 | 0.000 | 0.000 |
| **ULC179-nobin** | 0.014 | 0.000 | 0.000 | 0.000 | 0.026 | 0.000 | 0.011 | 0.005 |
| **ULC186-bin1** | 0.030 | 0.000 | 0.000 | 0.000 | 0.020 | 0.000 | 0.030 | 0.000 |
| **ULC186-nobin** | 0.194 | 0.011 | 0.011 | 0.022 | 0.183 | 0.000 | 0.172 | 0.011 |
| **ULC187-bin1** | 0.017 | 0.000 | 0.000 | 0.000 | 0.020 | 0.000 | 0.017 | 0.000 |
| **ULC187-nobin** | 0.122 | 0.010 | 8.723 | 0.020 | 3.175 | 0.000 | 1.177 | 0.000 |

| | ULC146-bin2 | ULC146-bin3 | ULC146-bin4 | ULC146-bin5 | ULC146-bin6 | ULC146-bin7 | ULC146-nobin | ULC165-bin1 |
|---|---|---|---|---|---|---|---|---|
| **ULC335-bin1** | | | | | | | | |
| **ULC335-bin2** | | | | | | | | |
| **ULC335-bin3** | | | | | | | | |
| **ULC335-bin4** | | | | | | | | |
| **ULC335-nobin** | | | | | | | | |
| **ULC007-bin1** | | | | | | | | |
| **ULC007-bin2** | | | | | | | | |
| **ULC007-nobin** | | | | | | | | |
| **ULC027-bin1** | | | | | | | | |
| **ULC027-bin2** | | | | | | | | |
| **ULC027-bin3** | | | | | | | | |
| **ULC027-bin4** | | | | | | | | |
| **ULC027-nobin** | | | | | | | | |
| **ULC041-bin1** | | | | | | | | |
| **ULC041-bin2** | | | | | | | | |
| **ULC041-nobin** | | | | | | | | |
| **ULC065-bin1** | | | | | | | | |
| **ULC065-bin2** | | | | | | | | |
| **ULC065-nobin** | | | | | | | | |
| **ULC066-bin1** | | | | | | | | |
| **ULC066-bin2** | | | | | | | | |
| **ULC066-bin3** | | | | | | | | |
| **ULC066-nobin** | | | | | | | | |
| **ULC068-bin1** | | | | | | | | |
| **ULC068-bin2** | | | | | | | | |
| **ULC068-nobin** | | | | | | | | |
| **ULC073-bin1** | | | | | | | | |
| **ULC073-bin2** | | | | | | | | |
| **ULC073-bin3** | | | | | | | | |
| **ULC073-bin4** | | | | | | | | |
| **ULC073-bin5** | | | | | | | | |
| **ULC073-bin6** | | | | | | | | |
| **ULC073-nobin** | | | | | | | | |
| **ULC077-bin1** | | | | | | | | |
| **ULC077-nobin** | | | | | | | | |
| **ULC082-bin1** | | | | | | | | |
| **ULC082-bin2** | | | | | | | | |
| **ULC082-bin3** | | | | | | | | |
| **ULC082-bin4** | | | | | | | | |
| **ULC082-bin5** | | | | | | | | |
| **ULC082-nobin** | | | | | | | | |
| **ULC084-bin1** | | | | | | | | |
| **ULC084-bin2** | | | | | | | | |
| **ULC084-bin3** | | | | | | | | |
| **ULC084-nobin** | | | | | | | | |
| **ULC129-bin1** | | | | | | | | |
| **ULC129-nobin** | | | | | | | | |
| **ULC146-bin1** | | | | | | | | |
| **ULC146-bin2** | | | | | | | | |
| **ULC146-bin3** | 0.000 | | | | | | | |
| **ULC146-bin4** | 0.000 | 0.000 | | | | | | |
| **ULC146-bin5** | 0.000 | 0.000 | 0.000 | | | | | |
| **ULC146-bin6** | 0.000 | 0.000 | 0.000 | 0.000 | | | | |
| **ULC146-bin7** | 0.000 | 0.000 | 0.000 | 0.000 | 0.000 | | | |
| **ULC146-nobin** | 0.006 | 0.007 | 0.077 | 0.000 | 0.000 | 0.000 | | |
| **ULC165-bin1** | 0.000 | 0.000 | 0.000 | 0.000 | 0.000 | 0.000 | 0.000 | |
| **ULC165-bin2** | 0.000 | 0.000 | 0.000 | 0.000 | 0.000 | 0.000 | 0.000 | 0.000 |
| **ULC165-bin3** | 0.000 | 0.000 | 0.000 | 0.000 | 0.000 | 0.000 | 0.437 | 0.000 |
| **ULC165-bin4** | 0.000 | 0.000 | 0.000 | 0.000 | 0.000 | 0.000 | 0.267 | 0.000 |
| **ULC165-nobin** | 0.006 | 4.124 | 0.000 | 0.168 | 4.342 | 3.269 | 1.334 | 0.016 |
| **ULC179-bin1** | 0.000 | 0.000 | 0.000 | 0.000 | 0.000 | 0.000 | 0.000 | 0.000 |
| **ULC179-bin2** | 0.000 | 0.000 | 0.000 | 0.000 | 0.000 | 0.000 | 6.177 | 0.000 |
| **ULC179-bin3** | 0.000 | 0.000 | 0.000 | 0.000 | 0.000 | 0.000 | 0.007 | 0.000 |
| **ULC179-bin4** | 0.000 | 0.000 | 0.000 | 0.000 | 0.000 | 0.000 | 0.000 | 0.000 |
| **ULC179-bin5** | 0.000 | 0.000 | 0.000 | 0.000 | 0.000 | 0.000 | 0.172 | 0.000 |
| **ULC179-bin6** | 0.000 | 0.000 | 0.000 | 0.000 | 0.000 | 0.000 | 2.205 | 0.000 |
| **ULC179-bin7** | 0.000 | 0.000 | 0.000 | 0.000 | 0.000 | 0.000 | 0.053 | 0.000 |
| **ULC179-nobin** | 0.006 | 0.022 | 0.000 | 0.000 | 0.000 | 0.000 | 0.735 | 0.000 |
| **ULC186-bin1** | 0.000 | 0.000 | 0.000 | 0.000 | 0.000 | 0.000 | 0.015 | 0.000 |
| **ULC186-nobin** | 0.000 | 0.000 | 0.000 | 0.000 | 0.000 | 0.000 | 0.205 | 0.000 |
| **ULC187-bin1** | 0.000 | 0.000 | 0.000 | 0.000 | 0.000 | 0.000 | 0.017 | 0.000 |
| **ULC187-nobin** | 0.010 | 19.860 | 0.010 | 0.011 | 0.332 | 11.464 | 2.475 | 0.010 |

| | ULC165-bin2 | ULC165-bin3 | ULC165-bin4 | ULC165-nobin | ULC179-bin1 | ULC179-bin2 | ULC179-bin3 | ULC179-bin4 |
|---|---|---|---|---|---|---|---|---|
| **ULC335-bin1** | | | | | | | | |
| **ULC335-bin2** | | | | | | | | |
| **ULC335-bin3** | | | | | | | | |
| **ULC335-bin4** | | | | | | | | |
| **ULC335-nobin** | | | | | | | | |
| **ULC007-bin1** | | | | | | | | |
| **ULC007-bin2** | | | | | | | | |
| **ULC007-nobin** | | | | | | | | |
| **ULC027-bin1** | | | | | | | | |
| **ULC027-bin2** | | | | | | | | |
| **ULC027-bin3** | | | | | | | | |
| **ULC027-bin4** | | | | | | | | |
| **ULC027-nobin** | | | | | | | | |
| **ULC041-bin1** | | | | | | | | |
| **ULC041-bin2** | | | | | | | | |
| **ULC041-nobin** | | | | | | | | |
| **ULC065-bin1** | | | | | | | | |
| **ULC065-bin2** | | | | | | | | |
| **ULC065-nobin** | | | | | | | | |
| **ULC066-bin1** | | | | | | | | |
| **ULC066-bin2** | | | | | | | | |
| **ULC066-bin3** | | | | | | | | |
| **ULC066-nobin** | | | | | | | | |
| **ULC068-bin1** | | | | | | | | |
| **ULC068-bin2** | | | | | | | | |
| **ULC068-nobin** | | | | | | | | |
| **ULC073-bin1** | | | | | | | | |
| **ULC073-bin2** | | | | | | | | |
| **ULC073-bin3** | | | | | | | | |
| **ULC073-bin4** | | | | | | | | |
| **ULC073-bin5** | | | | | | | | |
| **ULC073-bin6** | | | | | | | | |
| **ULC073-nobin** | | | | | | | | |
| **ULC077-bin1** | | | | | | | | |
| **ULC077-nobin** | | | | | | | | |
| **ULC082-bin1** | | | | | | | | |
| **ULC082-bin2** | | | | | | | | |
| **ULC082-bin3** | | | | | | | | |
| **ULC082-bin4** | | | | | | | | |
| **ULC082-bin5** | | | | | | | | |
| **ULC082-nobin** | | | | | | | | |
| **ULC084-bin1** | | | | | | | | |
| **ULC084-bin2** | | | | | | | | |
| **ULC084-bin3** | | | | | | | | |
| **ULC084-nobin** | | | | | | | | |
| **ULC129-bin1** | | | | | | | | |
| **ULC129-nobin** | | | | | | | | |
| **ULC146-bin1** | | | | | | | | |
| **ULC146-bin2** | | | | | | | | |
| **ULC146-bin3** | | | | | | | | |
| **ULC146-bin4** | | | | | | | | |
| **ULC146-bin5** | | | | | | | | |
| **ULC146-bin6** | | | | | | | | |
| **ULC146-bin7** | | | | | | | | |
| **ULC146-nobin** | | | | | | | | |
| **ULC165-bin1** | | | | | | | | |
| **ULC165-bin2** | | | | | | | | |
| **ULC165-bin3** | 0.000 | | | | | | | |
| **ULC165-bin4** | 0.000 | 0.000 | | | | | | |
| **ULC165-nobin** | 0.009 | 0.000 | 0.000 | | | | | |
| **ULC179-bin1** | 0.000 | 0.000 | 0.000 | 0.000 | | | | |
| **ULC179-bin2** | 0.000 | 0.000 | 0.000 | 0.000 | 0.000 | | | |
| **ULC179-bin3** | 0.000 | 0.000 | 0.000 | 0.000 | 0.000 | 0.000 | | |
| **ULC179-bin4** | 0.000 | 0.000 | 0.000 | 0.000 | 0.000 | 0.000 | 0.000 | |
| **ULC179-bin5** | 0.000 | 0.000 | 0.000 | 0.000 | 0.057 | 0.000 | 0.000 | 0.000 |
| **ULC179-bin6** | 0.000 | 0.000 | 0.000 | 0.000 | 0.000 | 0.000 | 0.000 | 0.000 |
| **ULC179-bin7** | 0.000 | 0.000 | 0.000 | 0.000 | 0.000 | 0.000 | 0.000 | 0.000 |
| **ULC179-nobin** | 0.000 | 0.000 | 0.000 | 0.021 | 0.005 | 0.005 | 0.033 | 0.000 |
| **ULC186-bin1** | 0.000 | 0.000 | 0.000 | 0.020 | 0.000 | 0.000 | 0.000 | 0.000 |
| **ULC186-nobin** | 0.011 | 0.000 | 0.000 | 0.259 | 0.000 | 0.000 | 0.000 | 0.000 |
| **ULC187-bin1** | 0.000 | 0.000 | 0.000 | 0.011 | 0.000 | 0.000 | 0.000 | 0.000 |
| **ULC187-nobin** | 9.058 | 0.000 | 0.000 | 0.700 | 0.000 | 0.000 | 0.000 | 0.000 |

| | ULC179-bin5 | ULC179-bin6 | ULC179-bin7 | ULC179-nobin | ULC186-bin1 | ULC186-nobin | ULC187-bin1 | ULC187-nobin |
|---|---|---|---|---|---|---|---|---|
| **ULC335-bin1** | | | | | | | | |
| **ULC335-bin2** | | | | | | | | |
| **ULC335-bin3** | | | | | | | | |
| **ULC335-bin4** | | | | | | | | |
| **ULC335-nobin** | | | | | | | | |
| **ULC007-bin1** | | | | | | | | |
| **ULC007-bin2** | | | | | | | | |
| **ULC007-nobin** | | | | | | | | |
| **ULC027-bin1** | | | | | | | | |
| **ULC027-bin2** | | | | | | | | |
| **ULC027-bin3** | | | | | | | | |
| **ULC027-bin4** | | | | | | | | |
| **ULC027-nobin** | | | | | | | | |
| **ULC041-bin1** | | | | | | | | |
| **ULC041-bin2** | | | | | | | | |
| **ULC041-nobin** | | | | | | | | |
| **ULC065-bin1** | | | | | | | | |
| **ULC065-bin2** | | | | | | | | |
| **ULC065-nobin** | | | | | | | | |
| **ULC066-bin1** | | | | | | | | |
| **ULC066-bin2** | | | | | | | | |
| **ULC066-bin3** | | | | | | | | |
| **ULC066-nobin** | | | | | | | | |
| **ULC068-bin1** | | | | | | | | |
| **ULC068-bin2** | | | | | | | | |
| **ULC068-nobin** | | | | | | | | |
| **ULC073-bin1** | | | | | | | | |
| **ULC073-bin2** | | | | | | | | |
| **ULC073-bin3** | | | | | | | | |
| **ULC073-bin4** | | | | | | | | |
| **ULC073-bin5** | | | | | | | | |
| **ULC073-bin6** | | | | | | | | |
| **ULC073-nobin** | | | | | | | | |
| **ULC077-bin1** | | | | | | | | |
| **ULC077-nobin** | | | | | | | | |
| **ULC082-bin1** | | | | | | | | |
| **ULC082-bin2** | | | | | | | | |
| **ULC082-bin3** | | | | | | | | |
| **ULC082-bin4** | | | | | | | | |
| **ULC082-bin5** | | | | | | | | |
| **ULC082-nobin** | | | | | | | | |
| **ULC084-bin1** | | | | | | | | |
| **ULC084-bin2** | | | | | | | | |
| **ULC084-bin3** | | | | | | | | |
| **ULC084-nobin** | | | | | | | | |
| **ULC129-bin1** | | | | | | | | |
| **ULC129-nobin** | | | | | | | | |
| **ULC146-bin1** | | | | | | | | |
| **ULC146-bin2** | | | | | | | | |
| **ULC146-bin3** | | | | | | | | |
| **ULC146-bin4** | | | | | | | | |
| **ULC146-bin5** | | | | | | | | |
| **ULC146-bin6** | | | | | | | | |
| **ULC146-bin7** | | | | | | | | |
| **ULC146-nobin** | | | | | | | | |
| **ULC165-bin1** | | | | | | | | |
| **ULC165-bin2** | | | | | | | | |
| **ULC165-bin3** | | | | | | | | |
| **ULC165-bin4** | | | | | | | | |
| **ULC165-nobin** | | | | | | | | |
| **ULC179-bin1** | | | | | | | | |
| **ULC179-bin2** | | | | | | | | |
| **ULC179-bin3** | | | | | | | | |
| **ULC179-bin4** | | | | | | | | |
| **ULC179-bin5** | | | | | | | | |
| **ULC179-bin6** | 0.000 | | | | | | | |
| **ULC179-bin7** | 0.000 | 0.000 | | | | | | |
| **ULC179-nobin** | 0.000 | 0.000 | 0.000 | | | | | |
| **ULC186-bin1** | 0.000 | 0.000 | 0.000 | 0.000 | | | | |
| **ULC186-nobin** | 0.000 | 0.000 | 0.000 | 0.022 | 0.269 | | | |
| **ULC187-bin1** | 0.000 | 0.000 | 0.000 | 0.000 | 0.000 | 0.043 | | |
| **ULC187-nobin** | 0.000 | 0.000 | 0.000 | 0.030 | 0.101 | 0.151 | 0.091 | |
